# SiR-XActin: A fluorescent probe for imaging actin dynamics in live cells

**DOI:** 10.1101/2025.02.04.636537

**Authors:** Veselin Nasufovic, Julian Kompa, Halli L. Lindamood, Merle Blümke, Birgit Koch, Victoria Le-vario-Diaz, Katharina Weber, Marlene Maager, Elisabetta Ada Cavalcanti-Adam, Eric A. Vitriol, Hans-Dieter Arndt, Kai Johnsson

## Abstract

Imaging actin-dependent processes in live cells is important for understanding numerous biological processes. However, currently used natural-product based fluorescent probes for actin filaments affect the dynamics of actin polymerization and can induce undesired cellular phenotypes. Here, we introduce SiR-XActin, a simplified jasplakinolide-based, far-red fluorescent probe that enables bright and photostable staining in various cell types without requiring genetic modifications. Due to its relatively weak binding affinity, the probe exhibits minimal cytotoxicity and labels actin filaments without significantly altering actin dynamics. Furthermore, SiR-XActin is suitable for time-resolved, live-cell super-resolution STED microscopy. Exchanging the SiR fluorophore in SiR-XActin for other fluorophores yields probes in different colors. All these properties make SiR-XActin and its analogs powerful tools for studying actin dynamics using live-cell fluorescence microscopy.

## Introduction

The actin network is an essential part of the eukaryotic cytoskeleton and has both structural roles and dynamic functions, for example in cytokinesis, cell-shape adaptation, cell division, and organelle trafficking.^[1]^ Actin is one of the most abundant proteins in eukaryotic cells with a high level of evolutionary sequence and structure preservation.^[2]^ Globular actin (G-actin, 42 kDa) polymerizes in a highly regulated process to form fibres (F-actin or microfilaments).^[3-4]^ Numerous actin-binding proteins (ABPs) interact with G-actin or F-actin to regulate its functions (**Figure 1A**).^[5]^ F-actin is often organized in stable networks (e.g. myofibrils, axon rings) or dynamic higher-order organizational networks (e.g. filopodia, lamellipodia, actin cortex, contractile rings, focal adhesions, stress fibres).^[6]^

**Figure 1.**
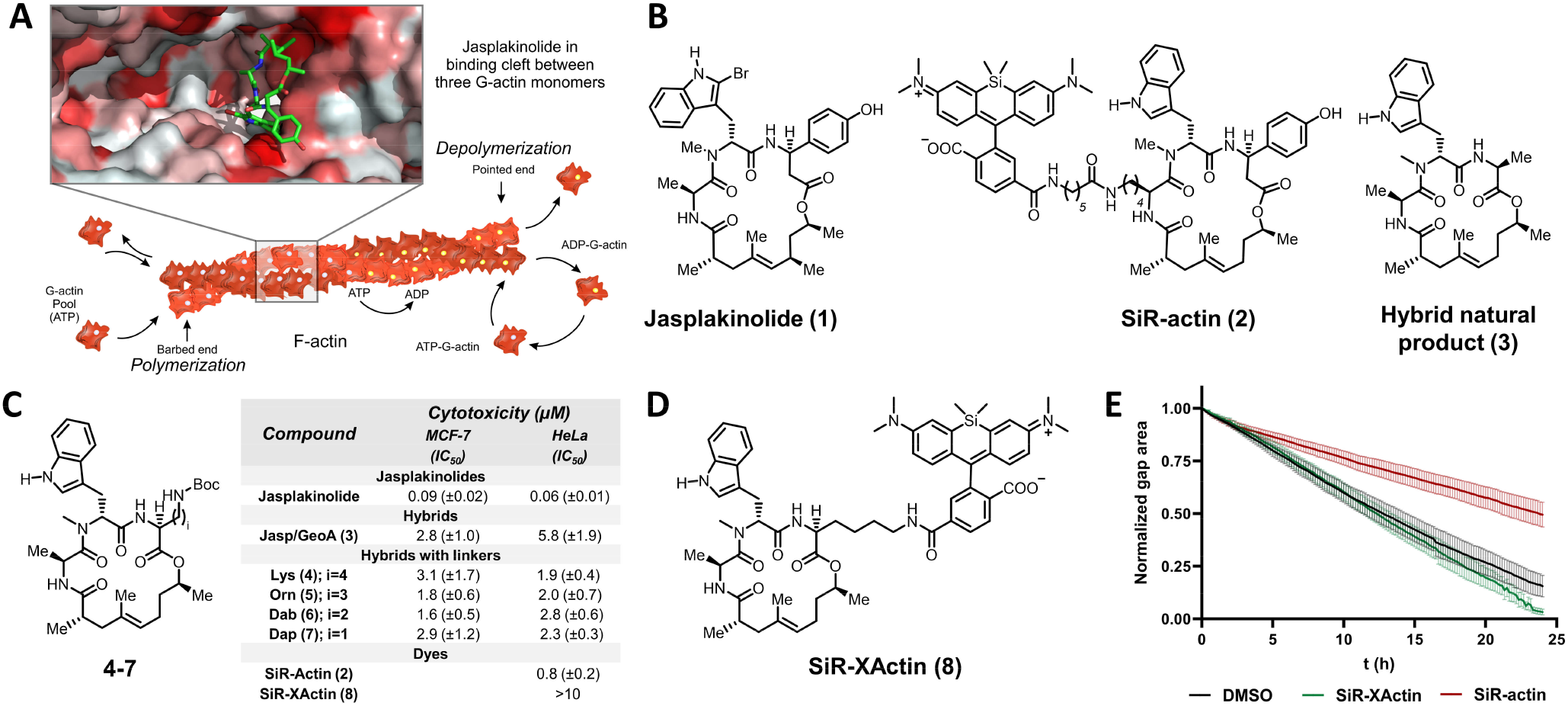
SiR-XActin is a hybrid natural product with reduced cytotoxicity and perturbation of cellular dynamics: **(A)** Scheme of actin polymerization/depolymerization machinery and binding cleft of jasplakinolide-like natural products. Cryo-EM structure of jasplakinolide-stabilized F-actin (PDB: 6T24) was used for generating cartoon of jasplakinolide binding to F-actin. **(B)** Structures of natural product jasplakinolide (**1**), cell permeable F-actin labeling tool SiR-actin (**2**), non-toxic hybrid natural product (**3**). **(C)** Synthesized analogs and cytotoxicity profiling. **(D)** Structure of SiR-XActin (**8**). **(E)** Comparison of SiR-actin and SiR-XActin influence on migration of HeLa cells in wound healing assay, 500 nM dye concentration used. Shown curves represent normalized wound gap area over time. Average values and standard deviation of five independent measurements.

An important advance in the visualization of F-actin in fixed and permeabilized cells was the development of fluorescent derivatives of the F-actin-binding natural product phalloidin.^[7]^ Phalloidin binds at the interface of three actin monomers in F-actin, increasing the stability of the fiber and slowing the dynamic actin polymerization/depolymerization process.^[8-11]^ In 2014, derivatives of the natural product jasplakinolide, which competes in binding with phalloidin, were conjugated to the far-red fluorogenic dye Si-rhodamine (SiR), resulting in the cell-permeable SiR-actin.^[12-14]^ This, enabled visualization of F-actin in living cells, such as immortalized cell lines, primary cells, organoids, and tissues.^[15-16]^ Sub-sequently, jasplakinolide derivatives with different fluorophores were introduced to complement SiR-actin.^[17-19]^ However, jasplakinolide and its fluorescent derivatives stabilize F-actin and can affect the dynamics of actin polymerization/depolymerization.^[20-22]^ More recently, another fluorescent probe, SPY650-FastAct, was introduced for imaging of dynamic actin populations, however, its structure wasn’t revealed.^[23]^

Bacterial actin-binding peptides (LifeAct),^[24]^ actin-binding proteins^[25]^, fluorescent protein fusions with G-actin^[26]^ and de-novo engineered small proteins^[27-28]^ as fused to fluorescent- or to self-labeling proteins are commonly used as genetically encoded alternatives for imaging F-actin. In particular, among these, LifeAct has become a popular tool for imaging F-actin, due to its relatively small effect on the dynamics of actin (de)polymerization, provided its expression levels are strictly controlled.^[20-21]^ However, LifeAct requires the genetic modification of cells, making small molecule probes the preferred- and often the only option for many applications.

Here we introduce new fluorescent simplified jasplakinolide derivatives that distinguish themselves by their reduced affinity for F-actin and associated low cytotoxicity. These probes enable the imaging of highly dynamic F-actin populations in live cells.

## Results and discussion

We have recently shown that a hybrid of the natural products jasplakinolide and geodiamolide A (**3, Figure 1B**) exhibits only weak cytotoxicity in comparison to parental jasplakinolide **1**.^[29]^ Based on this finding, we reasoned that this ligand would be an excellent starting point for the development of imaging probes with reduced toxicity. To convert **3** into a scaffold for fluorescent F-actin probes, we synthesized a series of jasplakinolide derivatives with different linker structures (**Figure 1C**). These analogs showed similar low cytotoxicity as **3** and we chose analog **4** (named XActin) bearing a lysine linker for attachment of fluorescent probes. Cryo-EM structures of jasplakinolide and its analogs binding to F-actin, as well as docking of possible conjugates, indicate that this lysine residue is pointing outside the binding cleft and should allow the attachment of fluorophores without disturbing the binding significantly (**Figure 1A, Figure SI1**).^[30-32]^ Attaching Si-rhodamine (SiR) to XActin yielded the far-red fluorescent conjugate SiR-XActin (**Figure 1D**). SiR-XActin showed no significant cytotoxicity in HeLa cells at all concentrations tested (IC_50_ > 10 µM), whereas SiR-actin showed sub-micromolar toxicity under these conditions (**Figure 1D, Figure SI2**). Since inhibition of actin dynamics directly effects cell motility,^[33]^ we compared the behavior of SiR-actin and SiR-XActin in a wound healing assay. Whereas SiR-XActin showed no significant influence on the migration of HeLa cells and gap closure, SiR-actin significantly slowed down cell motility, preventing wound fronts from closing the gap (**Figure 1E, Figure SI3**).

When incubated with fixed cells, SiR-XActin showed a typical F-actin staining profile by confocal laser-scanning microscopy (CLSM), comparable with SiR-actin (**Figure 2A-*i*** and **Figure SI4**). However, the specific signal from SiR-XActin-staining could be quickly washed out, contrary to the signal of SiR-actin, which remains stable even after washing (**Figure 2B-*ii***). Jasplakinolide outcompetes the binding of SiR-XActin in fixed cells completely, contrary to SiR-actin, where the fluorescent signal of F-actin could be still detected in the presence of jasplakinolide, indicating again lower affinity of the new probe compared to SiR-actin (**Figure SI4**). We then measured the apparent affinities of SiR-XActin and SiR-actin for F-actin in a microscopy-based assay. SiR-actin showed a Kd_app_ of 6.0 (±0.3) nM, which is in agreement with a previously reported value,^[34]^ whereas SiR-XActin showed an approximately 100-fold lower affinity with a Kd_app_ of 557 (±266) nM. These differences in binding affinities (**Figure SI5**) provide a rationale for the striking differences in cytotoxicity as well as in cell motility.

**Figure 2.**
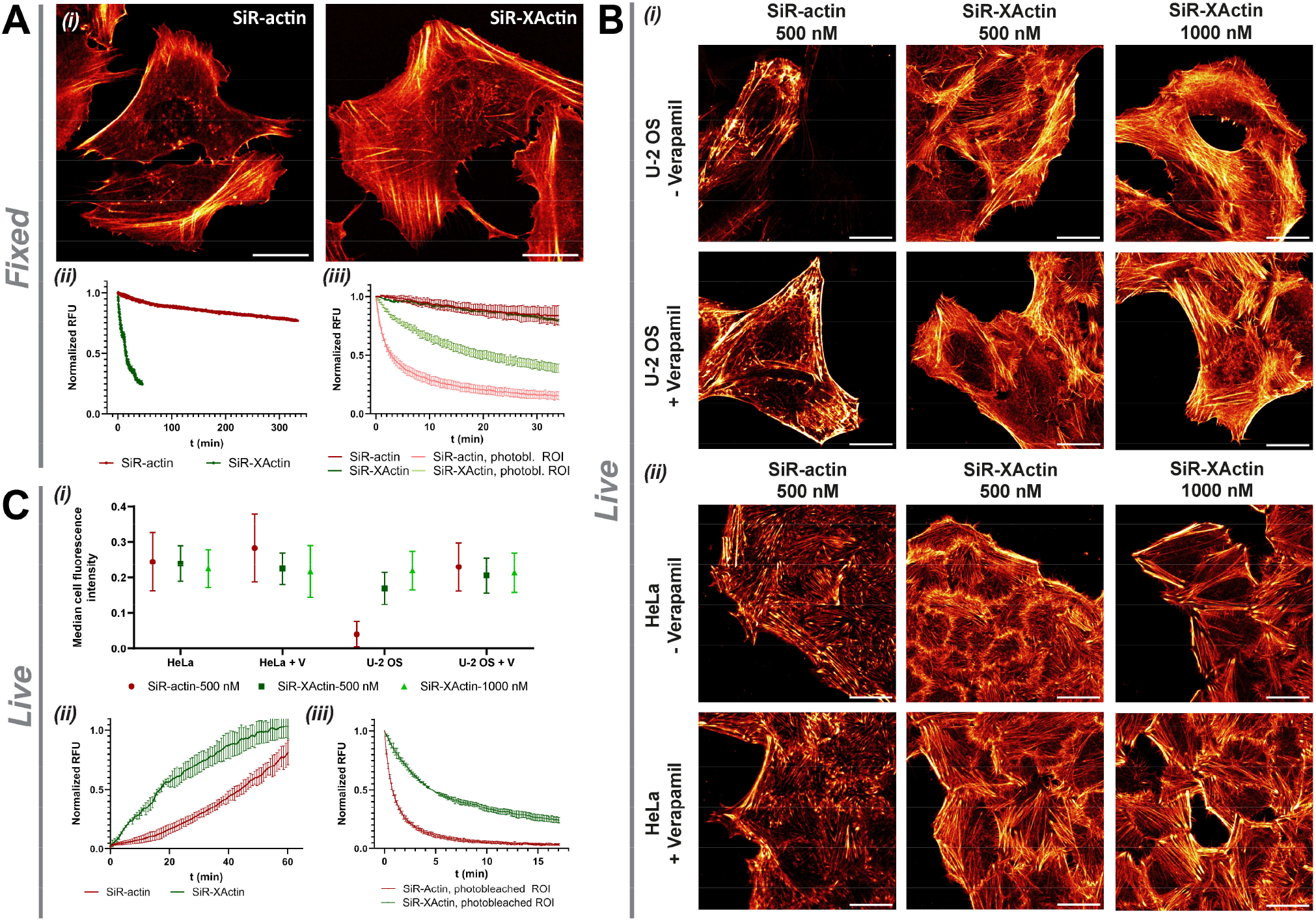
Labeling performance of SiR-XActin in fixed and live cells: **(A)** *i*: Fixed-cell no-wash CLSM images of U-2 OS cells. 652 nm excitation with 0.4% (SiR-actin) or 2% (SiR-XActin) laser power. *ii*: Fluorescence signal reduction in fixed U-2 OS cells labelled with 500 nM dye after medium exchange. The signal was normalized to the no-wash fluorescence intensity of each replicate. *iii*: Fluorescence intensity bleaching curve of SiR-actin and SiR-XActin (500 nM) in U-2 OS cells. 652 nm excitation (0.1% laser power) used for imaging, 652 nm (100% laser power) used for bleaching. The signal was normalized to the initial fluorescence intensity of each measurement. **(B)** *i*: U-2 OS and *ii*: HeLa cells stained with SiR-actin and SiR-XActin at the given dye concentrations in presence and absence of verapamil (10 µM), 4 h of incubation. **(C)** *i*: Median fluorescence signal in U-2 OS and HeLa cells (condition described in B-*i* and B*-ii*, V - verapamil). Average values and standard deviation from three independent measurements. *ii*: Live-cell labeling kinetics of SiR-actin and SiR-XActin (both 500 nM) in HeLa cells, signal was normalised to signal intensity measured after 4 h of incubation. Average values and standard deviation from three biological replicates / independent measurements. *iii*: Live-cell bleaching curves for SiR-actin and SiR-XActin (both 500 nM) in HeLa cells, incubation time 4 h, 652 nm excitation (0.1% laser power) used for imaging, 652 nm (100% laser power) used for bleaching. Red Hot lookup - pixel intensities were scaled between 0 (black) and 65535 (bright yellow) by Fiji. Scale bars 20 µm.

It has been previously demonstrated that weak-affinity fluorescent probes provide higher photostability, as the low affinity facilitates the exchange of bleached probes.^[35]^ Both SiR-actin and SiR-XActin show excellent photostability in regular imaging mode when staining fixed actin filaments (650 nm excitation, 0.1% laser power), however, in photobleaching mode (650 nm excitation, 100% laser power) SiR-XActin shows significantly slower signal decay (**Figure 2A-*iii***). At the same concentration (500 nM) SiR-actin shows a 1.8-fold higher fluorescent signal when staining fixed actin filaments than SiR-XActin (**Figure SI5**), whereas at concentrations above the respective Kd_app_, SiR-actin and SiR-XActin show almost identical brightness.

Next, we tested the ability of SiR-XActin to label F-actin populations in live cells (**Figure 2B** and **2C**). SiR-XActin labels efficiently F-actin in U-2 OS and HeLA cells, with and without the use of the efflux pump inhibitor verapamil. The staining profile of SiR-XActin in live cells is similar to the stained F-actin networks in fixed cells (**Figure 2A-*i***). However, SiR-actin showed a higher preference for stress fibers and for structures around cell edges (**Figure 2B-*i***). Additionally, SiR-actin staining in live cells resulted in bright condensates along the fibers, which were absent in fixed samples stained either by SiR-Actin or SiR-XActin as well as absent in live cells stained by SiR-XActin. These observations suggest that under these conditions SiR-actin is inducing non-physiological actin phenotypes. SiR-actin requires the presence of verapamil for efficient labeling of F-actin in U-2 OS cells, whereas SiR-XActin shows efficient labeling even in the absence of verapamil (**Figure 2B-*i* and 2C-*i***). Efficient labeling in the absence of verapamil was demonstrated for SiR-XActin in two additional cell lines (**Figure SI6**).

When HeLa cells were incubated with a range of concentrations of SiR-XActin and SiR-actin (0.04 µM-10 µM) for 6 h, only SiR-actin induced condensation of F-actin, similar to that previously reported for other jasplakinolide derivatives.^[36]^ SiR-actin induced the formation of brightly labelled aggregates in concentration ranges of 0.1-10 µM (**Figure SI7A**). In contrast, such aggregation was not observed with SiR-XActin, regardless of the concentration used (**Figure SI7A**). SiR-actin-treated (0.1-10 µM) HeLa cells showed membrane blebbing **(Figure SI7A)**, a phenotype that also has been reported previously for jasplakinolide itself.^[37]^ Membrane blebbing was not observed with SiR-XActin even at high concentrations (10 µM) **(Figure SI8**). In U-2 OS cells, after 16 h incubation time, neither SiR-actin nor SiR-XActin induces blebbing (**Figure SI9**), but SiR-actin staining results in bright condensates along stress fibers, even at lower concentrations (**Figure SI7B, Figure 2B-*i***). The above comparisons were done at concentrations of both probes that are 10-fold above the Kd_app_ of SiR-actin. We therefore also compared staining with SiR-actin at concentrations of 30 nM and SiR-XActin at 500 nM. After 2 h incubations, SiR-XActin reached labeling profile which did not change over time (over 8 h). In contrast, SiR-actin after 2 h showed only weak labeling in comparison, although its preference for stress fibers and focal adhesions can be seen (**Figure SI10**).

When the kinetics of labeling were recorded in live HeLa cells, significant differences were observed between SiR-XActin and SiR-actin (**Figure 2C-*ii***). SiR-XActin showed faster labeling (t_1/2_ ∼ 20 min) than SiR-actin (t_1/2_ ∼ 111 min), which also displayed a lag phase, suggesting that SiR-XActin possesses a higher cell permeability than SiR-actin (**Figure SI11, Video SI1**-**2**). As observed in fixed cells (**Figure 2B-*iii***), SiR-XActin also displayed higher photostability than SiR-actin in live cells (**Figure 2C-*iii***). The low cytotoxicity, high brightness and photostability make SiR-XActin attractive for long-term time-lapse imaging experiments. To demonstrate this, we incubated live HeLa cells with SiR-XActin at a concentration above its Kd_app_ (1 µM) for 24 h, followed by imaging every 30 s. The signal intensity of SiR-XActin stayed stable over 35 h and no obvious influence on the actin phenotype was observed, i.e. cells moved freely and divided. Multiple lamellipodiae, filopodiae and contractile rings can be seen during the time-lapse, indicating once again very low perturbance of this highly regulated network (**Figure SI12 and SI13, Video SI3 and SI4**). To demonstrate the usability of SiR-XActin, we performed dual-color CLSM imaging together with the focal adhesion marker mCherry-Vinculin (**Figure SI14A**) and mEmerald-Paxilin (**Figure SI14B**).^[38]^

SiR-actin has been very useful for application in live-cell stimulated emission depletion (STED) nanoscopy,[^14, 16^] and we therefore explored the potential of SiR-XActin for such applications (**Figure 3**). STED imaging of single HeLa cells reveals a higher abundance of actin stress fibers of SiR-actin in comparison to SiR-XActin staining (**Figure 3A**). The images obtained for SiR-XActin staining are similar to STED imaging of the genetically encoded LifeAct-HaloTag7 with a SiR-HaloTag Ligand (**Figure SI15**). Also, labeling with SiR-actin increases the thickness of stress fibers in the center of the cells (**Figure 3B-C**) and the appearance of filopodia on the cell’s edges is reduced (**Figure 3B**). SiR-XActin showed again higher photostability in time-lapsed STED imaging (**Figure 3D, Video SI5-7**) compared to SiR-actin. After an initial bleaching phase, SiR-XActin shows a stable signal of around 30±3% of its initial intensity. We postulate that this difference is mainly due to the lower binding affinity of SiR-XActin, which should facilitate the exchange of bleached probe, as demonstrated for other weak-affinity labels.^[39-40]^ SiR-XActin can be used in combination with other (exchangeable) labeling tools for STED imaging. We used SiR-XActin to label the F-actin network and either exchangeable HaloTagLig-ands (JF_585_-S5, Cox8A-HaloTag7)^[40]^ (**Figure 3E, Figure SI16A**) or PK Mito Orange (PKMO)^[41]^ (**Figure SI16B**) to label mitochondria in live hippocampal neurons. Both probes can be imaged with the same depletion laser and are compatible with 3D-STED imaging (**Figure SI16C, Video SI6**).

**Figure 3.**
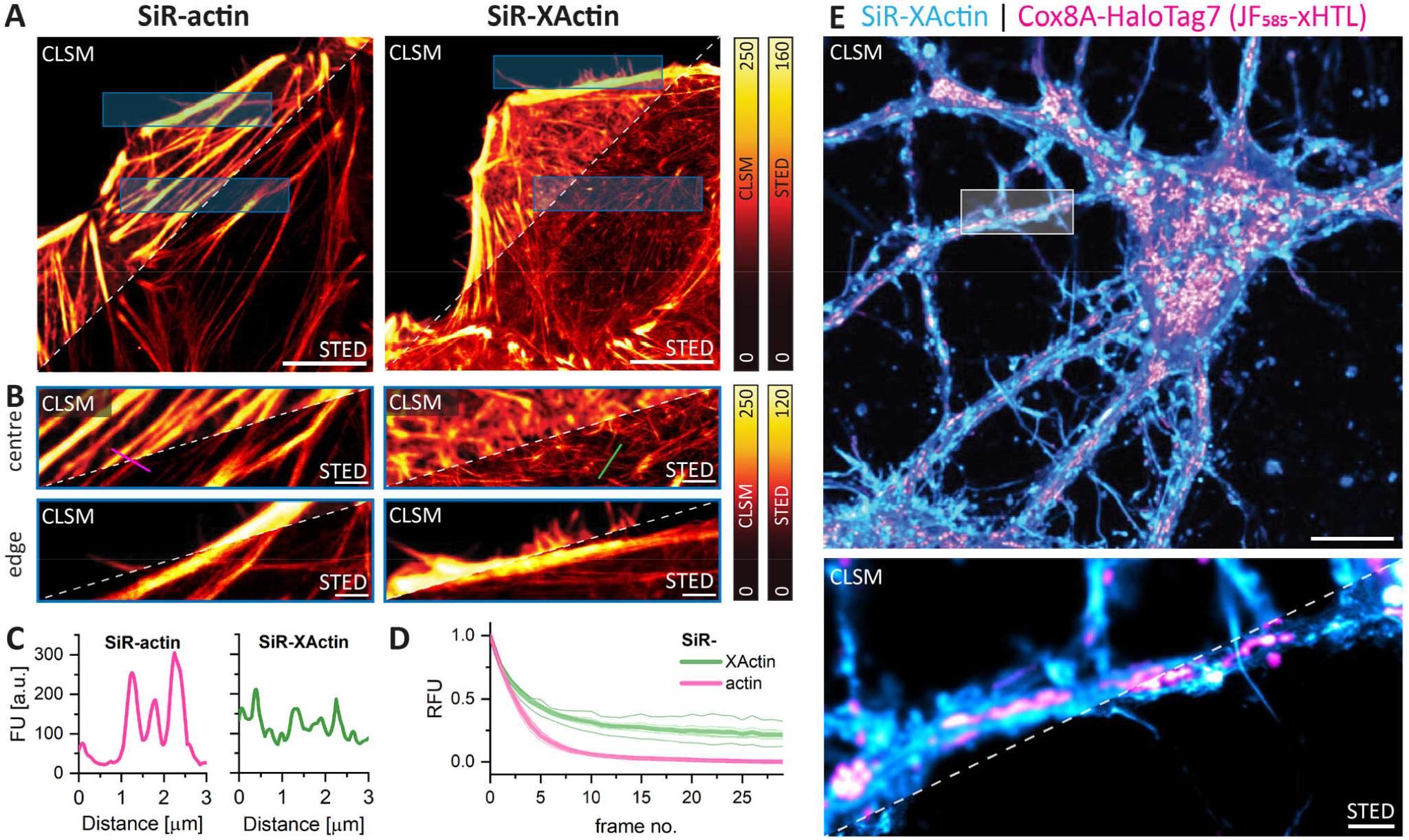
SiR-XActin is a tool to study F-actin in live cells by STED microscopy: **(A)** Phenotypic comparison of SiR-actin and SiR-XActin stainings of live HeLa cells. Cells were stained with 1 μM actin probes for 2 h, no-wash imaging. Representative CLSM and STED images. Pixel intensities were scaled between dark and bright colors according to the reference bars. Scale bars: 10 μm. **(B)** Magnification of blue ROIs from A. Scale bars: 1 μm. **(C)** Line-scan profiles shown in B. (STED) perpendicular to actin filaments. Staining with SiR-actin reveals increased number and thickness of stress fibres in the centre and a fewer filopodia on the cell’s edges in comparison to SiR-XActin staining. **(D)** Normalized fluorescence intensity of multi-frame STED imaging in a 10x10 μm ROI. U-2 OS cells were stained with 1 μM SiR-actin or SiR-XActin supplemented with 10 μM Verapamil for 2 h. After an initial bleaching phase, SiR-XActin reveals improved apparent photostability. **(E)** Combination of SiR-XActin with exchangeable HaloTag Ligands (xHTL) for multi-color STED. Hippocampal rat neurons were transduced with rAAVs with TOM20-HaloTag7 at seven DIV and imaged at twelve DIV. Exchangeable HaloTag Ligand (JF_585_-S5, 500 nM) and SiR-XActin (1 μM) were co-imaged with the same depletion laser (775 nm). Scale bars: 10 μm (overview), 1 μm (magnification).

XActin ligand **7** can serve as a platform to generate XActin probes with different fluorophores, covering the visible to near-infrared spectrum **(Figure SI17-S18, Figure 4)**. Conjugation of **7** to the SiR analog Si-Rhodamine700 yielded SiR700-XActin (**9**), which is further red-shifted relative to SiR-XActin with an excitation maximum at 698 nm. SiR700-XActin showed good labeling properties in fixed and live cells without inducing non-physiological actin phenotypes (**Figure 4, Figure SI19-S20**). Another popular dye for live-cell imaging is the fluorogenic rhodamine derivative MaP555 (**10**), which has an excitation maximum at 554 nm and thus is suitable for dual-color imaging with SiR-based probes using common laser lines.^[42]^ In order to further explore the spectral range of fluorogenic rhodamine dyes, we also coupled XActin to MaP derivatives of carbopyronine (MaP620, **11**) and the green rhodamine analog 500R (MaP500, **12**).^[42]^ All tested probes showed excellent labeling properties in fixed and live cells, including rat hippocampal neurons, and also showed no change in actin phenotype at the concentrations used for imaging (**Figure 4, Figure SI19-SI20**). MaP555- and MaP620-XActin showed satisfying performance in live cell STED imaging (**Figure SI21**). MaP500-XActin and MaP555-XActin stain F-actin in live U-2 OS cells better when incubated with verapamil, showing that also the fluorophore itself contributes to efflux (**Figure 4**).

**Figure 4.**
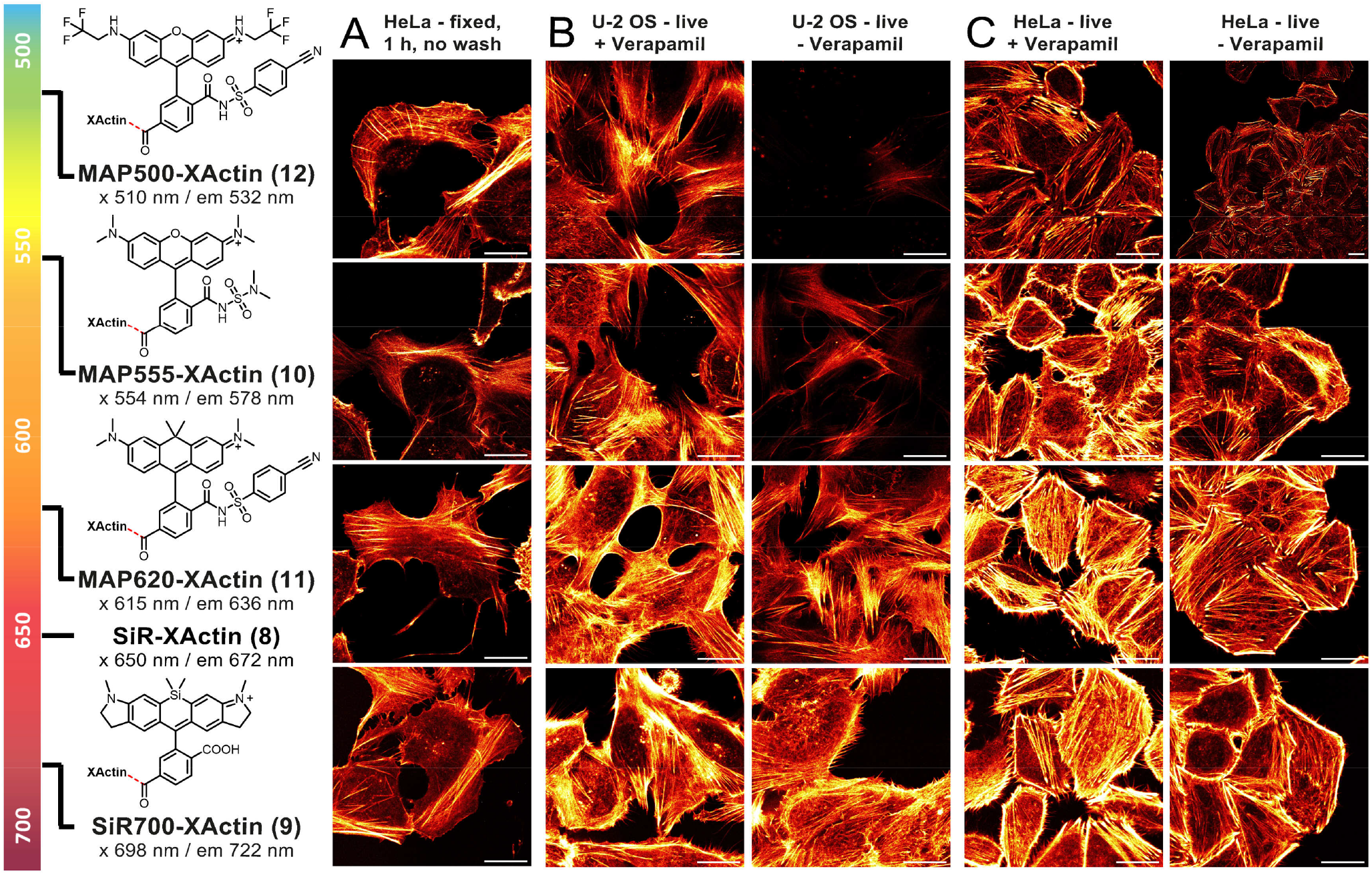
XActin as a conjugate with other fluorophores to cover the whole imaging spectral window: **(A)** Labeling of fixed HeLa cells, **(B)** labeling of live U-2 OS cells or **(C)** HeLa cells with XActin probes, 4 h incubation, with and without 10 μM verapamil. All conditions no-wash, 500 nM. Laser settings live cells: MAP500-XActin 0.3%, MAP555-XActin 0.3%, MAP620-XActin 0.1%, SiR700-XActin 0.3% laser power. Two Line Average. For Laser settings in fixed cells see Figure S19. Red Hot lookup - pixel intensities were scaled between 0 (black) and 65535 (bright yellow) by Fiji. Scale bars: 20 μm.

Next, we investigated the potential of SiR-XActin to image fast actin dynamics as they are observed at the cell leading edge in Cath.a-differentiated (CAD) cells, where the organization of actin, rate of filament assembly, and retrograde flow have been well documented.^[43-45]^ When compared to LifeAct-GFP, a standard tool for visualizing actin in live cells that faithfully recognizes most of the polymerized actin structures in CAD cells,^[46]^ SiR-XActin showed similar labeling of actin filaments at the rapidly assembling leading edge and was remarkably improved at visualizing these structures compared to the synthetic probe SPY-650 FastAct (**Figure 5A, Video SI7**). This was verified by measuring the Mander’s overlap coefficient in colocalization experiments where SiR-XActin and SPY-650 FastAct were used in cells already expressing LifeAct-GFP (**Figure 5B** and **5C, Video SI8**). Additionally, these experiments further reveal how SPY650-FastAct fails to efficiently label leading edge actin populations. Finally, it is always a concern that probes used to visualize actin will alter the properties of the structures they recognize. To determine if SiR-XActin altered rapid actin dynamics in CAD cells, we measured the rate of retrograde flow at the leading edge, which is directly proportional to the rate of actin assembly.^[47]^ There was no difference in cells where actin was labeled with SiR-XActin or LifeAct-GFP (**Figure 5D** and **5E**), and the retrograde flow rates measured with either probe were consistent with previously published results.^[43-45]^ In summary, SiR-XActin provided superior results at labeling leading-edge actin structures in comparison to SPY-650-FastAct, and did not appear to alter the behavior of the structures that it recognized (**Figure 5, Video SI7** and **SI8**).

**Figure 5.**
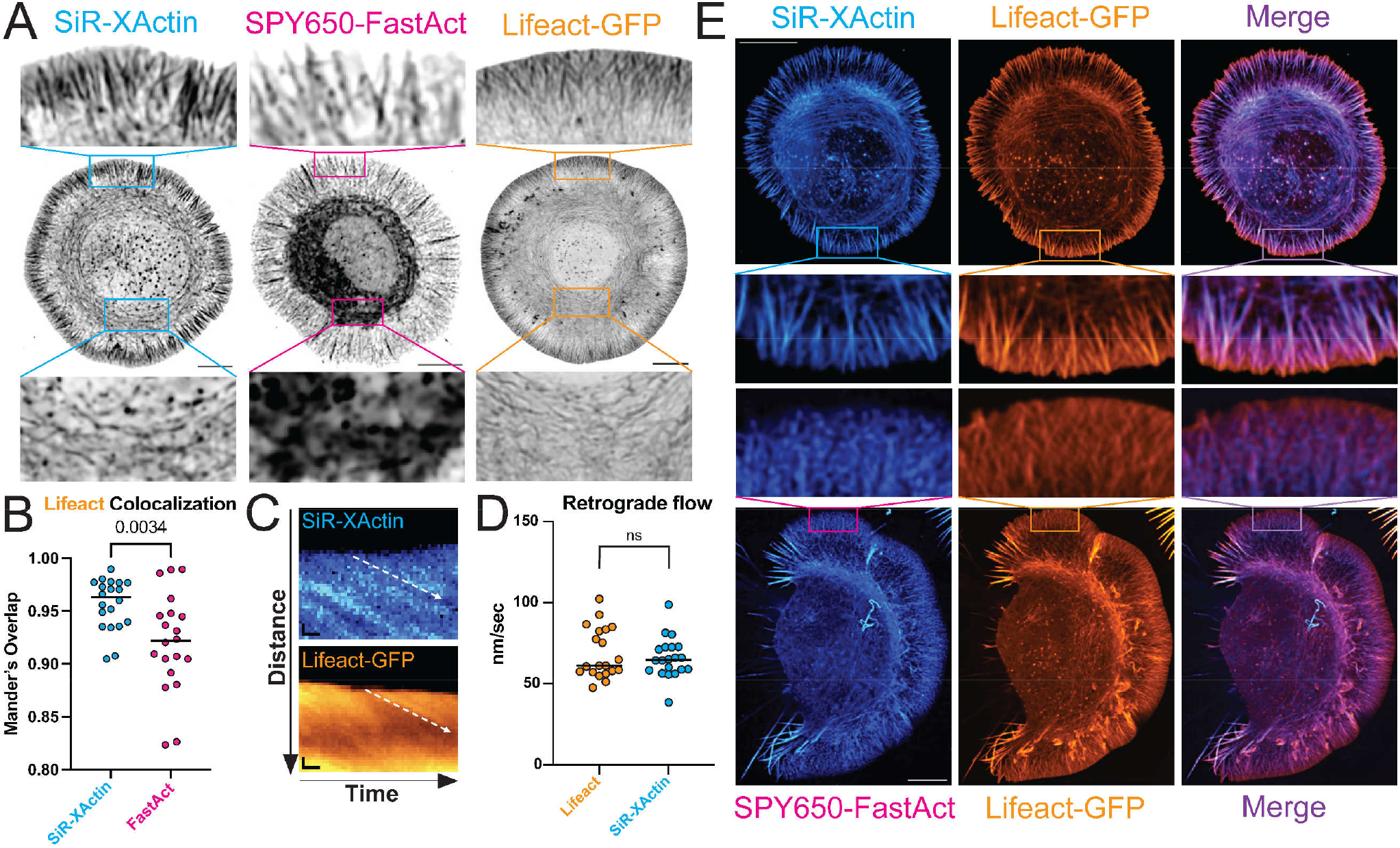
Labeling actin dynamic structures in live CAD cells: **(A)** Representative images of actin imaged with SiR-XActin, SPY650-FastAct, or Lifeact-GFP. Scale bars = 10 μm. **(B)** Colocalization analysis of SiR-XActin and SPY650-FastAct with Lifeact-GFP. n = 20 cells, unpaired t-test. **(C)** Representative kymographs of actin imaged with SiR-XActin and Lifeact-GFP. Scale bars, vertical = 1 μm and scale bars, horizontal = 10 seconds. **(D)** Scatter plot of retrograde flow measurements of Lifeact-GFP and SiR-XActin. Each point represents the average measurement of ten kymographs from each cell. n.s. = not significant, Mann-Whitney test. **(E)** Representative images of cells containing both Lifeact-GFP and SiR-XActin (above) or SPY650-FastAct (below). Scale bar = 10 μm.

## Conclusion

We have developed a new family of synthetic fluorescent probes for imaging F-actin in live cells that distinguish themselves through their low cytotoxicity and efficient labeling in different cell types. As judged by live-cell imaging, labeling of F-actin with SiR-XActin is possible without major changes in the dynamics of actin polymerization and depolymerization. The new probe is compatible with (super-resolution) microscopy and the generation of differently colored XActin derivatives for live-cell imaging is straightforward. SiR-XActin and derivatives of XActin in general should become popular tools for studying the role of actin in various biological processes.

## Supporting information

ESI_SiR-XActin_Nasufovic

## Acknowledgements

This work was supported by the Max Planck Society, the École Polytechnique Fédérale de Lausanne (EPFL) and the University of Jena. V.N. acknowledges support from DAAD during his doctoral studies. J.K. acknowledges support from the Heidelberg Biosciences International Graduate School (HBIGS). Additionally, this work was supported in part by the DFG (EXC 2051, project number 39713860; CRC1127, project number 239745522, to H.-D.A. and by the Maximizing Investigators’ Research Award from the National Institute of General Medical Sciences of the National Institutes of Health under grant number R35GM137959 to E.A. Vitriol. We thank Jana Kress for her help with AAV production. We acknowledge the mass spectrometry facility (S. Fabritz, T. Rudi and J. Kling) for its support. The authors thank all members of the Johnsson laboratory for critical discussions.

## Author Contributions

V.N., H.-D.A., K.J. designed, developed and understood the full potential of XActin ligand and fluorophore conjugates. V.N., J.K., K.J. designed and interpreted the experiments for profiling the properties of conjugates. V.N. synthesized all molecules reported in this study. M.B. performed cell viability experiments. J.K. performed STED imaging and photophysical characterization of conjugates. V.N., J.K., B.K, performed CLSM and widefield experiments for profiling the properties of conjugates. V. L-D., E. A. C-A., M. M., K. W. performed wound healing experiments. H. L. and E.A.V. designed and performed labeling actin dynamic structures in live CAD cells. V.N. and K.J. wrote the manuscript with input from all authors.

