## Supplementary material for "SiR-XActin: A fluorescent probe for imaging actin dynamics in live cells": ESI_SiR-XActin_Nasufovic

for

#### 1 Supplementary Figures

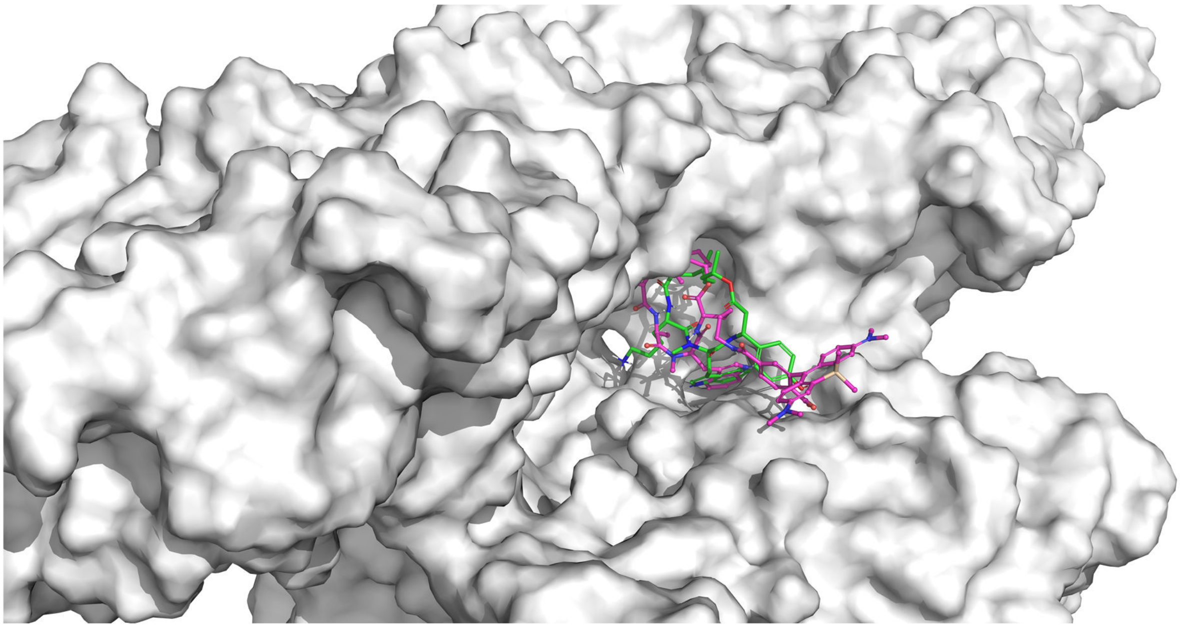

**Figure SI1. Ligands in the F-Actin binding cleft:** Cryo-EM binding structure of jasplakinolide represented in green (PDB 6T24) and docking based binding model of SiR-XActin (violet).

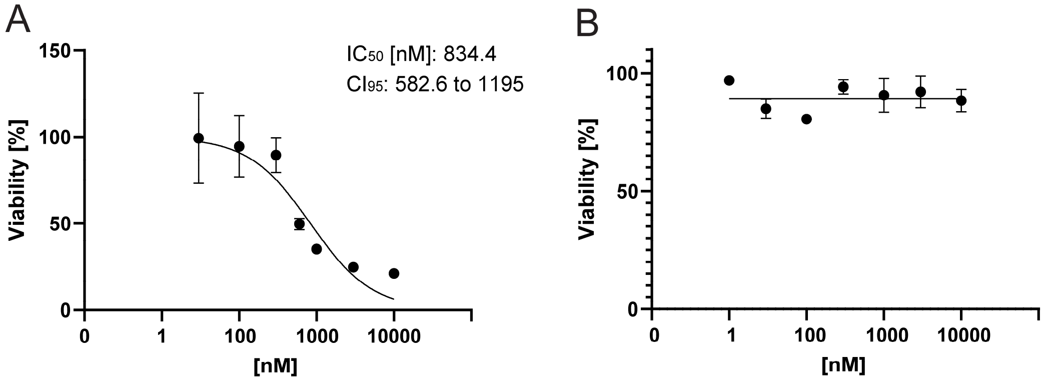

**Figure SI2. Cytotoxicity of SiR-actin and SiR-XActin:** Influence of SiR-Actin (A) and SiR-XActin (B) on viability of HeLa cells measured with a MTT viability assay for a tested range of 10 nM – 10,000 nM. For each concentration, point values are shown as the mean percentage of viable cells relative to controls ± standard deviation (SD) from a representative experiment out of three biological replicates performed in technical quadruplets. Curve fitting was conducted using GraphPad Prism.

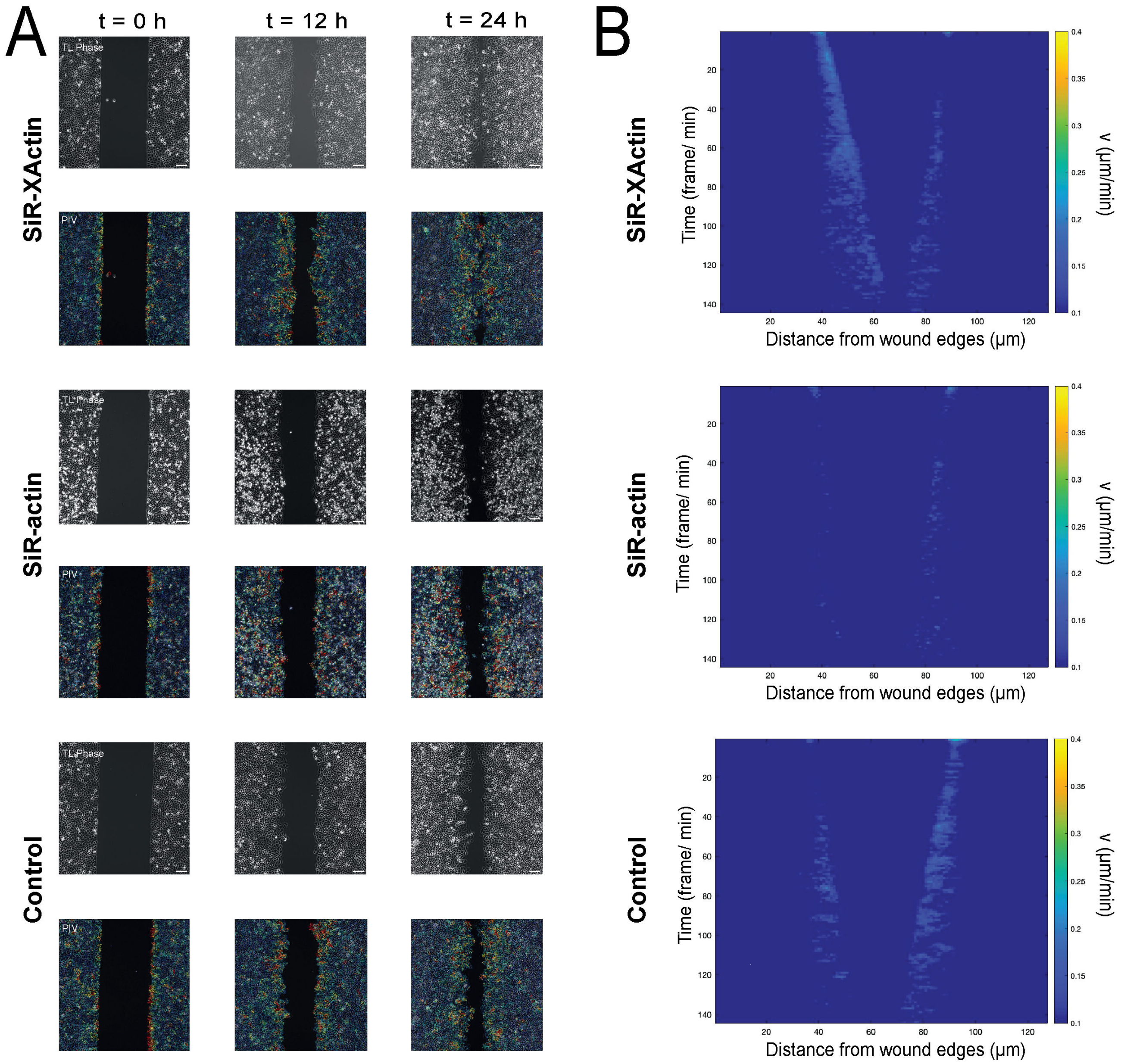

**Figure SI3. Wound healing assay in HeLa cells using SiR-actin and SiR-XActin: (A)** Bright-field images of HeLa cell-collective migration in a wound, created by insertion of a stencil of 500 µm in width. Images show the time-dependent differences following 12-h preincubation with SiR-actin, SiR-XActin or DMSO control (500 nM), scale bars: 100 nm. **(B)** Kymographs were generated for each field of view along the wound edge in time-lapse microscopy images. The field of view was orientated perpendicular to the wound axis, and pixel intensity profiles were extracted along the region for each frame. These profiles were sequentially stacked to create a spatiotemporal representation of cell movement and dynamics over time.

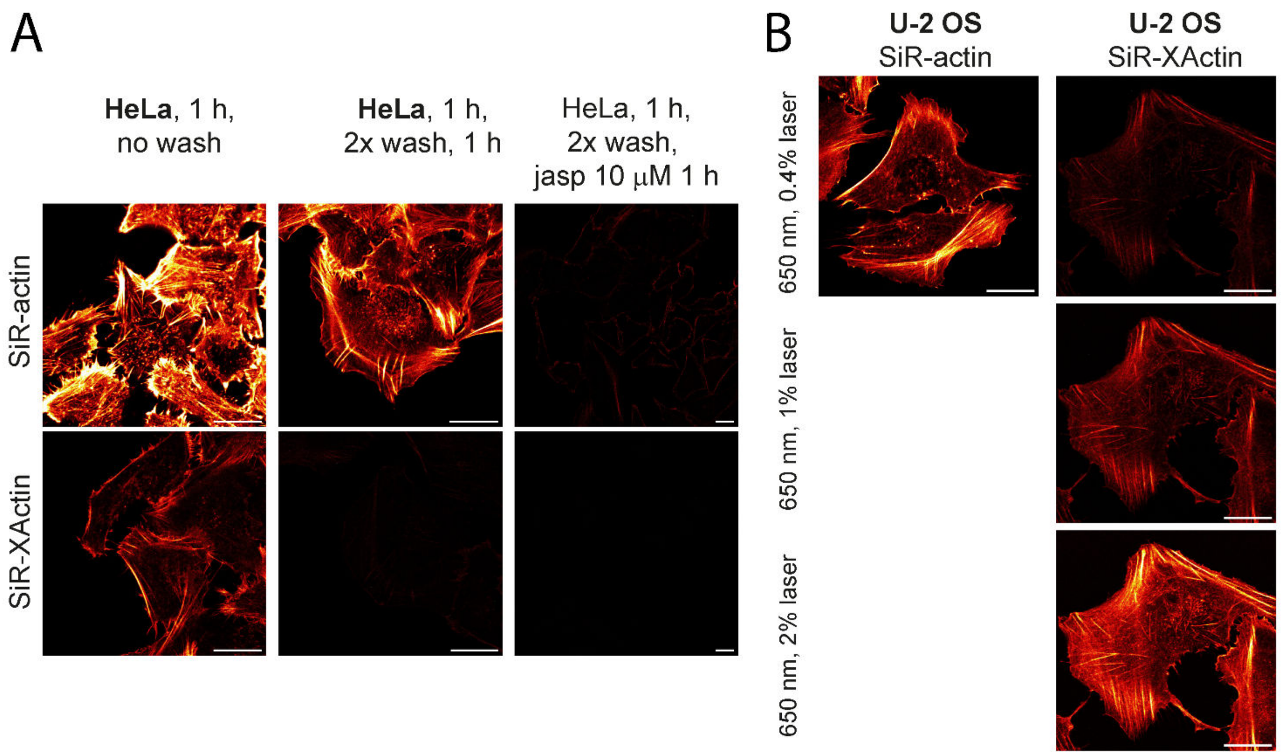

**Figure SI4. Comparison of SiR-actin and SiR-XActin in labelling of fixed cells: (A)** HeLa cells imaged with 0.4% laser power, excitation at 652 nm. **(B)** Fixed-cell no-wash CLSM imaging of U-2 OS cells: Laser settings: SiR-actin 0.4%, SiR-XActin 0.4%, 1% and 2% laser power. Red Hot lookup - pixel intensities were scaled between 0 (black) and 65535 (bright yellow) by Fiji. Scale bars 20 μm.

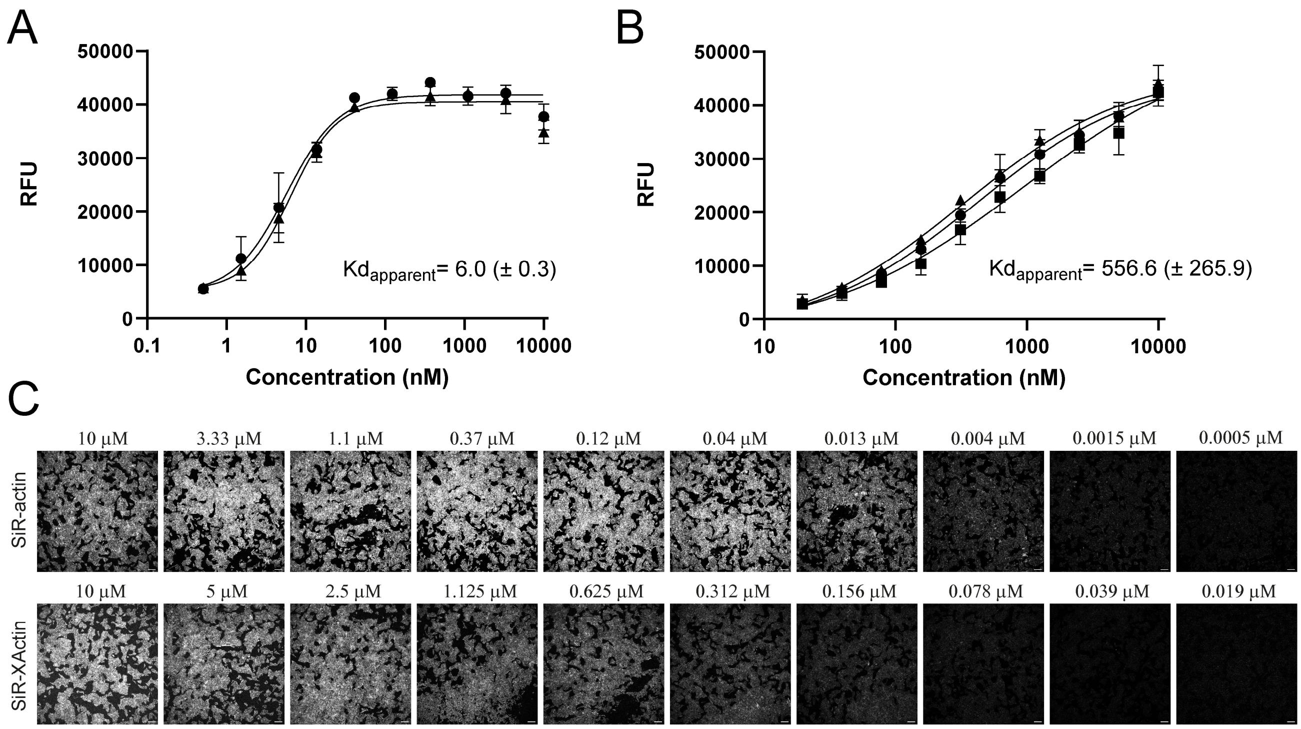

**Figure SI5. SiR-actin and SiR-XActin affinity ($Kd_{app}^{F\text{-}actin}$) determination in fixed U-2 OS cells:**[1] All experiments were performed in technical triplicates and two biological replicates for SiR-actin (A), three biological replicates for SiR-XActin (B). Median fluorescence intensity values (ImageJ 1.54f) of triplicates were fitted by sigmoidal dose response curve fit (GraphPad Prism) to obtain $EC_{50}$ values ($Kd_{app}^{F\text{-}actin}$), average values with SD are shown. (C) One representative dose-response image serial for each probe, Pixel Size 1.514 µm. Scale bars 20 µm.

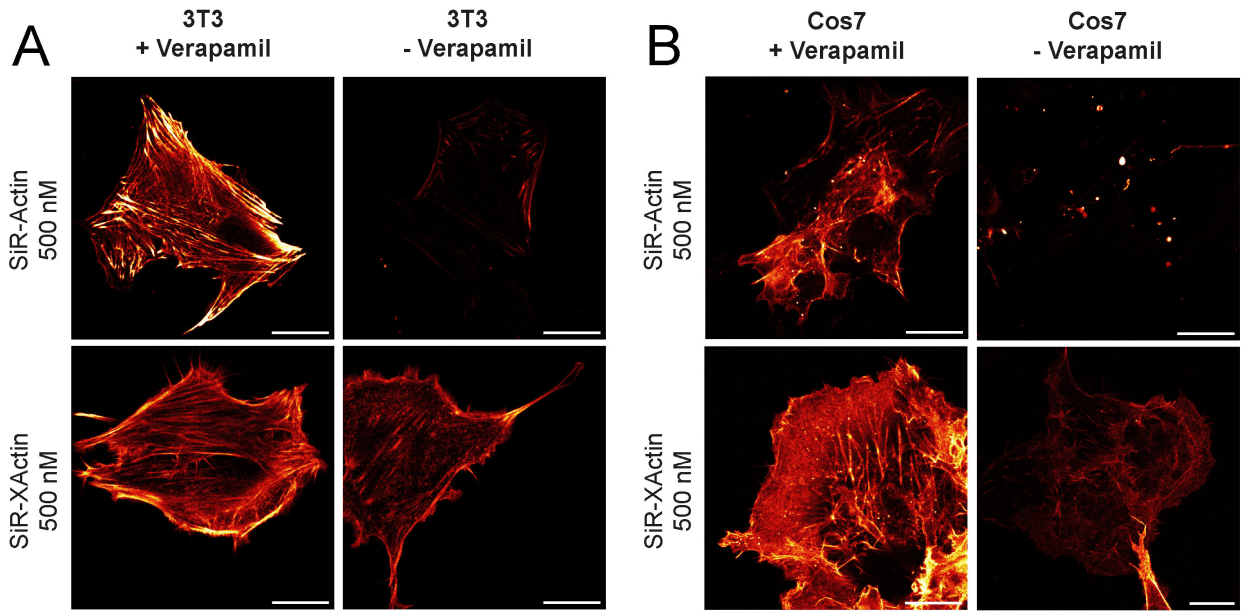

**Figure SI6. Labelling performance of SiR-actin and SiR-XActin: (A)** Live-cell no-wash CLSM imaging of 3T3 and (B) COS7 cells stained with SiR-actin and SiR-XActin in presence and absence of verapamil (10 µM), 6 h incubation. Red Hot lookup - pixel intensities were scaled between 0 (black) and 65535 (bright yellow) by Fiji. Laser settings: Laser settings: SiR-XActin 1% laser power. Scale bars 20 µm.

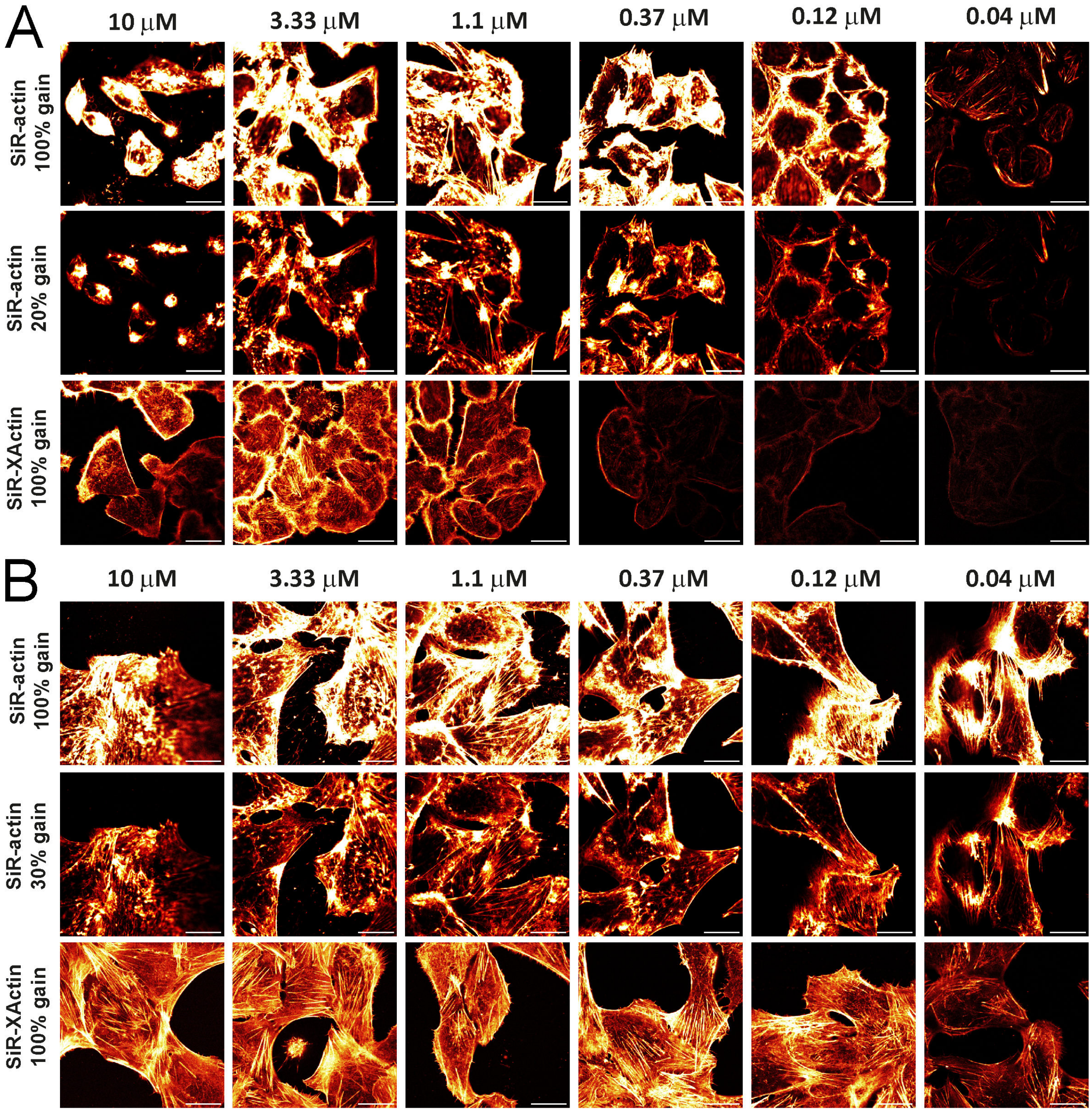

**Figure SI7. Phenotypic differences after incubation with varying concentrations of SiR-actin and SiR-XActin:** Live cell no-wash CLSM imaging of **(A)** HeLa cells (6 h incubation time) and **(B)** U-2 OS cells (16 h incubation time) stained with SiR-actin and SiR-XActin. In case of U-2 OS cells, Verapamil (10 μM) was used. Red Hot lookup - pixel intensities were scaled between 0 (black) and 65535 (bright yellow) by Fiji. Scale bars 20 μm.

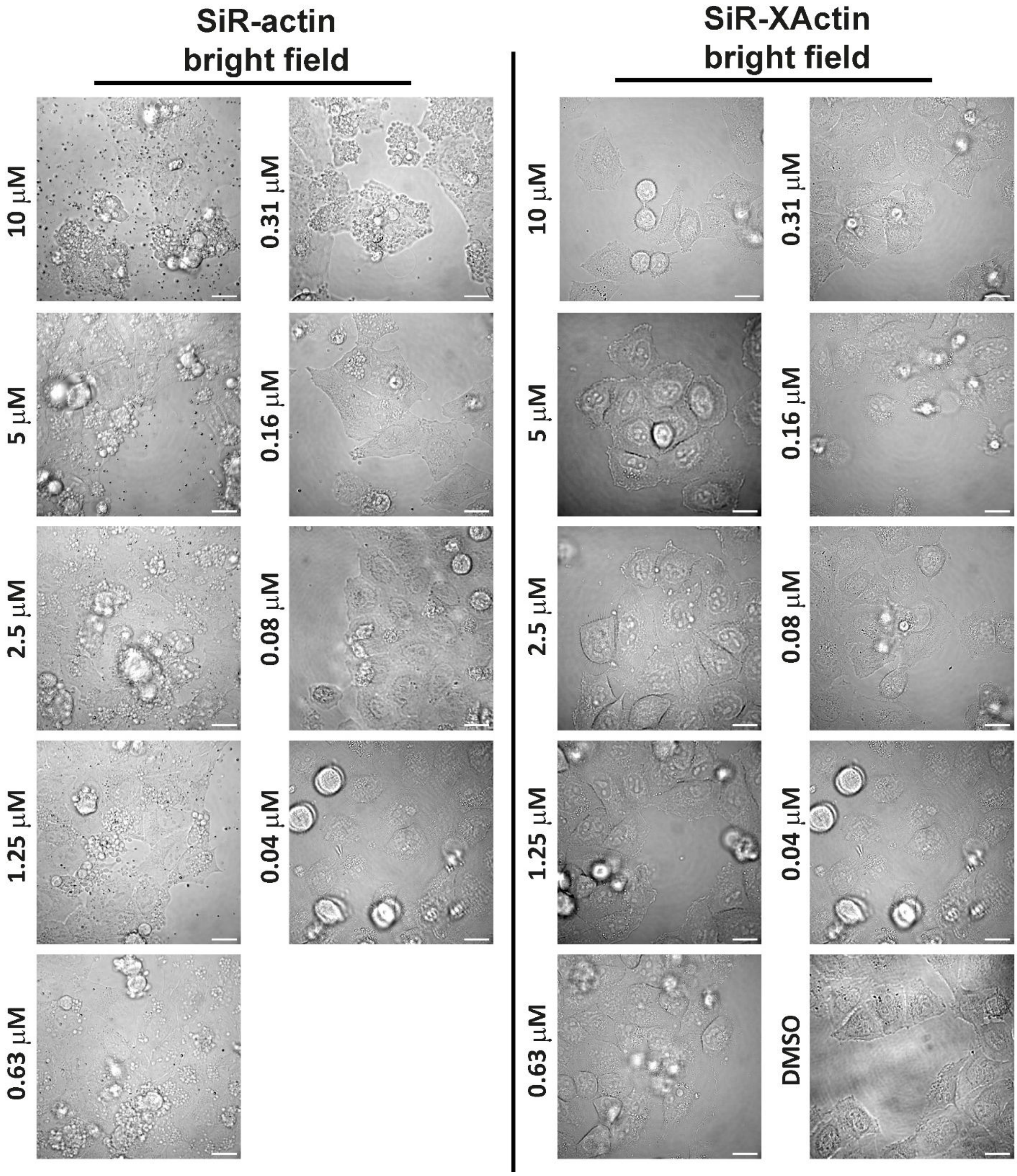

**Figure SI8. Imagining of blebbing phenotype in HeLa cells:** Live cell no-wash brightfield imaging of HeLa (6 h incubation time) cells stained with SiR-actin and SiR-XActin. Scale bars 20 µm.

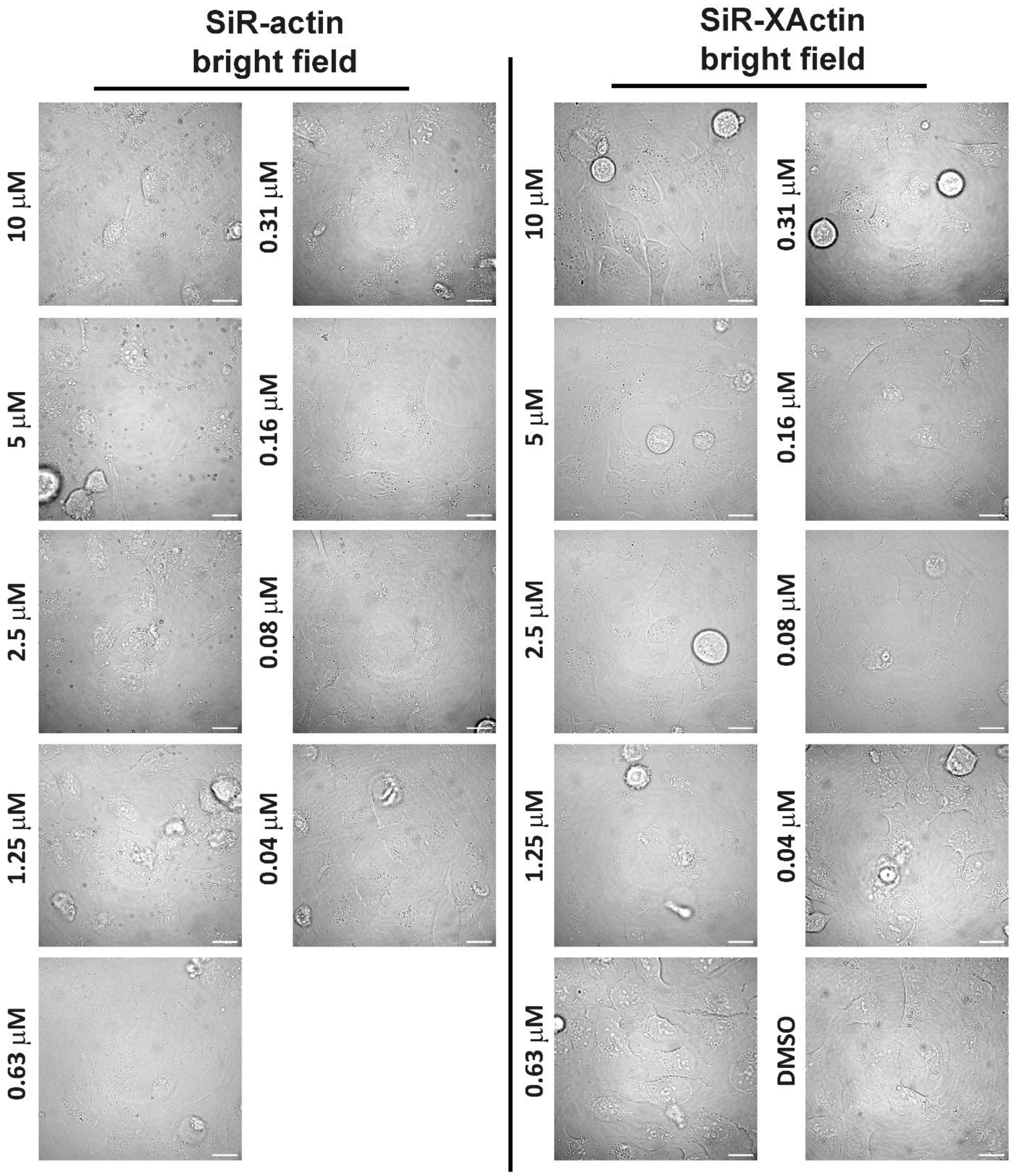

**Figure SI9. Imagining of blebbing phenotype in U-2 OS cells:** Live cell no-wash brightfield imaging of U-2 OS cells (16 h incubation time) stained with SiR-actin and SiR-XActin. Verapamil (10 µM) was used. Scale bars 20 µm.

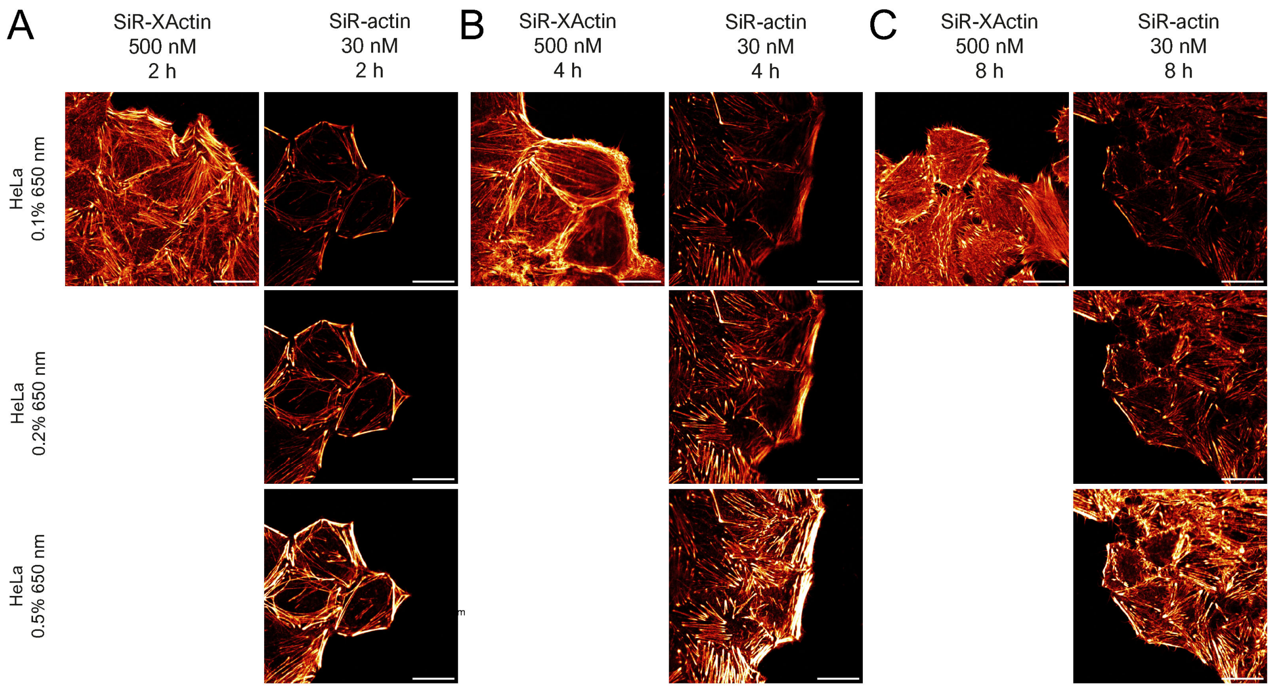

**Figure SI10.** Live cell no-wash CLSM imaging of F-actin HeLa cells with 500 nM of SiR-XActin or 30 nM of SiR-actin: incubation of 2 h (A), 4 h (B), and 8 h (A). Red Hot lookup - pixel intensities were scaled between 0 (black) and 65535 (bright yellow) by Fiji. Scale bars 20 µm.

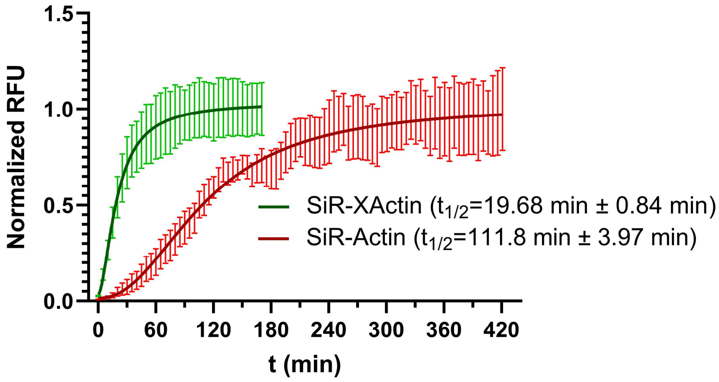

**Figure SI11. Live-cell permeation of SiR-actin and SiR-XActin**: Fluorescent signal increase was followed by CLSM in live HeLa cells, both probes used in 500 nM, signal was normalised to mean end point fluorescence signal. Sigmoidal curve fit was performed and $t_{1/2}$ extrapolated by GraphPad Prism.

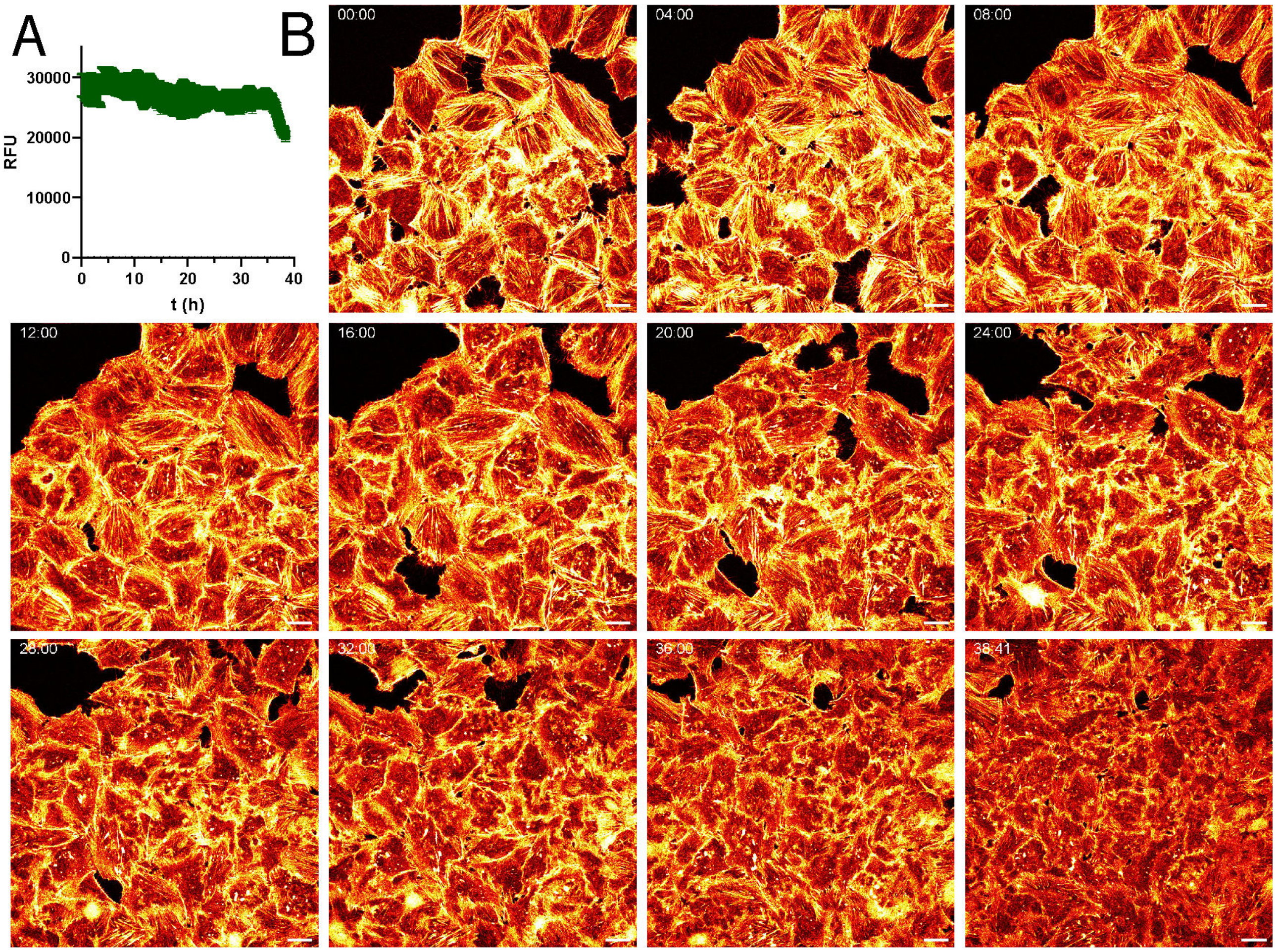

**Figure SI12. Time-lapse live-cell no-wash imaging of HeLa cells labelled with SiR-XActin**: (A) Average fluorescence signal stability during imaging, (B) Images of representative time points. imaging started after incubation with 1 µM of SiR-XActin for 24 h. imaging every 30 s for 40 h. Scale bars 20 µm. Red Hot lookup - pixel intensities were scaled between 0 (black) and 65535 (bright yellow) by Fiji. Please see Video SI3.

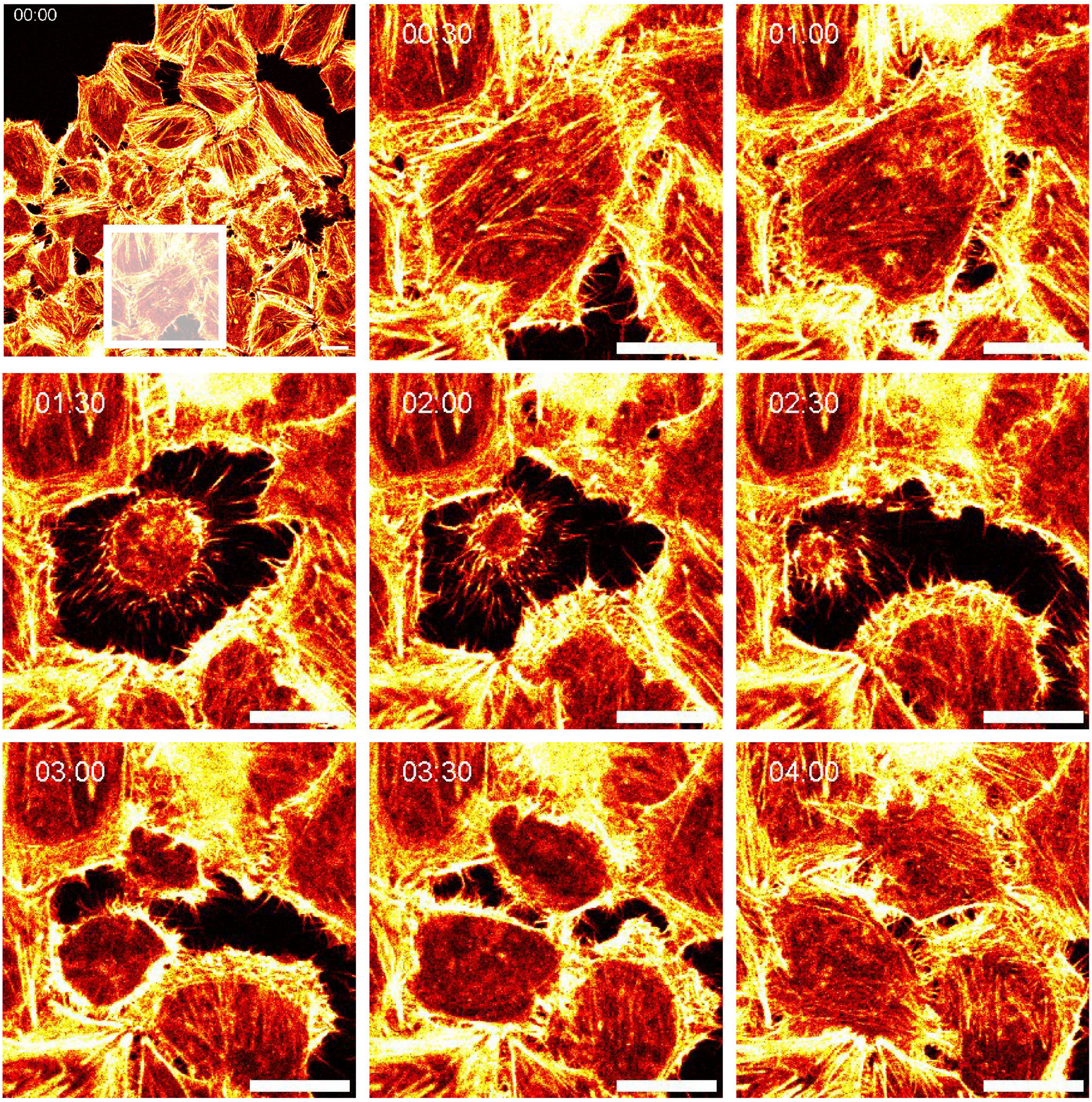

**Figure SI13. Time-lapse live-cell no-wash imaging of highly dynamic actin structures in HeLa cells labelled with SiR-XActin** (enlarged region from Video SI3): Images of representative time points chosen to highlight dynamic F-actin structures. White transparent rectangle indicates region which was enlarged. Please see Video SI3, SI4. Red Hot lookup - pixel intensities were scaled between 0 (black) and 65535 (bright yellow) by Fiji. Scale bars 20 µm.

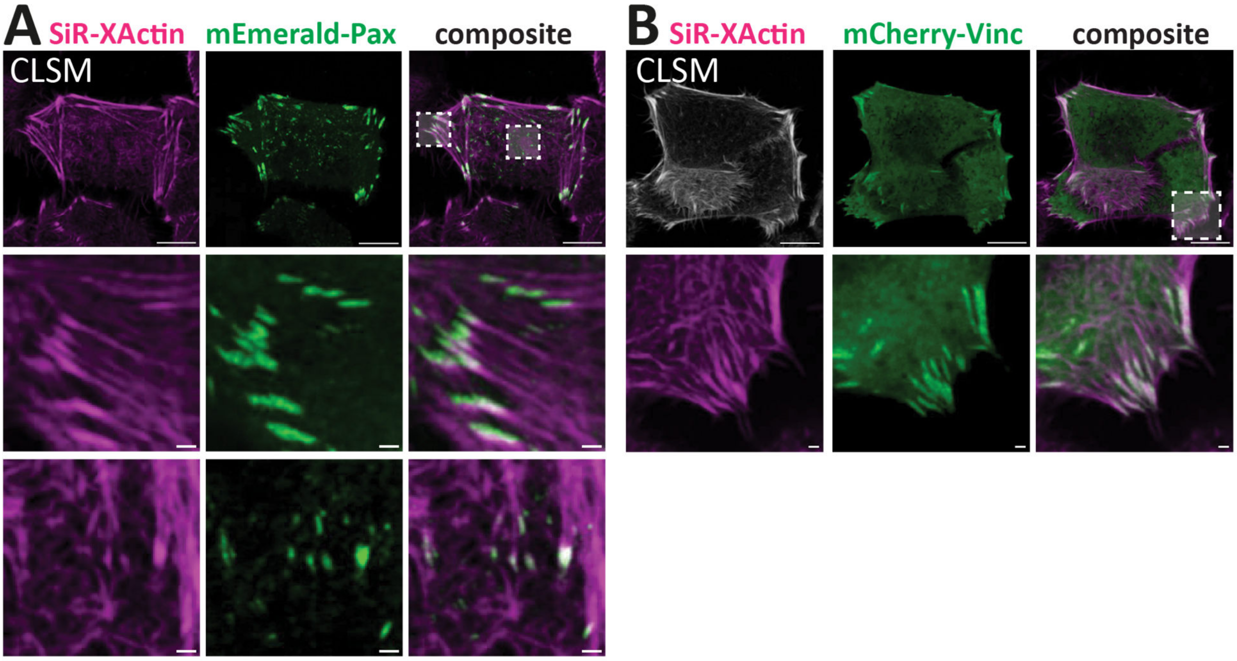

**Figure SI14. Co-imaging of SiR-XActin and focal adhesion marker in HeLa cells:** Combination of SiR-XActin (magenta) with transiently expressed **(A)** mEmerald-Paxilin (green) or (B) mCherry-Vinculin (green) for dual-color CLSM imaging.[2] SiR-XActin (1 µM) Scale bars: 10 µm (overview), 2 µm (magnification). Please see Supplementary Table 1 for imaging details.

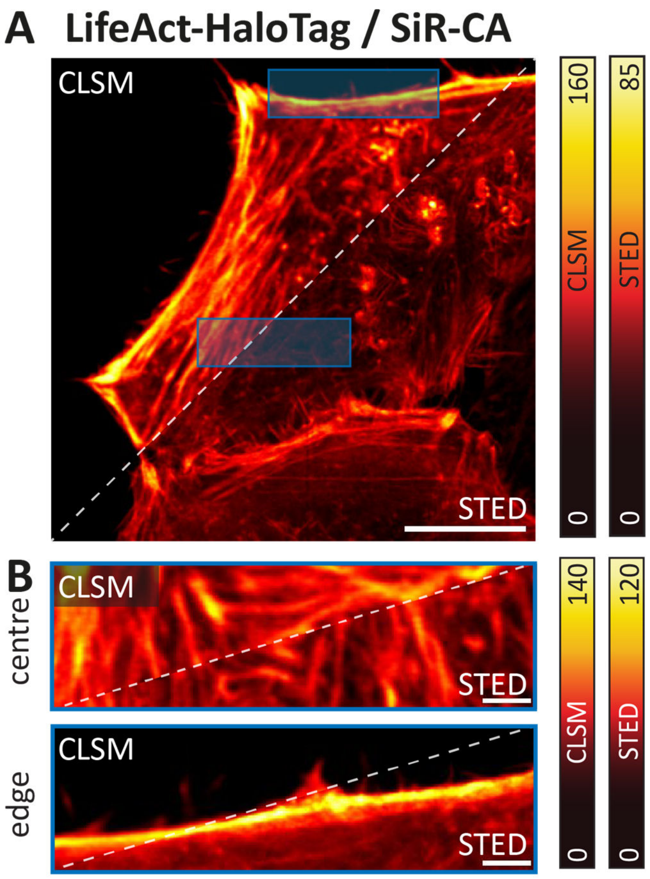

**Figure SI15. Genetically-encoded LifeAct-HaloTag7 imaged in live cells by STED microscopy:** (A) Phenotypic analysis of LifeAct-HaloTag7 labelling in live HeLa cells. Cells were transduced with AAVs and stained with 500 nM SiR-HTL probes for 2 h, no-wash imaging. Representative CLSM and STED images. Pixel intensities were scaled between dark and bright colours according to the reference bars. Scale bars: 10 µm. (B) Magnification of blue ROIs from A. Scale bars: 1 µm. Please see Supplementary Table 1 for imaging details.

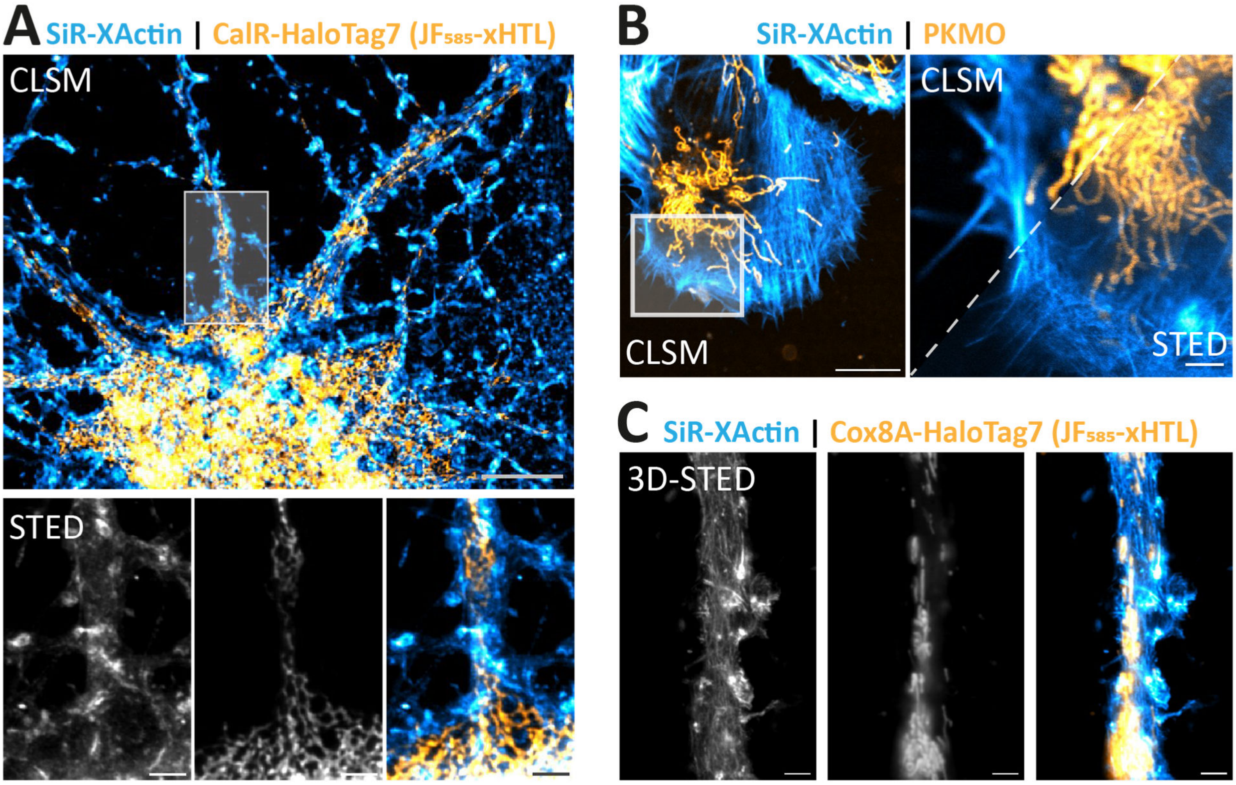

**Figure SI16. Dual-color STED microscopy of SiR-XActin and exchangeable labels. (A)** Combination of SiR-XActin with exchangeable HaloTag Ligands (xHTL) for multi-color STED. Hippocampal rat neurons were transduced with rAAVs with CalR-HaloTag7-KDEL at 7 DIV and imaged at 12 DIV. Exchangeable HaloTag Ligand (orange, $JF_{585}$-S5, 500 nM) and SiR-XActin (cyan, 1 µM). **(B)** Combination of SiR-XActin with PKMO[3] for transfection-free multi-color STED in U-2 OS cells. PKMO (orange, 10 nM) and SiR-XActin (cyan, 1 µM). **(C)** 3D-STED with exchangeable HaloTag Ligands (xHTL) for multi-color STED. Experimental details as in (A). Max. projection of 20 z-stacks. Scale bars: 10 µm (overview), 1 µm (magnification). Both dyes were co-imaged with the same depletion laser (775 nm). Please see Supplementary Table 1 for imaging details.

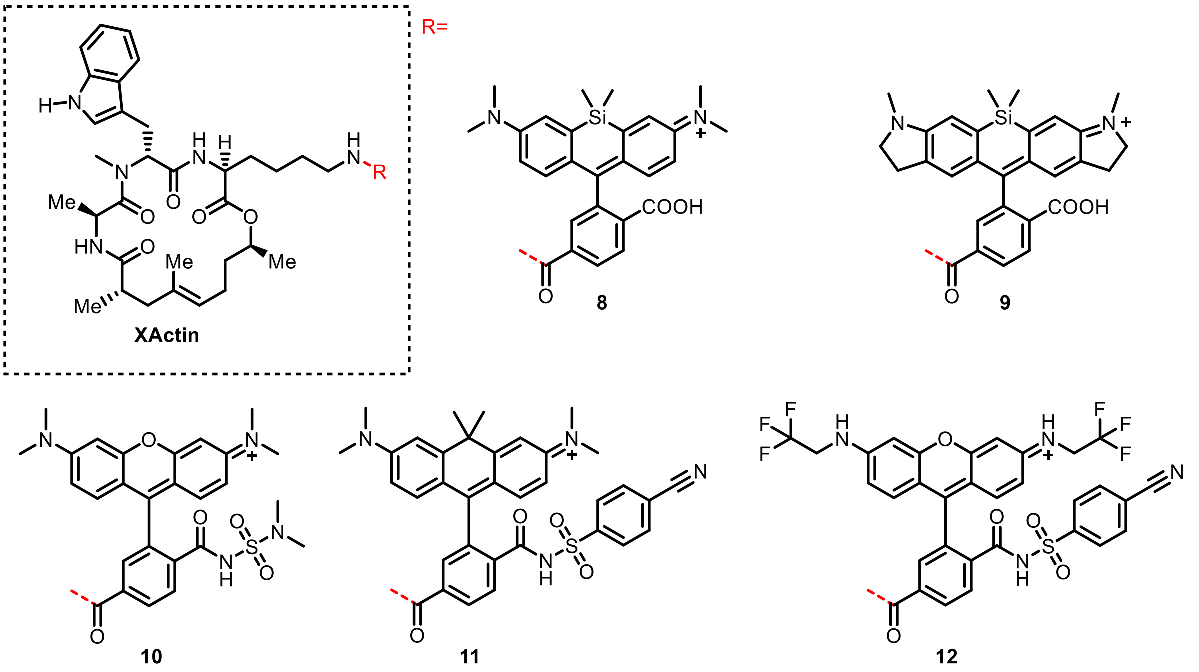

**Figure SI17**. **XActin conjugates synthesized and tested in this study**

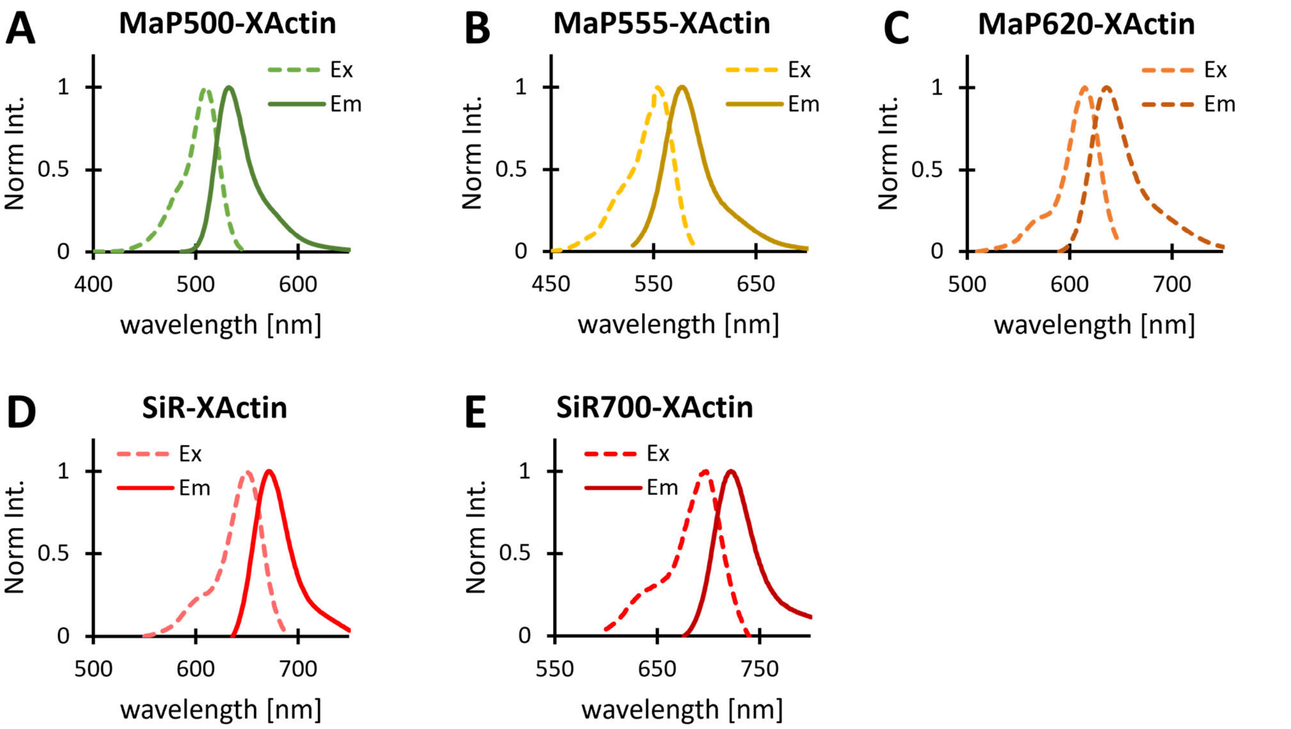

**Figure SI18. Fluorescence excitation and emission profiles of XActin derivatives.:** 1 µM dyes in 50 mM HEPES, 50 mM NaCl, 0.1% DMSO, 0.1% SDS, pH 7.3. Excitation and emissions spectra were recorded with 1 nm step size on a Spark20M Tecan plate reader using the detection or excitation settings summarized in Table SI2.

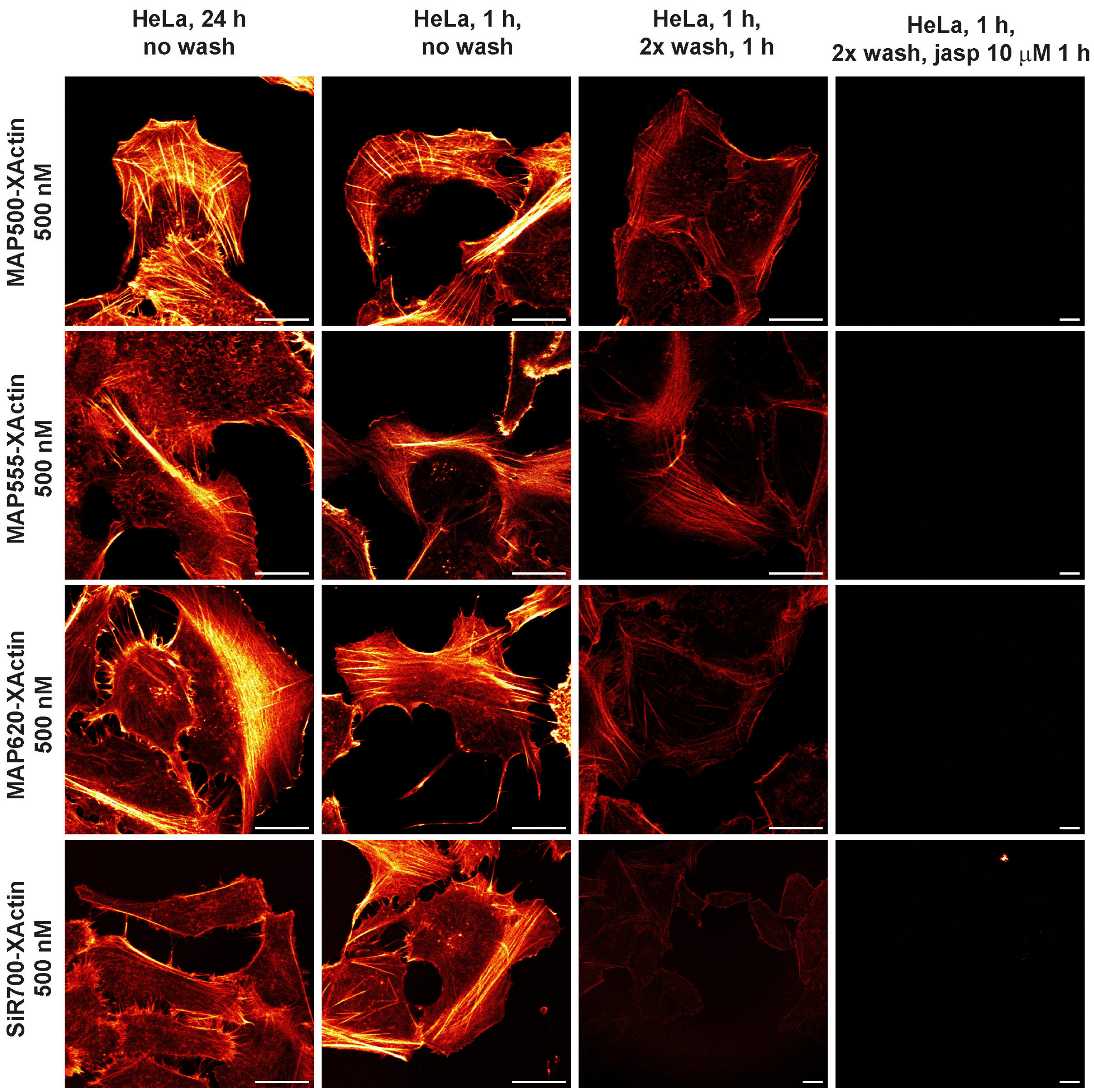

**Figure S19. XActin as a platform for a fluorophore of a choice - Imaging of fixed HeLA cells by no-wash CLSM:** Fixed cells (4% PFA in PBS for 15 min) and washed with PBS (3x) were labelled with corresponding probes in PBS supplemented with 1% of BSA. Laser settings: MAP500-XActin 5%, MAP555-XActin 2%, MAP620-XActin 0.5%, SiR700-XActin 6% laser power. Two Line Average. Red Hot lookup - pixel intensities were scaled between 0 (black) and 65535 (bright yellow) by Fiji. Scale bars: 20 µm (overview).

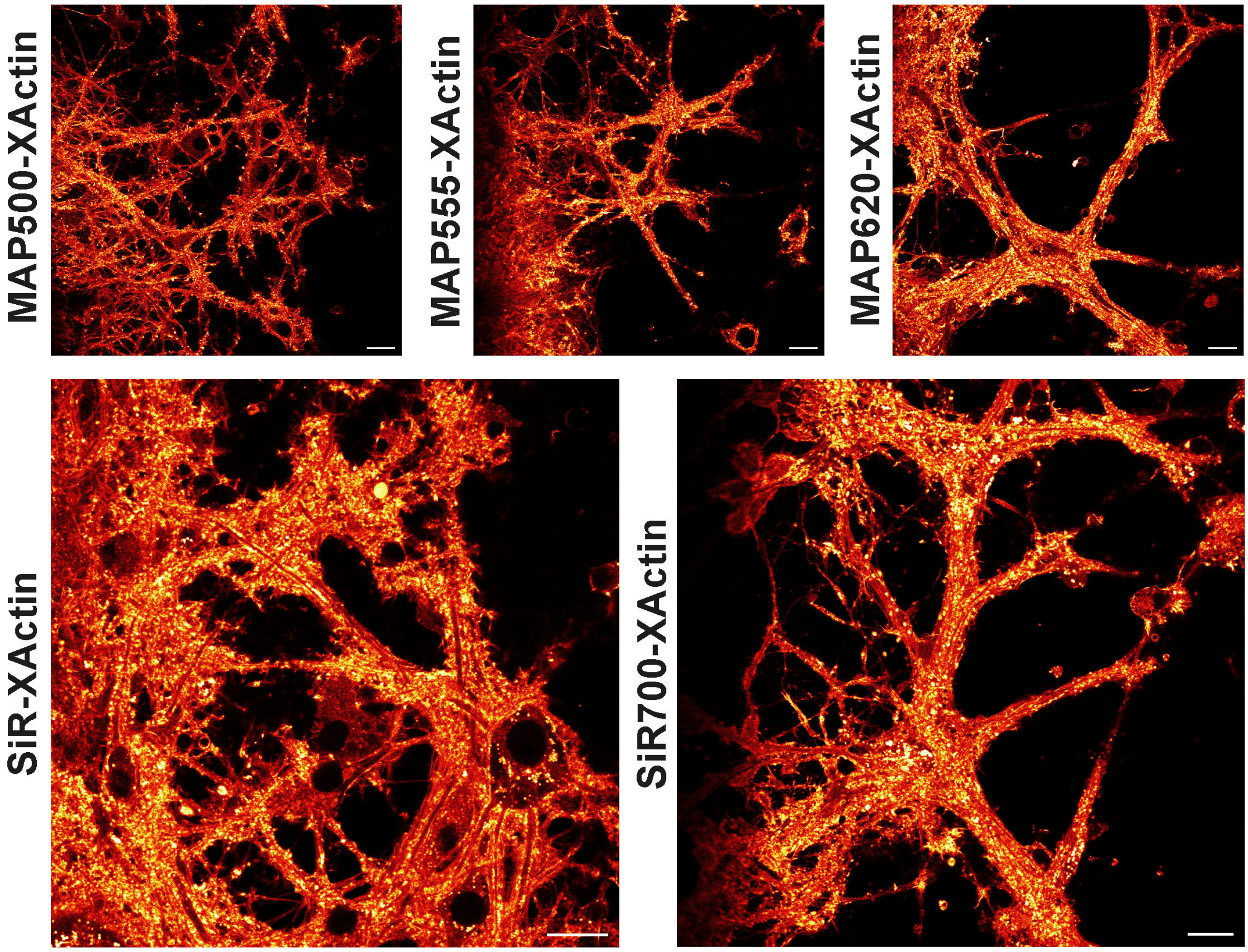

**Figure S20. No-wash imaging of live rat hippocampal neurons by CLSM:** Hippocampal rat neurons (14 DIV) were labelled by corresponding XActin probe (500 nM) and imaged after 2 h. Laser settings: MAP500-XActin 0.5%, MAP555-XActin 1%, MAP620-XActin 0.2%, SiR-XActin 0.6%, SiR700-XActin 0.8% laser power. Four Line Average was used. Red Hot lookup - pixel intensities were scaled between 0 (black) and 65535 (bright yellow) by Fiji. Scale bars: 20 µm.

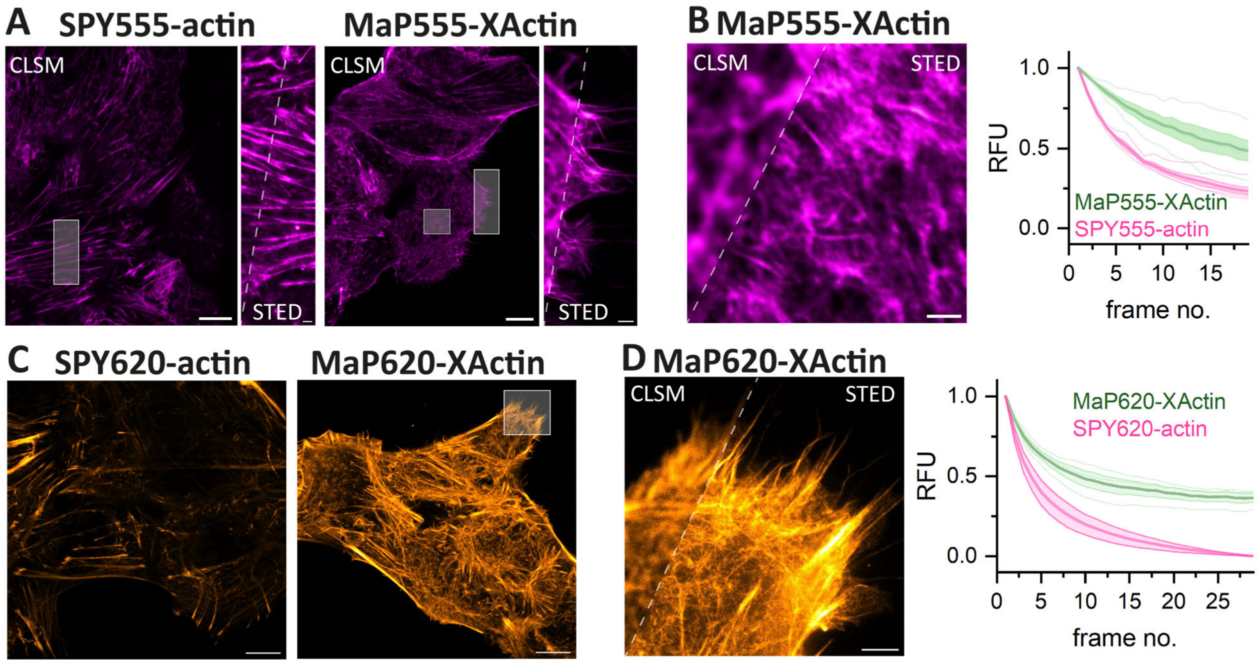

**Figure S21. Live-cell STED microscopy of U-2 OS cell with XActin probes of different colors. (A)** U-2 OS cells were stained with 1 μM SPY555-actin or MaP555-XActin for 2 h, no-wash imaging. Representative CLSM and STED images. Min. pixel intensity: 0. Max. pixel intensities: 700 (overview), 350 (CLSM), 220 (STED). Scale bars: 10 μm (overview) 1 μm (zoom). **(B)** Normalized fluorescence intensity of multi-frame STED imaging in a 10x10 μm ROI. **(C)** U-2 OS cells were stained with 1 μM SPY620-actin or MaP620-XActin or supplemented with 10 μM Verapamil for 2 h. Min. pixel intensity: 0. Max. pixel intensities: 1200 (overview), 400 (CLSM), 330 (STED). Scale bars: 10 μm (overview) 1 μm (zoom). **(D)** Same experiment as in B. Overall, SiR-XActin reveals improved apparent photostability for both dyes. Please see Supplementary Table 1 for imaging details.

#### 3 Supplementary Videos

**Video SI1:** Permeation of SiR-XActin in live HeLa cells followed by CLSM. 500 nM of SiR-XActin added in medium, 4 Z-planes imaged every 5 min, one representative plane time lapse shown, 1 fps. Scale bar: 50 µm. Excitation at 652 nm 0.1% laser power. Time is listed in hour:min.

**Video SI2:** Permeation of SiR-actin in live HeLa cells followed by CLSM. 500 nM of SiR-XActin added in medium, 4 Z-planes imaged every 5 min, one representative plane time lapse shown, 1 fps. Scale bar: 50 µm. Excitation at 652 nm 0.1% laser power. Time is listed in hour:min.

**Video SI3:** Time-lapse live-cell no-wash imaging of HeLa cells after 24 h incubation with 1 µM of SiR-XActin. Single Z-plane was imaged every 30 s for 40 h. For shown video 100 fps used, jpg compression used in Fiji. Time is listed in hour:min.

**Video SI4:** Enlarged region of Video SI3 with time-lapse live-cell no-wash imaging of HeLa cells after 24 h incubation with 1 µM of SiR-XActin. For shown video 100 fps used, jpg compression used in Fiji. Time is listed in hour:min.

**Video SI5:** Bleaching resistance comparison of SiR-XActin and SiR-actin staining in multi-frame STED. Live U-2 OS cell were stained with 1 µM actin probes for 3 h in presence of 10 µM Verapamil and respectively imaged in a 10 x 10 µm ROI over 20 consecutive frames. Frame number given in top right corner. Scale bar 2 µm. Please see Supplementary Table 1 for imaging details.

**Video SI6:** 3D-STED imaging of Actin and the ER. Hippocampal rat neurons were transduced with rAAVs with CalR-HaloTag7-KDEL at 7 DIV and imaged at 12 DIV. Exchangeable HaloTag Ligand (orange, $JF_{585}$-S5, 500 nM) and SiR-XActin (cyan, 1 µM) were co-imaged with the same depletion laser (775 nm) over 24 z-frames. 3D projections. Please see Supplementary Table 1 for imaging details.

**Video SI7:** Representative timelapses of CAD cells treated with SiR-XActin (ledt) or SPY650-FastAct (right). Time is listed in min:sec.

**Video SI8:** Timelapse of a CAD cell treated with SiR-XActin and expressing LifeAct-GFP. Time is listed in min:sec.

### 4 Supplementary Tables

**Table SI1:** Settings used for CLSM and STED microscopy:

| Fig. / Vid. | Dye | Excitation [nm] (%) | Emission [nm] | STED [nm] (%) | Pixel dwell time [µs] | Pixel size x-y / z [nm] | Size x-y-z [µm] | Comment | Set-up |
|---|---|---|---|---|---|---|---|---|---|
| **3A** | SiR | 640 (2) | 650 - 757 | - | 7 | 60 | 40 x 40 | CLSM, 8 Line accu. | **A** |
| | | 640 (3.5) | | 775 (20) | | | | STED, 8 Line accu. | |
| **3B** | SiR | 640 (2) | 650 - 757 | - | 8 | 50 | 20 x 6 | CLSM, 6 Line accu. | **A** |
| | | 640 (3.5) | | 775 (20) | | | | STED, 6 Line accu. | |
| **3D / Vid. S5** | SiR | 640 (3.5) | 650 - 757 | 775 (20) | 15 | 30 | 10 x 10 | STED, 3 Line accu. | **A** |
| **3E** | $JF_{585}$ | 561 (20) | 571 - 630 | - | 15 | 60 | 70 x 70 (ov) 25 x 12(z) | CLSM, 3 Line accu. | **A** |
| | | 561 (90) | | 775 (40) | | 40 | | STED, 3 Line accu. | |
| | SiR | 640 (2) | 660 - 746 | - | | 60 | | CLSM, 3 Line accu. | |
| | | 640 (5) | | 775 (20) | | 40 | | STED, 3 Line accu. | |
| **S14A** | mEm | 485 (40) | 495 – 571 | - | 15 | 80 (ov) 40 (z) | 50 x 50 (ov) 10 x 10 (z) | CLSM, 3 Line accu. | **A** |
| | SiR | 640 (0.2) | 650 - 757 | | | | | | |
| **S14B** | mChe | 561 (20) | 571 - 630 | - | 15 | 80 (ov) 40 (z) | 50 x 50 (ov) 18 x 18 (z) | CLSM, 3 Line accu. | **A** |
| | SiR | 640 (0.2) | 650 - 757 | | | | | | |
| **S15** | SiR | 640 (2) | 650 - 757 | - | 7 | 60 | 40 x 40 | CLSM, 8 Line accu. | **A** |
| | | 640 (3.5) | | 775 (20) | | | | STED, 8 Line accu. | |
| **S16A** | $JF_{585}$ | 561 (20) | 571 - 630 | - | 15 | 60 | 80 x 80 (ov) 10 x 15 (z) | CLSM, 3 Line accu. | **A** |
| | | 561 (90) | | 775 (40) | | 40 | | STED, 3 Line accu. | |
| | SiR | 640 (2) | 660 - 746 | - | | 60 | | CLSM, 3 Line accu. | |
| | | 640 (5) | | 775 (20) | | 40 | | STED, 3 Line accu. | |
| **S16B** | PKMO | 561 (20) | 571 - 630 | - | 15 | 60 | 80 x 80 (ov) 40 x 40 (z) | CLSM, 3 Line accu. | **A** |
| | | 561 (35) | | 775 (20) | | 40 | | STED, 3 Line accu. | |
| | SiR | 640 (1) | 660 - 747 | - | | 60 | | CLSM, 3 Line accu. | |
| | | 640 (3) | | 775 (20) | | 40 | | STED, 3 Line accu. | |
| **S16C** | $JF_{585}$ | 561 (20) | 571 - 630 | - | 15 | 60 | 10 x 15 | CLSM, 3 Line accu. | **A** |
| | | 561 (90) | | 775 (40) | | 40 | | STED, 3 Line accu. | |
| | SiR | 640 (2) | 660 - 746 | - | | 60 | | CLSM, 3 Line accu. | |
| | | 640 (5) | | 775 (20) | | 40 | | STED, 3 Line accu. | |
| **Vid. SI6** | $JF_{585}$ | 561 (90) | 571 - 630 | 775 (40) | 15 | 40 / 100 | 10 x 7.5 x 2.4 | 80% 3D-STED, 3 Line accu | **A** |
| | SiR | 640 (5) | 660 - 746 | 775 (20) | | | | | |
| **S21A** | MaP555 | 561 (10) | 571 – 650 | - | 15 | 50 | 80 x 80 (ov) 10 x 10 (z) | CLSM, 3 Line accu. | **B** |
| | | 561 (26) | | 660 (25) | | 30 | | STED, 3 Line accu | |
| **S21B** | | 561 (10) | | - | | 50 | 6 x 18 (z) | CLSM, 3 Line accu. | |
| | | 561 (26) | | 660 (25) | | 30 | | STED, 3 Line accu. | |
| **S21C** | MaP620 | 561 (5) | 584 – 713 | - | 15 | 60 | 60 x 60 | CLSM, 3 Line accu. | **A** |
| **S21D** | | 561 (5) | | - | 15 | 40 | 15 x 15 | CLSM, 3 Line accu. | |
| | | 561 (18) | | 775 (30) | | | | STED, 3 Line accu. | |

Microscopes: **A.** Abberior STED Expert Line 595/775/RESOLFT QUAD scanning microscope (used excitation lines: 485 nm, 561 nm, 640 nm. STED lines: 775 nm), UPlanSApo 100x/1.4 oil immersion objective lens.

**B.** Abberior STED Infinite line 660/775 QUAD scanning microscope (used excitation lines: 561 nm, 640 nm, STED lines: 655 nm and 775 nm) equipped with a 60x/1.42 UPLXAPO60XO oil immersion objective lens.

**Table SI2:** Used dyes and probes for labelling:

| Probe | Figure | Source |
|---|---|---|
| $JF_{585}$-S5 | 3E, S16 | Synthesized like previously described[4] |
| SiR-HTL | S15 | Synthesized like previously described[5] |
| PKMO | S16 | Spirochrome AG |
| SPY650-Fastact | 5 | Spirochrome AG |
| SiR-Actin | | Spirochrome AG |

### 5 Supplementary Schemes

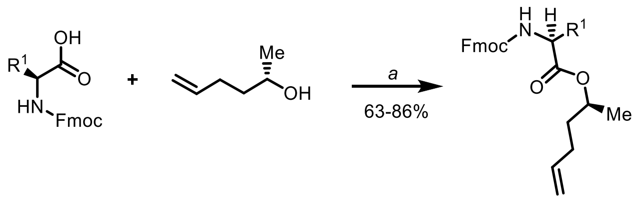

**13-16** **17** **19-22**

| *entry* | *amino acid* | *R¹* | *product* | *yield* |
|---|---|---|---|---|
| 1 | **13** | 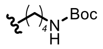 | **19** | 86% |
| 2 | **14** | 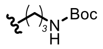 | **20** | 85% |
| 3 | **15** | 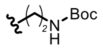 | **21** | 63% |
| 4 | **16** | 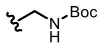 | **22** | 65% |

**Scheme S1.** Synthesis of esters**:** (a) **17** (1.5 equiv.), DCC (1.05 equiv.), $CH_2Cl_2$, 0 °C to 25 °C, 16 h.

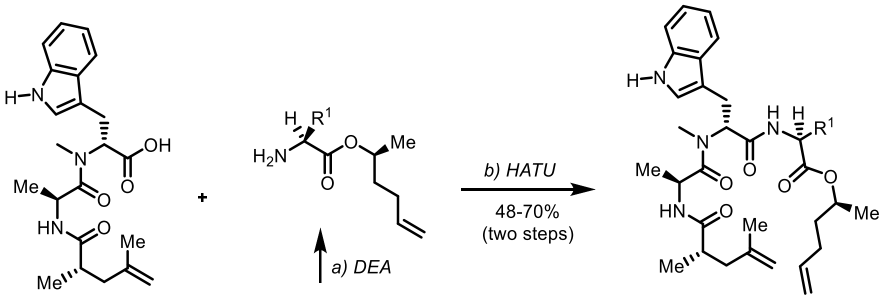

**18** **19-22** **23-26**

| *entry* | *ester* | *R¹* | *product* | *yield* |
|---|---|---|---|---|
| 1 | **19** | 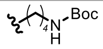 | **23** | 50% |
| 2 | **20** | 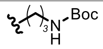 | **24** | 48% |
| 3 | **21** | 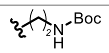 | **25** | 58% |
| 4 | **22** | 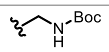 | **26** | 70% |

**Scheme S2.** Synthesis of linear diene precursors: (a) 50% Diethylamine in $CH_2Cl_2$, 0 °C, 2 h. (b) **di-peptide** (0.8 equiv.), HATU (0.8 equiv.), $EtN(i\text{-}Pr)_2$ (1.6 equiv.) THF/DMF then deprotected ester from (a), 16 h, 25 °C.

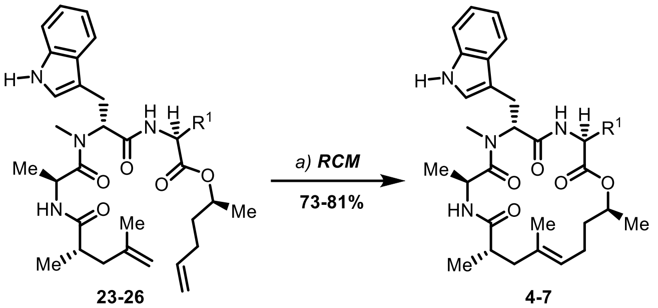

**23-26** **4-7**

| entry | $R^4$ | product | a: yield (RCM) |
|---|---|---|---|
| 1 | 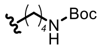 | **7** | 78% |
| 2 | 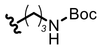 | **6** | 81% |
| 3 | 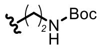 | **5** | 73% |
| 4 | 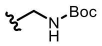 | **4** | 75% |

**Scheme S3. Synthesis of macrocycles by ring-closing metathesis:** (a) RCM*:* Grubbs catalyst 2nd generation (7 mol%), toluene, 1.2 mM concentration, reflux with constant purge of Ar, 2 h.

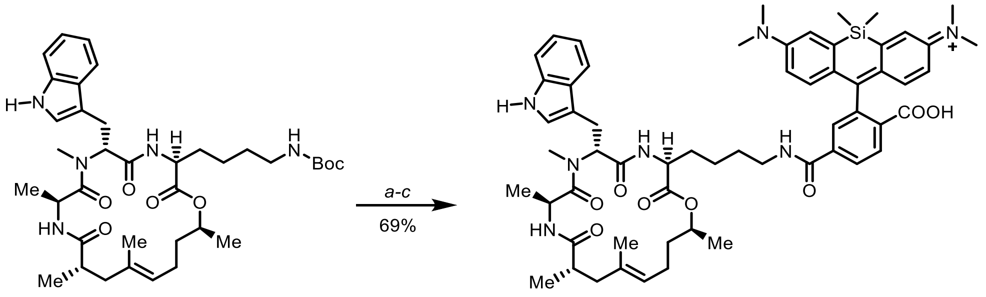

**7** **8**

**Scheme S4. Synthesis of SiR-XActin:** (a) Deprotection: TFA/$CH_2Cl_2$, 0 °C, 15 min. (b) Active ester formation: SiR-6-COOH **27** (1.1 equiv.), PyAOP (1.2 equiv.), EtN(*i*-Pr)$_2$ (4 equiv.), DMF, 25 °C, 5-10 min. (c) Deprotected **7** and activated **27**, 0 °C to 25 °C, 16 h, preparative-HPLC.

### 6 Methods

#### 6.1 General cell culture

**For imaging experiments** HeLa Kyoto FlpIN cells, which were kind gift of Dr. Amparo Andres-Pons (EMBL, Heidelberg), U-2 OS Flp-In T-Rex,[6] COS-7 (Merck, 87021302, African green monkey kidney, Fibroblast) cells were grown in high-glucose Dulbecco's Modified Eagle Medium (DMEM GlutaMAX™, phenol-red, Gibco) supplemented with 10% (v/v) fetal calf serum (FCS, Gibco). Flp-In™-3T3 Cell Line (Thermo Fisher Scientific, R761-07) was grown in high-glucose Dulbecco's Modified Eagle Medium (DMEM GlutaMAX™, no phenol red, Gibco) supplemented with 10% (v/v) newborn calf serum (NBCS) (FCS, Gibco). Passaging of cells was performed every 3 days after washing with phosphate buffered saline (PBS, pH 7.4, Gibco) and incubating with TrypLE™ Select Enzyme (1x, phenol-red free, Gibco). Regular tests for mycoplasma contamination were performed.

**For wound healing assay:** HeLa Kyoto cells were cultured in Dulbecco's modified Eagle's medium (DMEM, Gibco) with 1 g/L D-glucose supplemented with 10% foetal bovine serum (FBS, Invitrogen) and 1% penicillin/streptomycin (Pen/Strep, Invitrogen). MDCKII wild type (Madin-Darby canine kidney, WT) were cultured in DMEM GlutaMAX™ with 5% FBS and 1% Pen/Strep. All cell lines were cultured in 75 $cm^2$ flaks under standard cell culture conditions (37 °C, 5% $CO_2$).

**For viability assay:** HeLa (ACC-57, DSMZ) and MCF-7 (ACC-115, DSMZ) were grown in Dulbecco's modified Eagle`s medium high glucose (Gibco DMEM high glucose, Thermo Fisher Scientific 11965092) and supplemented with 10% heat inactivated fetal bovine serum (Gibco FBS, Thermo Fisher Scientific 11550356), 1% penicillin streptomycin, (gibco Penicillin-Streptomycin 10000IE/mL, Thermo Fisher Scientific 11548876) and 1% L-glutamin (200mM L-Glutamin, Sigma G7513-100ML). Cells were grown at 37°C in a 5% CO2 atmosphere.

##### Transfection

Plasmids for transfection of HeLa cells were gifted from Michael Davidson: mEmerald-Paxilin (Addgene plasmid #54219; http://n2t.net/addgene:54219; RRID: Addgene_54219) and mCherry-Vinculin (Addgene plasmid #55159; http://n2t.net/addgene:55159; RRID: Addgene_55159). Transient transfection was performed using the Lipofectamine 3000® reagent (ThermoFisher Scientific) according to the manufacturer's protocol.

##### Neuron preparation

Primary rat neurons were isolated from new born pups (0-1 days, WISTAR rats) in accordance with the Animal Welfare Act of the Federal Republic of Germany (Tierschutzgesetz der Bundesrepublik Deutschland, TierSchG) and the Animal Welfare Laboratory Animal Regulations (Tierschutzversuchsverordnung). The procedure for euthanizing rats performed in this study was supervised by animal welfare officers of the Max Planck Institute for Medical Research (MPImF) and conducted and documented according to the guidelines of the TierSchG (permit number assigned by the MPImF: MPI/T-35/18). The hippocampi were isolated by trypsin digestion and mechanical dissection using a pipette. The homogenous solution was filtered through a cell strainer (40 µm pore size) and transferred to pre-coated (poly-L-ornithine 100 µg/mL, laminin 1 µg/mL in 1x HBSS) µ-Slide 8-well glass-bottom plates (80826, Ibidi, Gräfelfing, Germany). 2 h post-seeding, the medium was replaced with fresh phenol-red free Neurobasal (NB) medium supplemented with Penicillin/Streptomycin (Life Technologies), GlutaMAX, and B27. All cells were maintained in a humidified tissue culture incubator at 37 °C with 5% $CO_2$

##### rAAV production and transduction

Recombinant AAVs (rAAVs) were produced according to previous protocols.[7] Plasmids pRV1, pH21, pFD6 and the AAV plasmid containing the gene of interest flanked by AAV2 packaging signals (ITRs) were co-transfected via PEI25000 into HEK293 cells. The cells were harvested 5 days post transfection, lysed using TNT extraction buffer (20 mM Tris pH7.5, 150 mM NaCl, 1% TX-100, 10 mM $MgCl_2$) and the cell debris was removed by centrifugation. The cell's supernatant was collected and treated with Benzonase before it was subjected to FPLC purification using AVB Sepharose columns. Subsequently the purified AAVs were concentrated using Amicon centrifugal filters (100 kDa MWCO, Merck KGaA, Darmstadt, Germany). Finally, the buffer was exchanged to PBS pH 7.3.

HeLa cells were seeded in µ-Slide 8-well glass-bottom plates and transduced with ~ $10^9$ -$10^{10}$ rAAV particles phenol-red free DMEM GlutaMAX™ the next day. Imaging experiments were performed 16 h after transduction. The CAG promotor was used for transgene expression. Hippocampal neurons were transduced after 7 days in culture with ~ $10^9$ -$10^{10}$ rAAV particles in phenol-red free NB medium and imaged after 12 days in vitro (DIV). The hSyn promotor was used for transgene expression.

#### 6.2 Imaging

**CLSM (confocal laser-scanning microscopy) and Widefield**

For imaging blebbing phenotype (Figure SI8 and SI9) brightfield mode of DMi8 widefield microscope (Leica) equipped with a HC PL APO 20x/0.8 (dry) was used. For CLSM in all figures except figures containing STED data (see later in text) and when otherwise indicated, Stellaris 5 inverted microscope (Leica) equipped with a white line laser and hybrid photodetectors at 37°C in 5% $CO_2$ atmosphere in a humidified chamber was used. For imaging shown in Figure S19 a confocal Leica SP8 microscope equipped with a Leica TCS SP8 X scan head; a SuperK white light laser, a 355 nm CW laser (Coherent) was used. Both confocal setups are equipped with HC PL FLUOTAR 5x/0.15 DRY, HC PL APO CS2 20x/0.75 (water immersion) objective, HC PL APO CS2 63x/1.40 (oil immersion) objective. Unless otherwise stated, two Line Average was used for all images.
Red Hot lookup from Fiji was used for 16-bit images, unless otherwise stated.

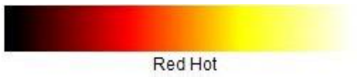

**Settings used in CLSM (unless otherwise stated under respective figure):**

| Probe | Excitation | Emission |
|---|---|---|
| **MaP500-XActin** | 500 nm | 510 nm – 739 nm |
| **MaP555-XActin** | 555 nm | 565 nm – 685 nm |
| **MaP620-XActin** | 610 nm | 620 nm – 750 nm |
| **SiR-XActin** | 652 nm | 662 nm – 829 nm |
| **SiR700-XActin** | 685 nm | 696 nm – 829 nm |

**STED**

Live cells CLSM and STED nanoscopy was performed on A) a Abberior STED Expert Line 595/775/RESOLFT QUAD scanning microscope (used excitation lines: 485 nm, 561 nm, 640 nm; STED lines: 775 nm) equipped with a UPlanSApo 100x/1.4 oil immersion objective lens or B) a Abberior STED Infinite line 660/775 QUAD scanning microscope (used excitation lines: 561 nm, 640 nm, STED lines: 655 nm and 775 nm) equipped with a 60x/1.42 UPLXAPO60XO oil immersion objective lens (both Abberior Instruments). Both systems were equipped with avalanche photodiodes (APD) for signal detection. Please see Supplementary Table 1 for imaging details. Pixel intensities were scaled between dark and bright colours using the ‘Red Hot’, ‘Cyan Hot’ ‘Magenta Hot’ or ‘Orange Hot’ lookup tables (Fiji version 1.54f) for data representation. The min. and max. pixel values are given in the respective reference bars or caption.

For time-lapsed STED microscopy, a 10 x 10 µm frame was imaged repetitively. The mean pixel values over the image series from at least 3 image series was extracted (excluding the boarders) using Fiji. To generate bleaching curves, the data was background corrected and normalized to the first frame intensity. Average bleaching curve and individual traces are shown.

#### 6.3 Fluorescence excitation and emission spectra

XActin derivatives coupled to different fluorophores were diluted from 1 mM DMSO stocks to 1µM in 50 mM HEPES, 50 mM NaCl, 0.1% SDS, pH 7.3. Fluorescence emission and excitation spectra were recorded in black flat bottom 96 well plate (Greiner, 150 µL) with a step size of 1 nm on a microplate reader (Spark20M, Tecan). The detection or excitation settings are summarized in Method Table 2. For representation, the fluorescence excitation or emission was normalized to the max. intensity measured for each dye. The data from three technical replicates were averaged.

**Settings used to record fluorescence excitation and emission spectra:**

| Probe | Excitation wavelength [nm] | Emission wavelength [nm] |
|---|---|---|
| **MaP500-XActin** | 580±10 | 460±10 |
| **MaP555-XActin** | 620±10 | 530±10 |
| **MaP620-XActin** | 650±10 | 610±10 |
| **SiR-XActin** | 715±10 | 590±10 |
| **SiR700-XActin** | 765±10 | 650±10 |

#### 6.4 Cytotoxicity profiling

##### MTT viability assay

Mitochondrial dehydrogenase activity in HeLa and MCF-7 cells was quantified spectrophotometrically measuring the reduction of 3-(4,5-dimethylthiazol-2-yl)-2,5-diphenyl tetrazolium bromide (MTT) to formazan, according to standard protocols. Cells were seeded in 96-well microtiter plates at cell line dependent density (HeLa: 7500 cells/well, MCF-7 8500 cells/well). After 24 h cells were treated with the test compounds. In the MTT experiments 1% DMSO was present as cosolvent for all compounds at all concentrations tested, to ensure full solubility of all compounds and comparability of the data. Under these conditions control cells were growing unaffected. Following 48 h of treatment, cells were incubated with MTT in cell line dependent concentration and time (HeLa: 0.5 mg/mL, 2.5 h, MCF-7 0.3 mg/mL, 5.5 h). Formazan crystals were then dissolved in isopropanol with 0.1N HCl and 10% Triton-X. Formazan absorbance was measured at 565 nm and background absorption at 800 nm to avoid absorption of the chromophore with a TECAN microplate Reader M200Pro (band width: 9 nM, flashes: 22). For statistical analysis the absorbance readings for untreated controls (cosolvent DMSO only) were set as 100% viability of the cells. Results are given as the mean percentage of viable cells relative to these controls ± standard deviation (SD) from representative experiment out of three independent trials performed in quadruplets. Graphical representations and statistical analysis of the data were performed using GraphPad Prism (GraphPad Software, La Jolla, CA, USA). Variable slope fitting for non-linear regression fit with dose-response curve: inhibition was used calculating a symmetrical (asymptotic) 95% confidence interval for the fits and showing SE of the parameters.

#### 6.5 Wound healing assay

**Cell migration imaging and F-actin staining**. To evaluate cell migration, a wound healing assay was performed using 3-well culture inserts (ibidi, Germany) adhered to surfaces coated with 10 µg $ml^{-1}$ fibronectin. HeLa Kyoto cells were seeded at a concentration of 50000 cells/insert and MDCKII WT were seeded at 30000 cells/insert. After 2 h incubation, each culture medium was replaced containing fluorescent F-actin dyes, silicon rhodamine SiR-actin (Spirochrome) or SiR-Xactin, at a final concentration of 0.5 µM. For MDCKII WT, 10 µM verapamil was added to the staining culture medium. After a confluent monolayer was obtained for each cell line after overnight incubation, culture inserts were removed to trigger cell migration and live cell images were recorded every 10 min for 24 h in culture media supplemented with 1% FBS (Axiovert Zeiss, Germany). A 1:2000 dilution of dimethylsulfoxide (DMSO, Invitrogen) in culture medium was used as control.

**Monolayer migration analysis**. Timelapse images were analyzed using particle image velocimetry (PIV). To that end, the MATLAB (The MathWorks, Natick, MA) software package MatPIV *(Sveen, 2014)* (MatPIV 1.6.1, freely available as a GNU general public license) was modified to suit the experimental setup. Interrogation window size was set to 32 × 32 pixels with 50% overlap. From velocity fields obtained from PIV analysis, kymographs depicting the dynamics of the cell monolayers over time were plotted and, the linear order parameter and the persistence length were derived.[8]

#### 6.6 Molecular docking

In-silico docking was perforemd using the Schrödinger Maestro 12.3 software package. SiR-Xactin was sketched with the 2D Sketcher module in Maesttro. Ligand was preprocessed in the Schroedinger ***ligprep*** module and most probable ionization states (Epik module, pH 7.4 ± 2.0). Cryo-EM structure of jasplakinolide in actin fibre (PDB 6T24) was at first simplified by removal of 2 G-actin monomers and 4 jasplakinolide ligands, resulting trimer of G-actin was prepared in the Prime module. Default settings (pH value 7.4) were used. A cubic docking grid was generated around jasplakinolide ligand molecule and subsequent docking was performed using Glide2 in standard precision configuration yielding up to 10 docking poses per ligand. During the docking process, no interactions were applied as constrains. Maestro was used for inspection of probable protein/ligand poses and PyMol Schrödinger 2.5.2 was used for graphics preparation and for overlay with jasplakinolide ligand in the binding cleft.

#### 6.7 Binding affinity determination

Binding affinity determination was performed similar like previously described, with minor adjustments.[1] U-2 OS cells (10 000) were seeded in Black 96 well culture plate with square wells and a clear bottom for high-throughput microscopy and grown overnight under cell culture conditions described above. Medium was aspirated and cells were washed 2x with PBS pH 7.4 after which cells were incubated at 37 °C with 4% PFA in PBS for 15 min. Next, cells were washed with PBS (3x) and with BSA (1% in PBS, 1x). Dose response dilutions of SiR-actin and SiR-XActin were prepared in PBS with 1% of BSA and cells were incubated for 16 h prior to measurement at room temperature. Each condition was performed in technical triplicate. Stellaris 5 equipped with HC PL FLUOTAR 5x/0.15 DRY was used (like above described) with 2% laser power, line average 2, image size 3,100 µm x 3,100 µm, Pixel Size 1.514 µm. Images were processed using ImageJ 1.54f, where median of fluorescence signal intensity was measured and technical triplicates were used for a sigmoidal dose response curve fit by GraphPad Prism to obtain apparent binding affinity values $Kd_{app}^{F\text{-}actin}$. Average values of biological independent replicates are shown at Figure SI5.

### 7 Synthesis

#### 7.1 General information

All reagents were obtained from Acros Chemicals, Alfa Aesar, Apollo Scientific, ABCR, Carbolution Chemicals, Carbosynth, Manchester Organics, Merck, Novabiochem, Sigma-Aldrich, TCI Europe, or VWR, and used without further purification. All solvents, if not purchased in purity or dryness suitable, were distilled using standard methods.[9] Tetrahydrofuran (THF) was distilled under an $N_2$ atmosphere from Na/benzophenone; dichloromethane ($CH_2Cl_2$) was distilled under an $N_2$ atmosphere from $CaH_2$ before use. Other solvents were passed through alumina columns (toluene) or molecular sieves columns (dimethylformamide - DMF) of a solvent purification system (Pure Solv, Innovative Technology, Inc., USA) by applying $N_2$ overpressure immediately before use.

All solvents for flash chromatography were distilled before use. All solvents used in reactions were anhydrous. Solvents were degassed by employing triple freeze-pump-thaw cycles when necessary or by purging with $N_2$ for a minimum of 15 minutes. Deionized water was used for all experiments. All reactions were performed in heat-dried glassware under an atmosphere of $N_2$ if not stated otherwise. TLC was carried out on Merck precoated silica gel plates (60F-254); compounds were visualized using ultraviolet light irradiation at 254 nm and 366 nm or by using the following staining agents (dip, dry & heat development): Potassium Permanganate: $KMnO_4$ (1 g), $K_2CO_3$ (6.6 g), 5% NaOH (1.7 mL) in $H_2O$ (90 mL); Ninhydrin: Ninhydrin (0.3 g) dissolved in *n*-butanol (100 mL) and acetic acid (3 mL).

All synthetic products were either purified *via* normal phase flash column chromatography (FCC) manually (Purifications were performed using silica gel from Macherey & Nagel (particle size 40-60 µm) under approximately 0.2-0.6 bar pressure)[10] or using an automated system (Biotage Isolera One) with pre-packed silica gel columns (ultrapure silica gel 12 g or 25 g, SiliCycle Inc.) or *via* reverse-phase high-performance liquid semipreparative chromatography (RP-HPLC) on a Thermo Fisher Scientific UltiMate 3000 system with a Supelco column (21.1 × 250 mm, 5 µm pore size, 8 mL/min flow rate) using a solvent gradient of 10-95% MeCN/$H_2O$ with constant 0.1% v/v trifluoroacetic acid additive (a standard RP-HPLC purification run took 45 min). For larger scales preparative RP-HPLC was used, a Varian Prostar system equipped with a fraction collector Prostar 701 and detection at 220 nm (UV/Vis Prostar 340) and a VP 250/21 Nucleodur C18 Gravity 5 µm column (Macherey & Nagel, Düren, Germany). *Method used for preparative HPLC:* isocratic: 2 min (30% MeCN in $H_2O$), → gradient over 38 min (30-100% MeCN in $H_2O$) → 10 min 100% MeCN. Flow 25 mL/min.

Fractions containing the product were combined and concentrated *in vacuo* (rotary evaporator, Hei-Vap value, heidolph) with heating in a water bath (40 °C) and/or dried on a lyophilizer (Christ Alpha 1-2 LDplus) connected to a vacuum pump (Vacuubrand). Final fluorophore conjugates were stored as DMSO stocks at -20 °C.

Samples for nuclear magnetic resonance (NMR) spectroscopy were dissolved in deuterated solvents and NMR spectra were recorded at 298 K on a Bruker Advance III HD 400 NMR spectrometer equipped with a CryoProbe™ ($^1$H: 400 MHz, $^{13}$C: 101 MHz), Bruker Avance I 250 system (250 MHz for $^1$H- and 63 MHz for $^{13}$C-NMR), a Bruker Fourier 300 system (300 MHz, $^1$H- and 75 MHz for $^{13}$C-NMR), a Bruker Avance I 400 system (400 MHz for $^1$H- and 101 MHz for $^{13}$C-NMR), Bruker Avance 600 system (600 MHz for $^1$H- and 151 MHz for $^{13}$C-NMR) or Avance III HD [500 MHz, probe: BBO (Prodigy)] . NMR spectra were analysed using MestReNova 14.1.0 (MestreLab Research). Multiplicities are reported as s = singlet, d = doublet, t = triplet, q = quartet, m = multiplet and chemical shifts (δ) are calibrated to the residual chemical shifts of the solvents ($CDCl_3$, MeOD-$d_4$, MeCN-$d_3$, DMSO-$d_6$, DMF-$d_7$)[11]. Coupling constants J are reported in Hz. Spectra were recorded at 298 K, if not stated otherwise. Chloroform-*d* was stored over molecular sieves at 5 °C. If mixtures of solvents were used, namely chloroform-*d* and methanol-$d_4$, spectra were calibrated to chloroform-$d_1$.

High-resolution mass spectrometry (HRMS) was acquired on a maXis II™ ETD-HRMS system (Bruker) using electron spray ionization (ESI) in positive mode conducted by the MS Core Facility (Max Planck for Medical Research, Heidelberg) or LC-coupled MAXIS Impact ESI-TOF spectrometer (Bruker Daltronics, Bremen, Germany) at the University of Jena.

IR (ATR): spectra were measured by using a Thermo Nicolet Spectrometer FT-IR (ATR): Avatar 370 fitted with an ATR unit. Spectra were analysed using Spectragryph v1.2.11. The following notations indicate the normalized intensity of the absorption bands: s = strong (intensity > 67%), m = medium (intensity: 34%-66%), w = weak (intensity < 34%). Normalization was performed in Microsoft Excel.

For melting point determination, a Büchi B-545 melting point apparatus and one-side open capillaries were used. All given values are an average of three measurements.

For specific optical rotation - optical rotations were recorded in a Jasco P-2000 polarimeter at 589 nm and at a given temperature 22-24 °C. The path length of cuvettes was *d = 10 mm*. Concentrations are given in g/100 mL solvent if not stated otherwise.

#### 7.2 Synthetic procedures

##### 7.2.1 General procedures

###### 7.2.1.1 General procedure 1: Esterification of amino acids

Amounts given for 2 mmol of respected protected amino acid. Amino acid (2 mmol) was dissolved in 50 mL of $CH_2Cl_2$, followed by the addition of alcohol **17** (3 mmol, 1.5 equiv. to respected acid, if other not noted) and DMAP (0.4 mmol, 0.2 equiv.). The mixture was cooled to -20 °C, and DCC (2.1 mmol, 1.05 equv.) was added in one portion. The reaction mixture was stirred for 16 h, allowing to reach 25 °C. The mixture was filtered, diluted with $CH_2Cl_2$ (~40 mL) and washed with a saturated aqueous solution of $NH_4Cl$ (~30 mL) and brine (~30 mL). The organic layer was dehydrated ($Na_2SO_4$), concentrated under reduced pressure, and the residue was purified by FCC.

###### 7.2.1.2 General procedure 2: Fmoc-Deprotection in solution

Fmoc-carbamate (0.1 mmol). Amino acid ester (0.1 mmol) was dissolved in 1 mL of $CH_2Cl_2$, the solution was cooled to 0 °C, and diethylamine was added in one portion (1 mL). The reaction mixture was followed by TLC, and after completion (average reaction time 3 h), toluene (3 mL) was added. Volatiles were removed under reduced pressure using a rotary evaporator (water bath at 25 °C) followed by drying under a high vacuum for 2-3 h. The residue was used in the coupling reaction without further purification.

###### 7.2.1.3 General procedure 3: Coupling of peptide fragments in solution

The peptide acid (0.08 mmol, 0.8 equiv.) was mixed with HATU (0.08 mmol) and EtN(*i*-Pr)$_2$ (0.24 mmol, 3 equiv.) in 5 mL of THF/DMF (1:1) at 25 °C and stirred for 3 min. The active ester reaction mixture was added to 0.1 mmol (1 equiv.) of deprotected ester (described in **General procedure 2**, and the reaction mixture was stirred for 16 h. The reaction mixture was diluted with EtOAc (~20 mL), washed with water (2 x ~10 mL), brine (2 x ~10 mL), and the organic layer was dehydrated ($Na_2SO_4$). Volatiles were evaporated under reduced pressure, and the product was purified by FCC.

###### 7.2.1.4 General procedure 4: Ring-closing metathesis

Toluene in two neck flask was heated to reflux and active purge with a argon for 30 min. The linear diene precursor (0.1 mmol, 1 equiv.) for metathesis was dissolved in a small amount of degassed $CH_2Cl_2$ and added to the degassed toluene at a with toluene, final concentration of ~1.2 mM. The solution was heated to reflux with continuous purging of argon for 30 min. Grubbs catalyst 2$^{nd}$ generation (7.5 mol%) was added at reflux. Reactions were carried out until LC/MS or TLC controlled showed full conversion (1-3 h) with a constant purge of argon at reflux. Reactions were followed by TLC. After completion, the solvent was evaporated under reduced pressure, and products were purified using FCC. For direct use in biological assays prep. HPLC (Method A) purification was used.

##### 7.2.2 Building blocks synthesized by previously reported literature

*N*$^α$-Fmoc-*N*$^α$-Me-*D*-Tryptophan,[12] (*S*)-2,4-dimethylpent-4-enoic acid,[12] MAP500-COOH (**31**),[13] MAP555-COOH (**28**),[14] SiR-6-COOH (**27**),[15], CPY-6-*O*-allyl (**32**),[13] SiR700-6-COOH (**30**).[16]

##### 7.2.3 Synthesis of amino acid esters esters

**(2*S*)-2-[Fmoc-L-Lys(Boc)-*O*]-hex-5-en (19):**

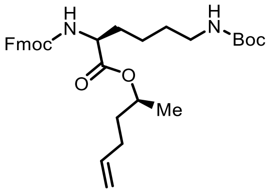

Ester **19** was synthesized following **General procedure 1**. Purification was performed by automated FCC (gradient of PE/EtOAc 9:1 to 7:3). Ester **19** was obtained as a colorless glass in a yield of 86% (2.05 g).

**TLC:** $R_f$ **=** 0.45 (PE/EtOAc = 9:1).

$[\alpha]_D^{22}$ **= -**1.3 (c = 1, $CHCl_3$/MeOH = 1:1).

**IR (ATR):** $\tilde{\nu}$ **=** 2935 (w), 2866 (w), 2488 (w), 1612 (w), 1523 (m), 1365 (m), 1338 (s), 1292 (m), 1249 (m), 1211 (m), 1126 (m), 1107 (m), 1087 (s), 1053 (m), 1033 (m), 995 (m), 910 (m), 864 (m), 775 (m), 756 (s), 732 (s), 644 (m), 621 (m) $cm^{-1}$.

**$^{1}$H NMR** (400 MHz, chloroform-*d*, 297 K): $\delta$ = 7.74 (t, *J* = 13.4 Hz, 2H), 7.61 (d, *J* = 6.0 Hz, 2H), 7.39 (dd, *J* = 15.1, 7.8 Hz, 2H), 7.26 (s, 2H), 5.89 – 5.65 (m, 1H), 5.39 (d, *J* = 58.2 Hz, 1H), 5.00 (t, *J* = 13.6 Hz, 3H), 4.75 – 3.97 (m, 5H), 3.12 (s, 2H), 2.07 (s, 2H), 1.83 (t, *J* = 14.6 Hz, 1H), 1.79 – 1.53 (m, 4H), 1.44 (s, 10H), 1.26 (d, *J* = 5.3 Hz, 3H) ppm.

**$^{13}$C{$^{1}$H} NMR** (101 MHz, chloroform-*d*, 297 K): $\delta$ = 172.1, 156.1, 156.1, 144.0, 143.9, 141.4, 137.5, 127.8, 127.2, 125.2, 120.1, 115.4, 72.1, 67.1, 54.0, 47.3, 40.2, 34.9, 32.4, 29.7, 29.6, 28.5, 22.5, 19.9 ppm.

**HRMS** (ESI−TOF) *m/Z*: [M + H]$^{+}$ calculated for $C_{32}H_{43}N_{2}O_{6}^{+}$ 551.3116; found: 551.3117.

**(2*S*)-2-[Fmoc-L-Orn(Boc)-*O*]-hex-5-en (20):**

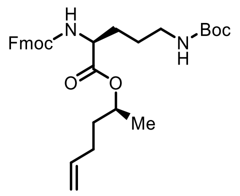

Ester **20** was synthesized following **General procedure 1**. Purification was performed by FCC (PE/EtOAc 85:14). Ester **20** was obtained as a colorless solid, in a yield of 85% (938 mg).

**TLC:** $R_f$ **=** 0.54 (PE/EtOAc = 7:3).

**$[\alpha]_D^{22}$ =** -2.3 (c = 1, $CHCl_3$/MeOH = 1:1).

**IR (ATR):** $\tilde{\nu}$ **=** 3352 (m), 3069 (w), 2975 (m), 2937 (m), 2879 (w), 1749 (m), 1684 (s), 1519 (s), 1449 (m), 1367 (m), 1283 (s), 1242 (s), 1218 (m), 1167 (s), 1102 (m), 1026 (s), 909 (m), 868 (m), 782 (m), 735 (s), 643 (m) $cm^{-1}$.

**MP =** 106 °C.

**$^{1}$H NMR** (400 MHz, chloroform-*d*, 297 K): $\delta$ = 7.79 (d, *J* = 7.5 Hz, 2H), 7.63 (d, *J* = 7.2 Hz, 2H), 7.46 – 7.30 (m, 4H), 5.87 – 5.69 (m, 1H), 5.47 (s, 1H), 5.15 – 4.89 (m, 3H), 4.43 (d, *J* = 7.1 Hz, 4H), 4.25 (t, *J* = 6.9 Hz, 1H), 4.10 – 3.91 (m, 1H), 3.17 (d, *J* = 5.9 Hz, 2H), 2.09 (dd, *J* = 12.1, 4.9 Hz, 2H), 1.92 (dd, *J* = 8.5, 4.8 Hz, 1H), 1.81 – 1.51 (m, 5H), 1.47 (s, 9H), 1.12 (dd, *J* = 125.1, 6.3 Hz, 3H) ppm.

**$^{13}C\{^1H\}$ NMR** (101 MHz, chloroform-*d*, 297 K): $\delta$ = 171.9, 155.9, 143.9, 143.8, 141.3, 137.4, 135.8, 127.7, 127.1, 125.1, 120.0, 119.9, 116.8, 115.3, 79.3, 77.2, 72.1, 69.8, 66.9, 53.9, 47.2, 40.0, 37.7, 34.9, 33.9, 32.4, 30.1, 29.6, 28.4, 26.1, 19.9, 16.6 ppm.

**HRMS** (ESI−TOF) *m/Z*: $[M + H]^+$ calculated for $C_{31}H_{41}N_2O_6^+$ 537.2959; found: 537.2967.

**(2*S*)-2-[Fmoc-L-Dab(Boc)-*O*]-hex-5-en (21):**

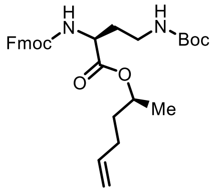

Ester **21** was synthesized following **General procedure 1**. Purification was performed by automated FCC (PE/EtOAc 9:1 to 7:3). Ester **21** was obtained as colorless solid, in a yield of 63% (118 mg).

**TLC: $R_f$ =** 0.45 (PE/EtOAc = 7:3).

**$[\alpha]_D^{22}$ =** -12.8 (c = 0.1, $CHCl_3$/MeOH = 1:1).

**IR (ATR):** $\tilde{\nu}$ **=** 2975 (w), 2933 (w), 2189 (w), 2164 (w), 2112 (w), 2084 (w), 2056 (w), 1955 (w), 1730 (m), 1684 (m), 1523 (m), 1449 (m), 1390 (w), 1251 (m), 1167 (m), 1106 (m), 1086 (m), 1039 (m), 1007 (m), 912 (m), 867 (w), 736 (m), 619 (m) $cm^{-1}$.

**MP =** 72 °C.

**$^1H$ NMR** (300 MHz, chloroform-*d*, 298 K): $\delta$ = 7.78 (d, *J* = 7.6 Hz, 2H), 7.62 (d, *J* = 7.3 Hz, 2H), 7.42 (s, 2H), 7.33 (s, 2H), 5.89 – 5.68 (m, 1H), 5.57 (d, *J* = 7.1 Hz, 1H), 5.20 – 4.90 (m, 4H), 4.43 (t, *J* = 7.0 Hz, 3H), 4.34 – 3.85 (m, 2H), 3.44 (s, 1H), 3.14 – 2.84 (m, 1H), 2.08 (d, *J* = 5.0 Hz, 3H), 1.86 – 1.53 (m, 3H), 1.46 (s, 9H), 1.27 (d, *J* = 6.3 Hz, 3H), 0.95 (d, *J* = 6.5 Hz, 1H) ppm.

**$^{13}C\{^1H\}$ NMR** (75 MHz, chloroform-*d*, 298 K): $\delta$ = 171.9, 156.4, 155.9, 143.8, 143.6, 141.3, 137.3, 135.8, 127.7, 127.1, 125.1, 125.0, 120.0, 119.9, 116.8, 115.3, 79.4, 72.4, 69.9, 67.1, 51.7, 47.2, 37.6, 36.5, 34.9, 33.6, 32.3, 29.5, 28.4, 19.9, 16.6 ppm.

**HRMS** (ESI−TOF) *m/Z*: $[M + H]^+$ calculated for $C_{30}H_{39}N_2O_6^+$ 523.2803; found: 523.2805.

**(2*S*)-2-[Fmoc-L-Dap(Boc)-*O*]-hex-5-en (22):**

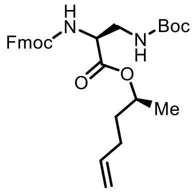

Ester **22** was synthesized following **General procedure 1**. Purification was performed by automated FCC (PE/EtOAc 9:1 to 7:3). Ester **22** was obtained as a colorless glass, in a yield of 65% (156 mg).

**TLC:** $R_f$ **=** 0.40 (PE/EtOAc = 7:3).

$[\alpha]_D^{22}$ **=** 2.0 (c = 0.1, $CHCl_3$/MeOH = 1:1).

**IR (ATR):** $\tilde{\nu}$ **=** 3348 (w), 3160 (w), 3068 (w), 2771 (w), 2319 (w), 1828 (w), 1691 (m), 1518 (m), 1449 (m), 1392 (m), 1366 (m), 1337 (m), 1251 (m), 1200 (m), 1162 (m), 1083 (m), 1036 (m), 993 (m), 950 (m), 912 (m), 861 (m), 737 (m), 620 (m) $cm^{-1}$.

**$^1$H NMR** (300 MHz, chloroform-*d*, 298 K): $\delta$ = 7.78 (d, *J* = 7.5 Hz, 2H), 7.62 (d, *J* = 7.2 Hz, 2H), 7.42 (t, *J* = 7.4 Hz, 2H), 7.32 (t, *J* = 7.4 Hz, 2H), 5.78 (d, *J* = 6.7 Hz, 2H), 4.98 (dt, *J* = 57.1, 23.8 Hz, 4H), 4.40 (s, 3H), 4.24 (t, *J* = 7.0 Hz, 2H), 3.58 (s, 2H), 2.24 – 1.88 (m, 2H), 1.68 (d, *J* = 44.0 Hz, 3H), 1.46 (s, 9H), 1.27 (d, *J* = 6.2 Hz, 3H), 0.95 (d, *J* = 6.3 Hz, 1H) ppm.

**$^{13}$C{$^1$H} NMR** (75 MHz, chloroform-*d*, 298 K): $\delta$ = 169.9, 156.4, 143.7, 141.3, 137.4, 127.7, 127.1, 125.2, 119.9, 115.3, 80.1, 72.6, 67.2, 55.3, 47.1, 42.3, 34.8, 29.5, 28.3, 19.8 ppm.

**HRMS** (ESI−TOF) *m/Z*: $[M + H]^+$ calculated for $C_{29}H_{37}N_2O_6^+$ 509.2646; found: 509.2650.

##### 7.2.3.1 Synthesis of acylated dipeptide

**L-Pea-L-Ala-D-*N*-Me-Trp-OH (18):**

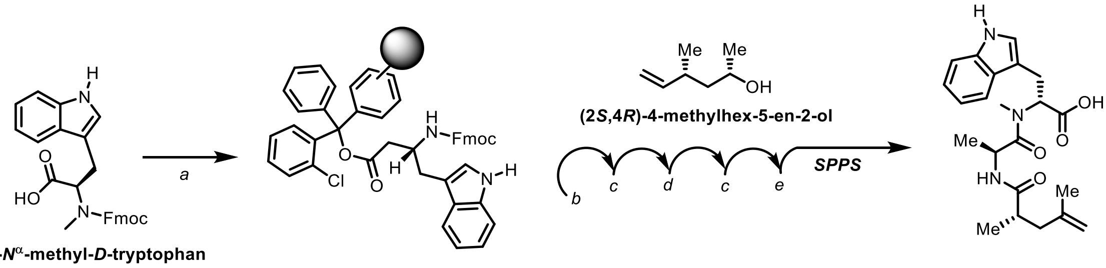

**Comments:** (a) **loading** of *N*α-Fmoc-*N*α-Me-*D*-Tryptophan building block; (b) **deprotection** – piperidine**;** (c) **coupling;** (d) **deprotection piperidine/DBU**; (e) **cleavage from resin**: HFIP.

Synthesis of the building block **18** was performed like previously described for similar building blocks,[17] with small adjustments in the second deprotection step, after coupling of L-Alanine, like described before in total synthesis of Seragamide A.[18]

*N*α-Fmoc-*N*α-Me-*D*-Tryptophan was loaded on the solid support and loading was quantified like previously described (loading 0.54 mmol/g).[17] Amounts of amino acid building blocks used were: 4 equiv. of *N*-Fmoc-L-Alanine, reaction duration 16 h, and 1.2 equiv. of (*S*)-2,4-dimethylpent-4-enoic acid, reaction duration 4 h. After cleaving the synthesized products from the solid support, the peptide **18** was obtained (87%, 1.4 g) by purification using FCC ($CH_2Cl_2$/MeOH 95:5 + 0.1% HCOOH) as a slightly yellow resin.

**TLC: $R_f$ =** 0.42 ($CH_2Cl_2$/MeOH 9:1 + 1% HCOOH).

**$[\alpha]_D^{22}$ =** +6.4 (c = 0.1, $CHCl_3$).

**IR (ATR):** $\tilde{\nu}$ **=** 3309 (w), 2970 (w), 2931 (w), 1716 (m), 1612 (s), 1523 (m), 1489 (m), 1454 (m), 1415 (m), 1373 (m), 1342 (m), 1207 (m), 1099 (m), 1010 (w), 945 (m), 898 (m), 813 (w), 736 (s), 648 (m) $cm^{-1}$.

**$^1$H NMR** (300 MHz, chloroform-*d,* 298 K): *δ* = 10.58 (s, 1H), 8.94 (s, 1H), 7.83 (t, *J* = 10.6 Hz, 1H), 7.68 – 7.29 (m, 4H), 5.48 (dd, *J* = 10.7, 4.4 Hz, 1H), 5.15 – 5.05 (m, 1H), 5.03 (s, 1H), 4.95 (d, *J* = 10.3 Hz, 1H), 3.79 (dd, *J* = 15.0, 4.2 Hz, 1H), 3.67 – 3.51 (m, 1H), 3.25 (s, 1H), 3.11 (s, 3H), 2.84 – 2.71 (m, 1H), 2.64 (dd, *J* = 14.0, 6.9 Hz, 1H), 2.42 – 2.25 (m, 2H), 1.95 (s, 3H), 1.52 (dd, *J* = 15.5, 8.9 Hz, 1H), 1.35 (dd, *J* = 22.4, 6.7 Hz, 3H), 1.18 (d, *J* = 6.7 Hz, 3H) ppm.

$^{13}$**C{$^{1}$H} NMR** (101 MHz, chloroform-*d*, 298 K): $\delta$ = 176.7, 174.0, 173.1, 172.4, 163.32 150.1, 142.8, 136.2, 127.1, 122.9, 122.1, 119.5, 118.2, 112.5, 111.4, 110.5, 59.6, 45.4, 41.7, 39.0, 36.9, 33.9, 25.4, 24.2, 22.2, 17.3, 17.2 ppm.

**HRMS** (ESI−TOF) *m/Z*: $[M + H]^+$, calculated for $C_{22}H_{30}N_3O_4^+$ 400.2231; found 400.2233.

##### 7.2.3.2 Synthesis of linear diene precursors

**(2*S*)-2-[L-Pea-L-Ala-D-*N*-Me-Trp-L-Lys(Boc)-*O*]-hex-5-en (23):**

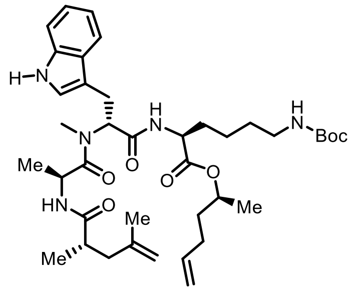

Diene **23** was synthesized by **General procedure 2 and 3** from ester **19** (287 mg) and dipeptide **18** (160 mg). By using FCC (EtOAc/PE 1:1), peptide **23** was obtained as a colorless solid (142 mg, 50% yield).

**TLC: $R_f$ =** 0.18 (PE/EtOAc = 1:1).

**$[\alpha]_D^{24}$ =** -71.4 (c = 0.1, $CHCl_3$/MeOH = 1:1).

**IR (ATR):** $\tilde{\nu}$ **=** 3589 (w), 3309 (w), 3078 (w), 2975 (w), 2934 (w), 2716 (w), 2373 (w), 2294 (w), 2008 (w), 1829 (w), 1638 (m), 1518 (m), 1457 (m), 1366 (m), 1249 (m), 1168 (m), 1009 (w), 914 (w), 842 (s), 740 (m), 601 (m) $cm^{-1}$.

$^{1}$**H NMR** (400 MHz, chloroform-*d*:methanol-$d_4$ = 9:1, 297 K): $\delta$ = 7.55 (d, *J* = 7.8 Hz, 1H), 7.34 – 7.28 (m, 1H), 7.06 (ddd, *J* = 14.9, 14.0, 7.0 Hz, 2H), 6.95 (s, 1H), 5.69 (tdd, *J* = 17.4, 8.7, 5.4 Hz, 1H), 5.43 (d, *J* = 7.6 Hz, 1H), 5.08 – 4.78 (m, 3H), 4.74 – 4.57 (m, 3H), 4.48 – 4.30 (m, 1H), 4.02 – 3.79 (m, 1H), 3.50 – 3.34 (m, 14H), 3.34 – 3.15 (m, 3H), 3.14 – 2.87 (m, 6H), 2.75 (s, 5H), 2.40 (dt, *J* = 13.2, 6.7 Hz, 1H), 2.28 (dd, *J* = 14.1, 6.7 Hz, 1H), 2.13 – 1.68 (m, 5H), 1.66 – 1.59 (m, 4H), 1.59 – 1.45 (m, 2H), 1.43 – 1.35 (m, 10H), 1.27 – 1.01 (m, 8H), 0.99 – 0.78 (m, 4H) ppm.

$^{13}$**C{$^{1}$H} NMR** (101 MHz, chloroform-*d*:methanol-$d_4$ = 9:1, 297 K): $\delta$ = 174.0, 171.8, 170.2, 142.8, 137.3, 136.2, 135.7, 127.1, 122.7, 121.6, 119.0, 118.2, 116.6, 115.1, 112.2, 111.3,

109.8, 79.3, 71.9, 69.6, 56.9, 52.2, 45.5, 41.6, 40.1, 38.6, 38.4, 37.5, 34.8, 32.3, 31.1, 29.4, 29.2, 28.3, 23.9, 22.6, 21.9, 19.6, 17.1, 16.4 ppm.

**HRMS** (ESI−TOF) *m/Z*: [M + H]$^+$ calculated for $C_{39}H_{60}N_5O_7^+$ 710.4487; found: 710.4493.

**(2*S*)-2-[L-Pea-L-Ala-D-*N*-Me-Trp-L-Orn(Boc)-*O*]-hex-5-en (24):**

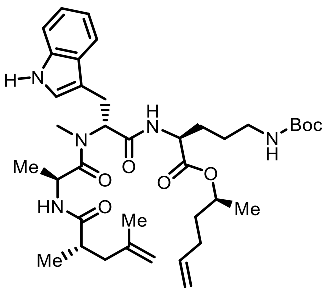

Diene **24** was synthesized by **General procedure 2 and 3** from ester **20** (200 mg) and dipeptide **18** (121 mg). By using FCC (EtOAc/PE 1:1), peptide **24** was obtained as an off-white solid (101 mg, 48% yield).

**TLC:** $\boldsymbol{R_f}$ **=** 0.40 (PE/EtOAc = 3:7).

**[α]$_D^{24}$ =** -36.8 (c = 0.1, $CHCl_3$/MeOH = 1:1).

**IR (ATR):** $\tilde{\nu}$ **=** 3132 (w), 3107 (w), 2953 (m), 2889 (m), 2760 (w), 2515 (w), 2216 (w), 2094 (w), 2071 (w), 2019 (w), 1861 (w), 1585 (m), 1471 (s), 1425 (s), 1383 (s), 1325 (s), 1217 (s), 1114 (s), 1055 (s), 964 (m), 935 (s), 873 (s), 815 (m), 713 (s), 623 (s) cm$^{-1}$.

**$^1$H NMR** (300 MHz, chloroform-*d*:methanol-$d_4$ = 9:1, 298 K): $\delta$ = 8.18 (s, 1H), 7.83 (d, *J* = 7.6 Hz, 1H), 7.60 (d, *J* = 7.9 Hz, 1H), 7.34 (dd, *J* = 14.3, 7.6 Hz, 2H), 6.01 (s, 1H), 5.73 (dd, *J* = 37.2, 25.1 Hz, 2H), 5.37 – 5.11 (m, 3H), 4.99 (s, 1H), 4.93 (s, 1H), 4.82 (d, *J* = 6.7 Hz, 1H), 4.76 – 4.64 (m, 1H), 4.22 (d, *J* = 7.9 Hz, 3H), 3.72 (s, 1H), 3.41 (d, *J* = 10.0 Hz, 1H), 3.19 (s, 2H), 3.09 (s, 2H), 3.04 (s, 4H), 2.72 (s, 1H), 2.56 (s, 1H), 2.28 (s, 3H), 2.10 (d, *J* = 7.2 Hz, 1H), 1.92 (s, 6H), 1.68 (s, 12H), 1.48 (d, *J* = 6.2 Hz, 4H), 1.32 (d, *J* = 6.8 Hz, 4H), 1.17 (d, *J* = 6.7 Hz, 4H) ppm.

**$^{13}$C{$^1$H} NMR** (75 MHz, chloroform-*d*:methanol-$d_4$ = 9:1, 298 K): $\delta$ = 176.7, 174.0, 172.2, 171.6, 170.6, 165.7, 163.0, 156.5, 142.5, 137.1, 136.2, 136.1, 135.6, 126.9, 122.7, 122.5, 121.3, 118.6, 117.9, 116.4, 114.9, 112.0, 111.1, 109.5, 78.9, 71.8, 69.4, 60.3, 56.8, 52.1, 45.6, 45.5, 41.3, 39.3, 38.1, 37.3, 36.3, 34.6, 32.0, 31.1, 30.9, 29.2, 28.1, 28.0, 25.6, 23.6, 21.7, 19.4, 16.7, 16.1, 15.9 ppm.

**HRMS** (ESI−TOF) *m/Z*: [M + H]$^+$ calculated for $C_{38}H_{58}N_5O_7^+$ 696.4331; found: 696.4339.

**(2*S*)-2-[L-Pea-L-Ala-D-*N*-Me-Trp-L-Dab(Boc)-*O*]-hex-5-en (25):**

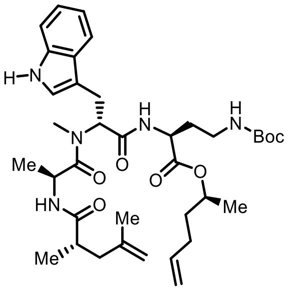

Diene **25** was synthesized by **General procedure 2 and 3** from ester **21** (100 mg) and dipeptide **18** (69 mg). By using FCC (EtOAc/PE 1:1), peptide **25** was obtained as an off-white solid (71 mg, 58% yield).

**TLC: $R_f$ =** 0.45 (PE/EtOAc = 3:7).

**$[\alpha]_D^{22}$ =** +92.0 (c = 0.1, $CHCl_3$/MeOH = 1:1).

**IR (ATR):** $\tilde{\nu}$ **=** 2978 (w), 2934 (w), 2481 (w), 2436 (w), 2405 (w), 2359 (w), 2238 (w), 2115 (w), 2074 (w), 2013 (w), 1980 (w), 1793 (w), 1637 (s), 1520 (w), 1451 (m), 1366 (m), 1164 (m), 1099 (m), 1044 (w), 998 (w), 841 (s), 741 (s) $cm^{-1}$.

**$^1$H NMR** (400 MHz, chloroform-*d*:methanol-$d_4$ = 9:1, 297 K): $\delta$ = 7.85 (d, *J* = 7.8 Hz, 1H), 7.66 (dt, *J* = 70.6, 37.6 Hz, 3H), 7.65 – 7.58 (m, 1H), 7.38 (dt, *J* = 14.8, 7.2 Hz, 2H), 6.03 (d, *J* = 6.7 Hz, 1H), 5.80 (s, 1H), 5.24 (dd, *J* = 25.1, 13.0 Hz, 3H), 5.01 (s, 1H), 4.93 (s, 1H), 4.82 (dd, *J* = 15.4, 11.1 Hz, 2H), 4.38 (d, *J* = 7.1 Hz, 1H), 4.23 (dd, *J* = 24.7, 6.0 Hz, 1H), 3.72 (s, 4H), 3.44 (d, *J* = 5.2 Hz, 2H), 3.12 (s, 1H), 2.72 (d, *J* = 7.1 Hz, 2H), 2.31 (s, 6H), 1.89 (d, *J* = 33.4 Hz, 6H), 1.70 (s, 9H), 1.52 (s, 5H), 1.33 (d, *J* = 6.9 Hz, 3H), 1.19 (s, 3H) ppm.

**$^{13}$C{$^1$H} NMR** (101 MHz, chloroform-*d*:methanol-$d_4$ = 9:1, 297 K): $\delta$ = 176.9, 176.9, 174.3, 172.3, 171.8, 171.7, 170.7, 156.4, 142.7, 137.3, 136.2, 135.8, 127.1, 122.6, 122.5, 121.7, 119.0, 118.2, 116.7, 115.2, 112.2, 111.3, 109.9, 79.5, 72.2, 69.9, 60.6, 57.2, 50.0, 45.8, 45.7, 41.5, 38.4, 37.5, 36.3, 34.8, 32.2, 31.6, 31.3, 29.5, 28.3, 23.6, 21.9, 20.9, 19.6, 18.0, 16.4, 16.2, 14.0 ppm.

**HRMS** (ESI−TOF) *m/Z*: [M + H]$^+$ calculated for $C_{37}H_{56}N_5O_7^+$ 682.4174; found: 682.4173.

**(2*S*)-2-[L-Pea-L-Ala-D-*N*-Me-Trp-L-Dap(Boc)-*O*]-hex-5-en (26):**

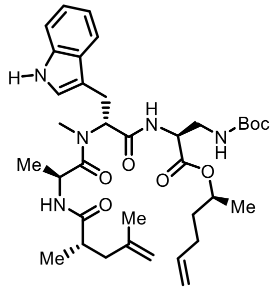

Diene **26** was synthesized by **General procedure 2 and 3** from ester **22** (130 mg) and dipeptide **18** (93 mg). By using FCC (EtOAc/PE 1:1), peptide **26** was obtained as a colorless solid (109 mg, 70% yield).

**TLC:** $R_f$ **=** 0.52 (PE/EtOAc = 3:7).

**$[\alpha]_D^{22}$ =** +292.2 (c = 0.1, $CHCl_3$/MeOH = 1:1).

**IR (ATR):** $\tilde{\nu}$ **=** 3408 (w), 3074 (w), 3056 (w), 2977 (w), 2167 (w), 2007 (w), 1982 (w), 1719 (m), 1629 (s), 1521 (m), 1439 (m), 1414 (m), 1367 (m), 1335 (w), 1298 (w), 1061 (w), 1009 (w), 843 (s), 786 (w), 740 (s), 647 (w), 634 (w), 622 (w), 602 (w) $cm^{-1}$.

**$^1$H NMR** (400 MHz, chloroform-*d*:methanol-$d_4$ = 8:2, 297 K): $\delta$ = 7.72 (dd, *J* = 89.8, 7.9 Hz, 3H), 7.36 (d, *J* = 29.3 Hz, 2H), 6.92 (s, 1H), 6.04 (dd, *J* = 22.9, 12.6 Hz, 2H), 5.42 – 5.14 (m, 3H), 5.03 (s, 1H), 4.94 (s, 1H), 4.80 (dd, *J* = 6.5, 3.1 Hz, 1H), 4.72 (d, *J* = 6.9 Hz, 1H), 4.44 – 4.15 (m, 1H), 4.02 (s, 1H), 3.80 (d, *J* = 4.7 Hz, 1H), 3.65 (s, 1H), 3.51 – 3.37 (m, 1H), 2.78 (d, *J* = 7.8 Hz, 1H), 2.66 – 2.53 (m, 1H), 2.33 (d, *J* = 36.3 Hz, 4H), 1.95 (s, 5H), 1.79 – 1.62 (m, 9H), 1.53 (dd, *J* = 11.0, 4.2 Hz, 4H), 1.35 (d, *J* = 6.9 Hz, 3H), 1.20 (dd, *J* = 25.5, 6.7 Hz, 4H) ppm.

**$^{13}$C{$^1$H} NMR** (101 MHz, chloroform-*d*:methanol-$d_4$ = 8:2, 297 K): $\delta$ = 177.9, 177.9, 174.4, 171.2, 170.5, 169.9, 156.7, 156.6, 142.6, 137.5, 136.5, 136.3, 135.9, 127.2, 122.5, 122.3, 121.5, 118.8, 118.1, 116.5, 115.0, 112.2, 111.3, 110.0, 109.9, 79.7, 72.4, 69.9, 60.6, 56.7, 54.1, 53.9, 45.9, 45.8, 41.3, 41.1, 40.9, 38.2, 38.1, 37.6, 34.7, 32.3, 31.2, 29.5, 29.4, 28.2, 28.1, 23.3, 21.9, 19.5, 17.2, 16.3, 15.2, 15.2, 13.9 ppm.

**HRMS** (ESI−TOF) *m/Z*: $[M + H]^+$ calculated for $C_{36}H_{54}N_5O_7^+$ 668.4018; found: 668.4025.

##### 7.2.3.3 Synthesis of macrocycles

***cyclo*-[(2*S*,4*E*,8*S*)Hdn-L-Ala-D-*N*-Me-Trp-L-Lys(Boc)] (7):**

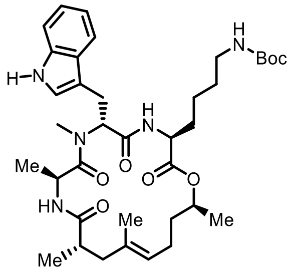

Macrocycle **7** was synthesized by **General procedure 4**, from diene **23** (116 mg). By using FCC (EtOAc/PE 7:3), cyclodepsipeptide **7** was obtained as an off-white solid (86 mg, 78% yield).

**TLC: $R_f$ =** 0.35 (PE/EtOAc = 3:7).

**$[\alpha]_D^{22}$ =** -2.7 (c = 1, $CHCl_3$/MeOH = 1:1).

**IR (ATR):** $\tilde{\nu}$ = 2394 (w), 2347 (s), 2285 (w), 2259 (w), 2224 (w), 2200 (w), 2162 (w), 1900 (w), 1695 (s), 1664 (s), 1583 (m), 1530 (m), 1499 (m), 1385 (s), 1195 (m), 1115 (m), 1022 (m), 873 (m), 791 (m), 769 (m), 700 (m), 679 (m), 626 (m) $cm^{-1}$.

**$^{1}$H NMR** (400 MHz, chloroform-*d*:methanol-$d_4$ = 9:1, 297 K): $\delta$ = 7.48 (d, *J* = 7.3 Hz, 1H), 7.28 – 7.19 (m, 2H), 6.97 (ddd, *J* = 33.6, 21.7, 5.8 Hz, 3H), 6.68 (d, *J* = 5.7 Hz, 1H), 5.27 (dd, *J* = 13.9, 6.7 Hz, 1H), 4.93 (dd, *J* = 28.6, 6.2 Hz, 1H), 4.70 (dd, *J* = 12.4, 6.3 Hz, 2H), 4.33 – 4.06 (m, 1H), 3.86 – 3.54 (m, 1H), 3.31 – 3.12 (m, 3H), 3.11 – 2.95 (m, 4H), 2.85 (d, *J* = 5.7 Hz, 2H), 2.11 (ddd, *J* = 36.4, 30.6, 23.7 Hz, 2H), 2.00 – 1.57 (m, 3H), 1.50 – 1.39 (m, 4H), 1.35 – 1.11 (m, 12H), 1.02 (dd, *J* = 25.4, 5.6 Hz, 5H), 0.89 (d, *J* = 6.0 Hz, 5H), 0.62 (d, *J* = 4.9 Hz, 1H) ppm.

**$^{13}$C{$^{1}$H} NMR** (101 MHz, chloroform-*d*:methanol-$d_4$ = 9:1, 297 K): $\delta$ = 175.4, 173.6, 170.8, 170.3, 169.8, 156.8, 136.4, 133.4, 126.8, 124.9, 124.2, 123.0, 121.6, 118.9, 118.1, 111.4, 109.2, 79.4, 71.6, 69.7, 55.9, 52.9, 49.4, 49.1, 48.9, 48.7, 48.5, 48.3, 48.1, 45.2, 45.1, 44.5, 43.7, 40.6, 40.5, 39.9, 35.2, 33.9, 31.3, 30.6, 30.4, 29.2, 28.2, 25.3, 22.8, 22.4, 18.9, 18.3, 18.0, 16.2, 15.8 ppm.

**HRMS** (ESI−TOF) *m/Z*: $[M + H]^+$ calculated for $C_{37}H_{56}N_5O_7^+$ 682.4174; found: 682.4187.

**cyclo-[(2S,4E,8S)Hdn-L-Ala-D-N-Me-Trp-L-Orn(Boc)] (6):**

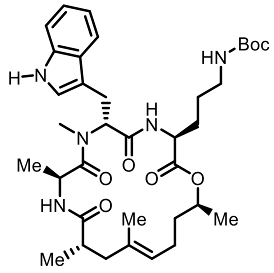

Macrocycle **6** was synthesized by **General procedure 4**, from diene **24** (75 mg). By using FCC (EtOAc/PE 7:3, then 1:0), cyclodepsipeptide **6** was obtained as an off-white solid (59 mg, 81% yield).

**TLC:** $R_f$ **=** 0.28 (PE/EtOAc = 3:7).

$[\alpha]_D^{22}$ **=** -1.5 (c = 1, $CHCl_3$/MeOH = 1:1).

**IR (ATR):** $\tilde{\nu}$ = 2957 (s), 2884 (m), 2394 (w), 2336 (w), 2227 (w), 2193 (w), 2036 (w), 2009 (w), 1961 (w), 1881 (w), 1858 (w), 1664 (s), 1586 (m), 1479 (s), 1403 (s), 1306 (m), 1183 (s), 1120 (s), 1046 (s), 971 (m), 931 (m), 885 (m), 837 (m), 818 (m), 708 (m), 630 (s) $cm^{-1}$.

**$^1$H NMR** (250 MHz, chloroform-*d,* 297 K): $\delta$ = 8.69 (s, 1H), 7.65 (d, *J* = 7.4 Hz, 1H), 7.41 (d, *J* = 7.4 Hz, 1H), 7.26 – 7.12 (m, 2H), 7.04 (s, 1H), 6.50 (dd, *J* = 17.1, 7.0 Hz, 1H), 6.18 (d, *J* = 8.7 Hz, 1H), 5.44 (d, *J* = 2.5 Hz, 1H), 5.08 (s, 1H), 4.90 (dd, *J* = 8.8, 6.6 Hz, 2H), 4.65 – 4.41 (m, 2H), 4.15 (d, *J* = 7.1 Hz, 1H), 3.46 (dd, *J* = 14.3, 9.4 Hz, 1H), 3.13 (d, *J* = 3.9 Hz, 3H), 3.05 (dd, *J* = 11.5, 6.4 Hz, 2H), 2.83 (s, 2H), 2.30 (t, *J* = 8.5 Hz, 2H), 2.17 – 1.64 (m, 6H), 1.62 – 1.43 (m, 14H), 1.17 (dd, *J* = 6.6, 4.1 Hz, 12H) ppm.

**$^{13}$C{$^1$H} NMR** (63 MHz, chloroform-*d,* 297 K): $\delta$ = 174.6, 173.8, 169.8, 156.2, 136.3, 133.9, 127.1, 124.9, 122.1, 122.0, 119.6, 118.5, 111.4, 110.3, 71.8, 56.3, 52.7, 45.5, 43.8, 41.1, 40.0, 38.6, 35.6, 30.7, 29.5, 28.5, 26.2, 24.6, 24.5, 23.2, 21.1, 19.6, 18.8, 18.7, 16.8, 14.2, 16.8, 14.2 ppm.

**HRMS** (ESI−TOF) *m/Z*: [M + H]$^+$ calculated for $C_{36}H_{54}N_5O_7^+$ 668.4018; found: 668.4019.

***cyclo*-[(2*S*,4*E*,8*S*)Hdn-L-Ala-D-*N*-Me-Trp-L-Dab(Boc)] (5):**

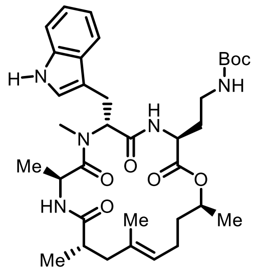

Macrocycle **5** was synthesized by **General procedure 4**, from diene **25** (55 mg). By using FCC (EtOAc/PE 7:3), the cyclodepsipeptide **5** was obtained as a colorless solid (37 mg, 71% yield).

**TLC: *R<sub>f</sub>* =** 0.55 (EtOAc = 100%).

**$[\alpha]_D^{22}$ =** 1.8 (c = 0.1, $CHCl_3$/MeOH = 1:1).

**IR (ATR):** $\tilde{\nu}$ = 2952 (m), 2335 (w), 2232 (w), 2190 (w), 2149 (w), 2077 (w), 2043 (w), 1673 (s), 1583 (m), 1471 (s), 1424 (m), 1401 (m), 1380 (m), 1322 (m), 1197 (s), 1119 (s), 1025 (m), 885 (w), 840 (m), 796 (m), 713 (m), 647 (s), 626 (s) $cm^{-1}$.

**$^1$H NMR** (400 MHz, chloroform-*d*:methanol-$d_4$ = 9:1, 297 K): $\delta$ = 7.86 (d, *J* = 7.6 Hz, 1H), 7.66 – 7.55 (m, 1H), 7.38 (d, *J* = 22.9 Hz, 3H), 6.94 (d, *J* = 7.2 Hz, 1H), 5.71 (s, 1H), 5.28 (s, 1H), 5.17 – 4.93 (m, 2H), 4.79 – 4.56 (m, 1H), 4.22 – 3.98 (m, 1H), 3.75 – 3.51 (m, 6H), 3.40 (d, *J* = 8.3 Hz, 1H), 3.27 (dd, *J* = 14.8, 6.3 Hz, 1H), 2.96 – 2.74 (m, 1H), 2.67 – 2.55 (m, 1H), 2.45 (s, 1H), 2.29 (s, 1H), 2.21 (d, *J* = 13.0 Hz, 2H), 2.17 – 2.03 (m, 3H), 1.81 (d, *J* = 4.8 Hz, 5H), 1.70 (d, *J* = 5.1 Hz, 12H), 1.44 (d, *J* = 6.2 Hz, 3H), 1.41 – 1.31 (m, 3H), 1.20 (d, *J* = 6.7 Hz, 3H), 1.04 (t, *J* = 26.3 Hz, 1H) ppm.

**$^{13}$C{$^1$H} NMR** (101 MHz, chloroform-*d*:methanol-$d_4$ = 9:1, 297 K): $\delta$ = 175.3, 175.2, 173.8, 171.6, 171.2, 170.6, 170.4, 156.4, 136.5, 136.3, 134.1, 133.5, 126.9, 125.0, 124.2, 123.1, 122.9, 121.9, 119.1, 118.3, 111.4, 109.6, 79.6, 73.3, 71.9, 69.9, 60.6, 55.9, 50.9, 50.9, 45.3, 45.2, 44.5, 43.8, 40.8, 40.7, 36.3, 35.4, 33.9, 31.9, 30.7, 30.6, 30.5, 29.6, 28.3, 25.0, 24.8, 23.0, 20.9, 20.3, 19.1, 18.4, 18.3, 16.4, 15.9, 15.9, 14.1 ppm.

**HRMS** (ESI−TOF) *m/Z*: $[M + H]^+$ calculated for $C_{35}H_{52}N_5O_7^+$ 654.3861; found: 654.3860.

***cyclo*-[(2*S*,4*E*,8*S*)Hdn-L-Ala-D-*N*-Me-Trp-L-Dap(Boc)] (4):**

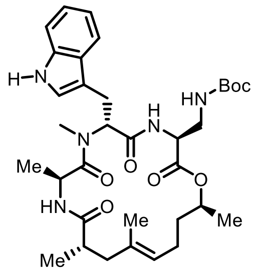

Macrocycle **4** was synthesized by **General procedure 4**, from diene **26** (95 mg). By using automated FCC (gradient EtOAc/PE 7:3 to 1:0), the cyclopeptide **4** was obtained as a colorless solid (67 mg, 75% yield).

**TLC:** $\boldsymbol{R_f}$ **=** 0.22 (PE/EtOAc = 3:7).

$\mathbf{[\alpha]_D^{22}}$ **=** 5.1 (c = 1, $CHCl_3$/MeOH = 1:1).

**IR (ATR):** $\tilde{\nu}$ = 2335 (m), 2235 (w), 2195 (w), 2173 (m), 2150 (w), 2110 (m), 2030 (w), 2001 (w), 1953 (w), 1931 (w), 1880 (m), 1857 (m), 1752 (w), 1663 (s), 1585 (m), 1467 (s), 1426 (m), 1377 (m), 1218 (m), 1116 (s), 1019 (s), 930 (s), 884 (s), 706 (s), 646 (s), 623 (s) $cm^{-1}$.

$^{1}$**H NMR** (500 MHz, chloroform-*d*:methanol-$d_4$ = 9:1, 297 K): $\delta$ = 7.50 (d, *J* = 7.7 Hz, 1H), 7.25 (dd, *J* = 12.8, 4.4 Hz, 1H), 7.04 (t, *J* = 7.5 Hz, 1H), 6.98 (t, *J* = 7.4 Hz, 1H), 6.88 (s, 1H), 6.62 (d, *J* = 7.0 Hz, 1H), 5.43 (s, 1H), 4.90 (s, 1H), 4.78 (d, *J* = 6.0 Hz, 1H), 4.65 (s, 1H), 4.33 (s, 1H), 3.74 (s, 3H), 3.39 – 3.29 (m, 1H), 3.29 – 3.19 (m, 1H), 3.14 (t, *J* = 11.4 Hz, 2H), 2.93 (s, 3H), 2.23 (s, 1H), 1.93 (s, 1H), 1.85 (t, *J* = 17.5 Hz, 1H), 1.81 – 1.69 (m, 2H), 1.44 (s, 3H), 1.41 – 1.25 (m, 13H), 1.11 (d, *J* = 6.1 Hz, 3H), 0.98 (d, *J* = 6.4 Hz, 4H), 0.78 (ddd, *J* = 20.9, 19.0, 7.3 Hz, 2H), 0.68 (d, *J* = 6.6 Hz, 3H) ppm.

$^{13}$**C{**$^{1}$**H} NMR** (125 MHz, chloroform-*d*:methanol-$d_4$ = 9:1, 297 K): $\delta$ = 175.4, 175.3, 175.2, 173.8, 173.6, 171.7, 171.0, 170.9, 169.1, 156.8, 142.5, 136.2, 133.2, 126.8, 125.0, 122.5, 121.7, 119.0, 118.2, 111.3, 109.5, 79.9, 72.1, 69.9, 60.5, 55.7, 54.3, 45.1, 45.0, 43.7, 41.3, 40.6, 40.5, 35.2, 30.5, 28.1, 24.9, 22.9, 20.8, 19.1, 18.2, 18.1, 18.0, 16.3, 13.9 ppm.

**HRMS** (ESI−TOF) *m/Z*: $[M + H]^+$ calculated for $C_{34}H_{50}N_5O_7^+$ 640.3705; found: 640.3700.

##### 7.2.3.4 Synthesis of fluorophores:

**MAP620(*p*-CN)-6-COOH (34):**

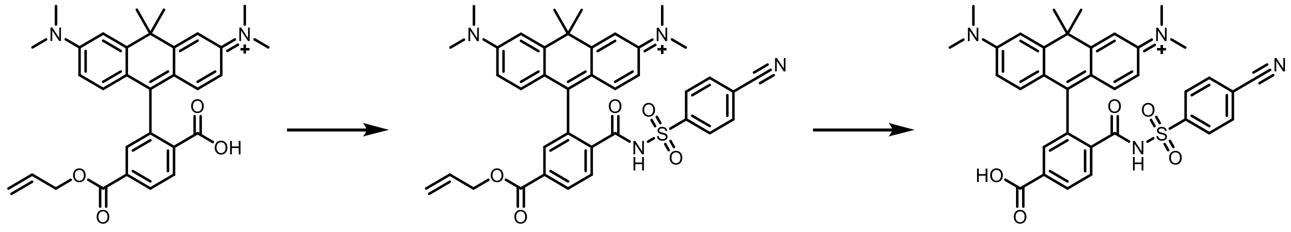

**32** **33** **34: MAP620-6-COOH**

Allyl ester **32** (12 mg, 0.024 mmol, 1 equiv.) was dissolved in $CH_2Cl_2$ (~3 mL) and $POCl_3$ (55 mg, 33.8 µL, 362 µmol, 15 equiv.) was added in one portion at 25 °C. Reaction mixture was refluxed for 2 h. Volatiles were removed in flow of nitrogen gas and slurry rest was cooled to 0 °C, diluted with anhydrous MeCN (~3 mL) followed by addition of 4-cyanobenzene-1-sulfonamide (44 mg, 0. 242 mmol, 10 eq.) and DIPEA (93 mg, 120 µL, 725 µmol, 30 equiv.). Reaction mixture was further stirred at 70 °C for 2 h and quenched by addition of saturated solution of $NaHCO_3$ (~2 mL). Mixtures was diluted with $CH_2Cl_2$ (~50 mL), washed 2x with solution of NaHCO3 (~10 mL) and anhydrated with solid $Na_2SO_4$. Product was used in the next step without further purification.

Crude product (**33**) was dissolved in mixture of $CH_2Cl_2$ and MeOH (1:4, 5 mL) at 25 °C, 1,3-dimethyl-1,3-diazinane-2,4,6-trione (22.6 mg, 0.145 mmol, 6 equiv.) and tetrakis(tri-phenylphosphine)palladium(0) (8.38 mg, 0.007 mmol, 0.3 equiv.) were added. Reaction mix-ture was stirred for 16 h and complete conversion was observed by TLC and LC/MS. Blue solid product (**34**, 4 mg 27% yield) was isolated by preparative HPLC.

**$^1$H NMR** (400 MHz, DMSO-$d_6$, 297 K) $\delta$ = 8.0 (dd, $J$ = 21.7, 7.1 Hz, 4H), 7.5 (d, $J$ = 8.1 Hz, 2H), 7.1 (d, $J$ = 73.2 Hz, 3H), 6.4 (dd, $J$ = 57.3, 8.8 Hz, 4H), 2.0 (s, 3H), 1.9 (s, 3H), 1.5 – 1.0 (m, 6H) ppm.

**HRMS** (ESI−TOF) *m/Z*: $[M + H]^+$ calculated for $C_{35}H_{33}N_4O_5S^+$ 621.2166; found: 621.2159.

##### 7.2.3.5 Synthesis of fluorophore conjugates

**SiR-XActin (8):**

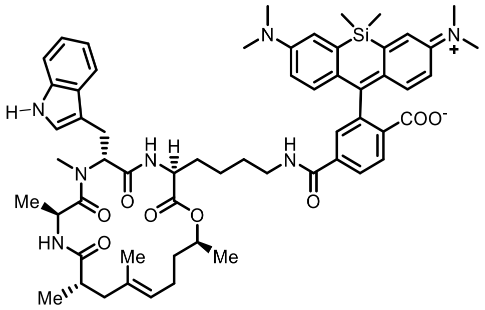

**Active ester of SiR-6-COOH:** SiR-6-COOH (16 mg, 0.035 mmol, 1.1 equiv.) was dissolved in DMF (~3 mL) and DIPEA (16 mg, 0.13 mmol, 4 equiv.) was added followed by addition of PyAOP (20 mg, 0.0.038 mmol, 1.2 equiv.). Reaction mixture was stirred for 10 min.

**Carbamate deprotection**: Hybrid carbamate **7** (22 mg, 0.032 mmol, 1 equiv.) was dissolved in DCM (~5 mL) and cooled to 0 °C. TFA (2 mL) was added and reaction mixture stirred for 15 min at the constant temperature. LC/MS and TLC monitoring showed full conversion and minimal hydrolysis. **Coupling:** After removing volatiles (at 25 °C), active ester of SiR-6-COOH was added at 25 °C to the crude ammonium residue in DMF (~ 3 mL) and reaction was stirred for 2 h. Product was isolated by preparative HPLC to provide 23 mg (69% yield) of fluorophore conjugate **8** as a green to blue solid.

**TLC: *$R_f$* =** 0.48 ($CH_2Cl_2$/MeOH = 95:5).

**$[\alpha]_D^{25}$ =** -6.1 (c = 1, $CHCl_3$/MeOH = 1:1).

**IR (ATR):** $\tilde{\nu}$ = 2224 (w), 2171 (w), 2023 (w), 1932 (w), 1787 (w), 1723 (w), 1612 (s), 1570 (m), 1466 (s), 1326 (s), 1205 (m), 1150 (s), 1113 (s), 1074 (s), 983 (s), 890 (m), 790 (s), 762 (s), 713 (s), 660 (s), 625 (s) $cm^{-1}$.

**$^1$H NMR** (500 MHz, DMSO-$d_6$, 297 K): $\delta$ = 10.80 (d, *J* = 2.5 Hz, 1H), 8.78 – 8.56 (m, 2H), 8.11 – 7.92 (m, 2H), 7.72 – 7.60 (m, 3H), 7.55 (d, *J* = 8.6 Hz, 1H), 7.29 (d, *J* = 8.0 Hz, 1H), 7.09 – 6.98 (m, 5H), 6.93 (s, 1H), 6.67 – 6.55 (m, 5H), 5.67 – 5.50 (m, 1H), 4.85 (d, *J* = 6.2 Hz, 1H), 4.82 – 4.54 (m, 1H), 4.20 (d, *J* = 6.0 Hz, 1H), 3.99 (d, *J* = 6.4 Hz, 1H), 3.56 (s, 1H), 3.20 (d, *J* = 5.8 Hz, 4H), 3.02 (s, 5H), 2.04 (s, 2H), 1.66 (d, *J* = 5.4 Hz, 5H), 1.51 (d, *J* = 5.1 Hz, 8H), 1.24 (s, 5H), 1.03 (s, 1H), 0.91 (d, *J* = 6.7 Hz, 4H), 0.64 (s, 6H), 0.52 (s, 6H) ppm.

**$^{13}$C{$^1$H} NMR** (125 MHz, DMSO-$d_6$, 297 K): $\delta$ = 174.4, 174.2, 173.7, 173.3, 172.7, 172.3, 172.0, 171.9, 170.1, 165.4, 155.6, 149.9, 140.8, 136.9, 136.5, 134.1, 133.1, 131.2, 131.2,

128.8, 128.4, 127.9, 127.7, 126.1, 124.7, 124.6, 123.9, 123.7, 123.4, 121.6, 119.5, 118.8, 117.1, 114.5, 111.8, 110.5, 110.3, 91.9, 70.8, 69.5, 55.2, 55.0, 54.4, 52.9, 45.3, 44.2, 43.8, 38.9, 38.6, 35.7, 34.1, 31.5, 31.3, 31.2, 30.7, 30.6, 29.2, 26.6, 23.7, 23.6, 23.4, 19.8, 19.4, 19.2, 18.5, 18.4, 17.3, 16.6, 16.1, 0.8, -0.5 ppm.

**HRMS** (ESI−TOF) *m/Z*: $[M]^+$ calculated for $C_{59}H_{74}N_7O_8Si^+$ 1036.5363; Found: 1036.5368.

**SiR700-XActin (9):**

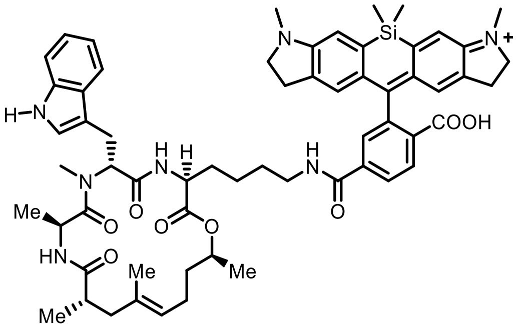

Conjugate (**910**) was synthesized following procedure described for SiR-XActin starting with XActin ligand (**7**) and fluorophore (**3029**). Preparative HPLC (collection at 680 nm) gave red solid product in yield of 45% (3.5 mg).

**HRMS** (ESI−TOF) *m/Z*: $[M]^+$ calculated for $C_{61}H_{74}N_7O_8Si^+$ 1060.5363, found 1060.5374.

**MAP555-XActin (10):**

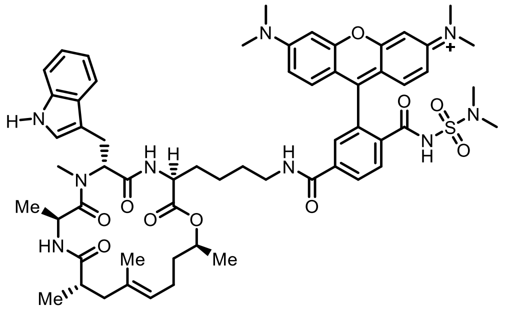

Conjugate (**10**) was synthesized following procedure described for SiR-XActin starting with XActin ligand (**7**) and fluorophore (**28**). Preparative HPLC (collection at 550 nm) gave red solid product in yield of 49% (4 mg).

**HRMS** (ESI−TOF) *m/Z*: $[M]^+$ calculated for C59H74N9O10S+ 1100.5274, found 1100.5285.

**MAP500-XActin (12):**

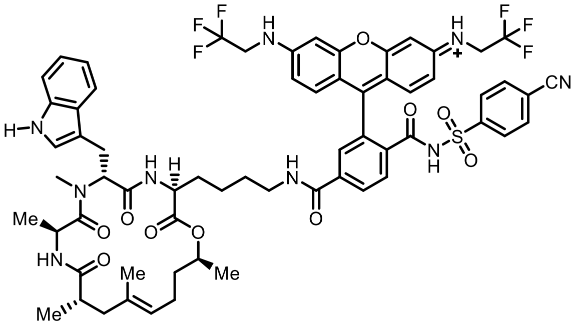

Conjugate (**12**) was synthesized following procedure described for SiR-XActin starting with XActin ligand (**7**) and fluorophore (**31**). Preparative HPLC (collection at 240 nm) gave pale pink product in yield of 32% (3 mg).

**HRMS** (ESI−TOF) *m/Z*: $[M+1]^{2+}$ calculated for $C_{64}H_{67}N_9O_{10}F_6S^{2+}$ 633.7312, found 633.7310.

**MAP620-XActin (11):**

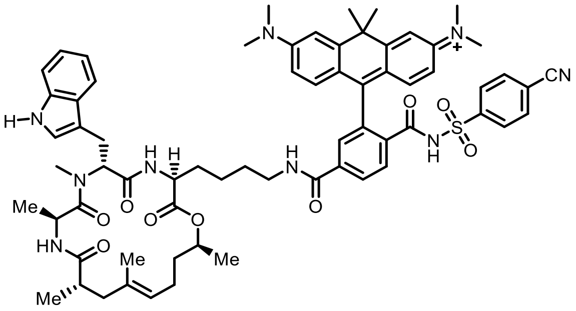

Conjugate (**11**) was synthesized following procedure described for SiR-XActin starting with XActin ligand (**7**) and fluorophore (**34**). Preparative HPLC (collection at 240 nm) gave blue solid product in yield of 34% (3 mg).

**HRMS** (ESI−TOF) *m/Z*: $[M+1]^{2+}$ calculated for $C_{67}H_{79}N_9O_9S^{2+}$ 592.7855, found 592.7856.

#### 9 NMR Spectra

#### [1]H spectrum of compound 22: 300 MHz, chloroform-*d*, 298 K

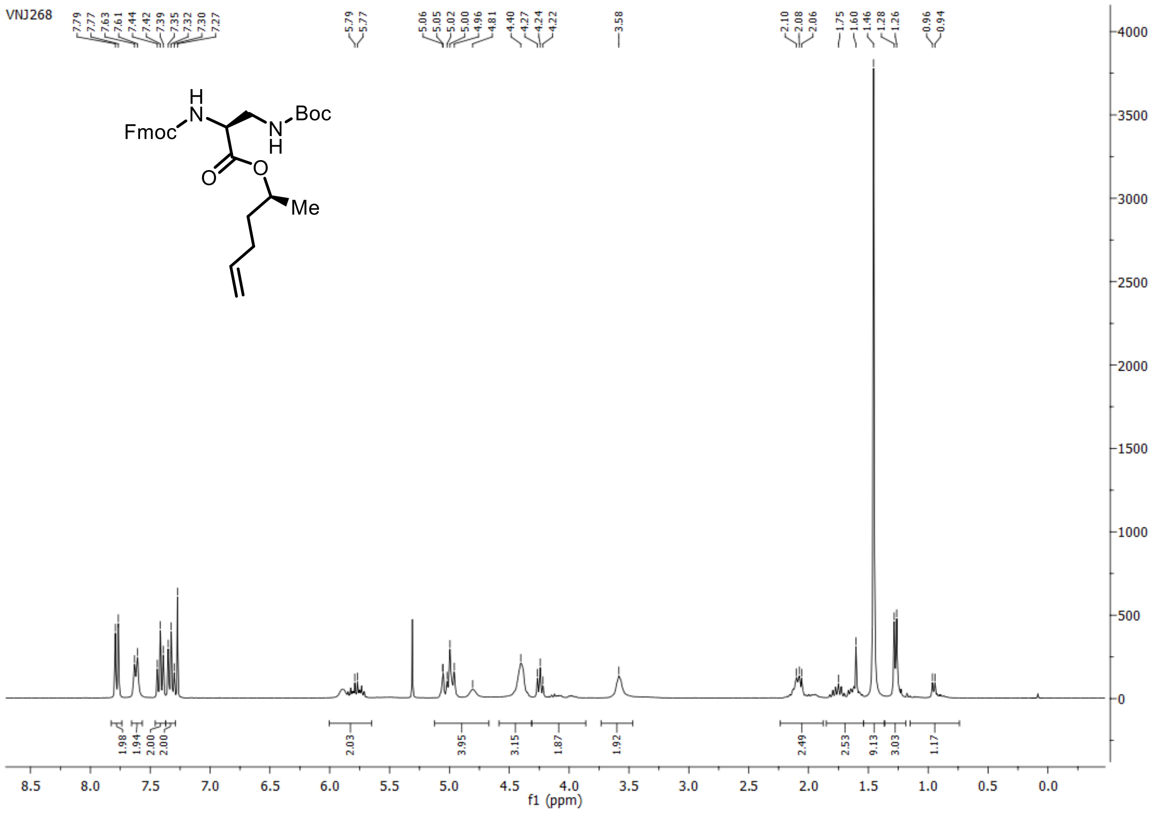

#### [13]C{[1]H} spectrum of compound 22: 75 MHz, chloroform-*d*, 298 K

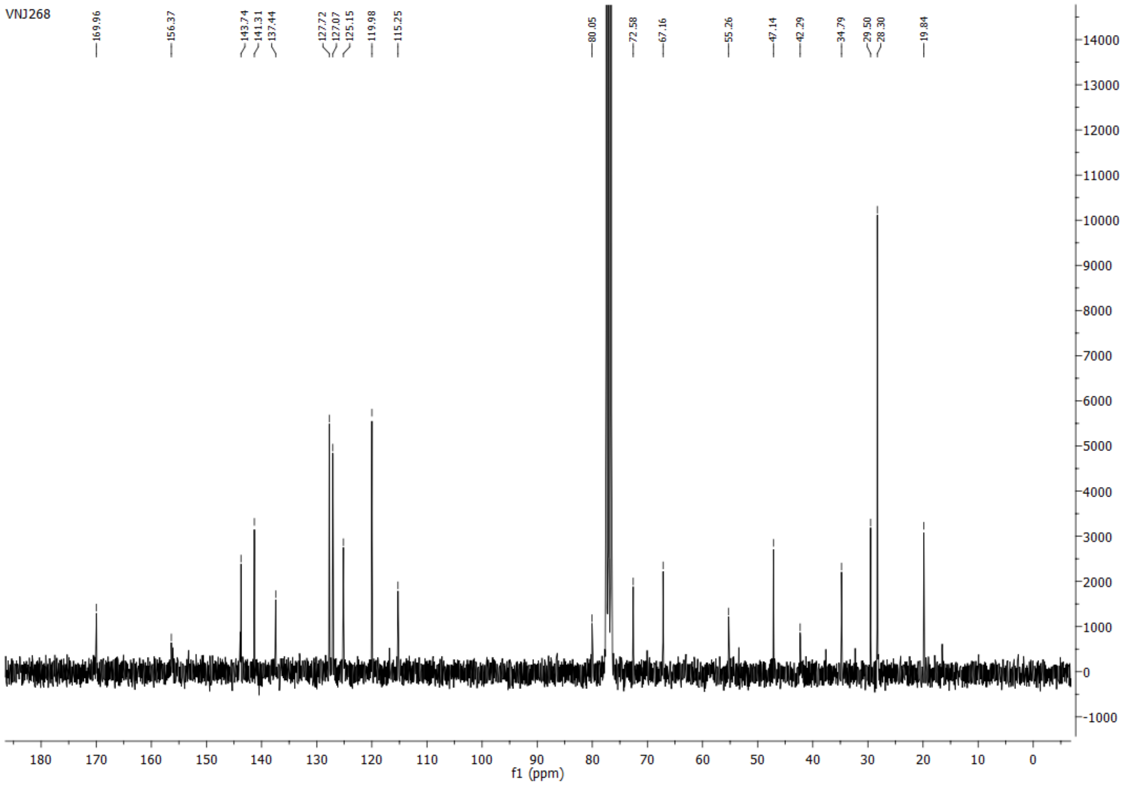

**[1]H spectrum of compound 21: 300 MHz, chloroform-*d*, 298 K**

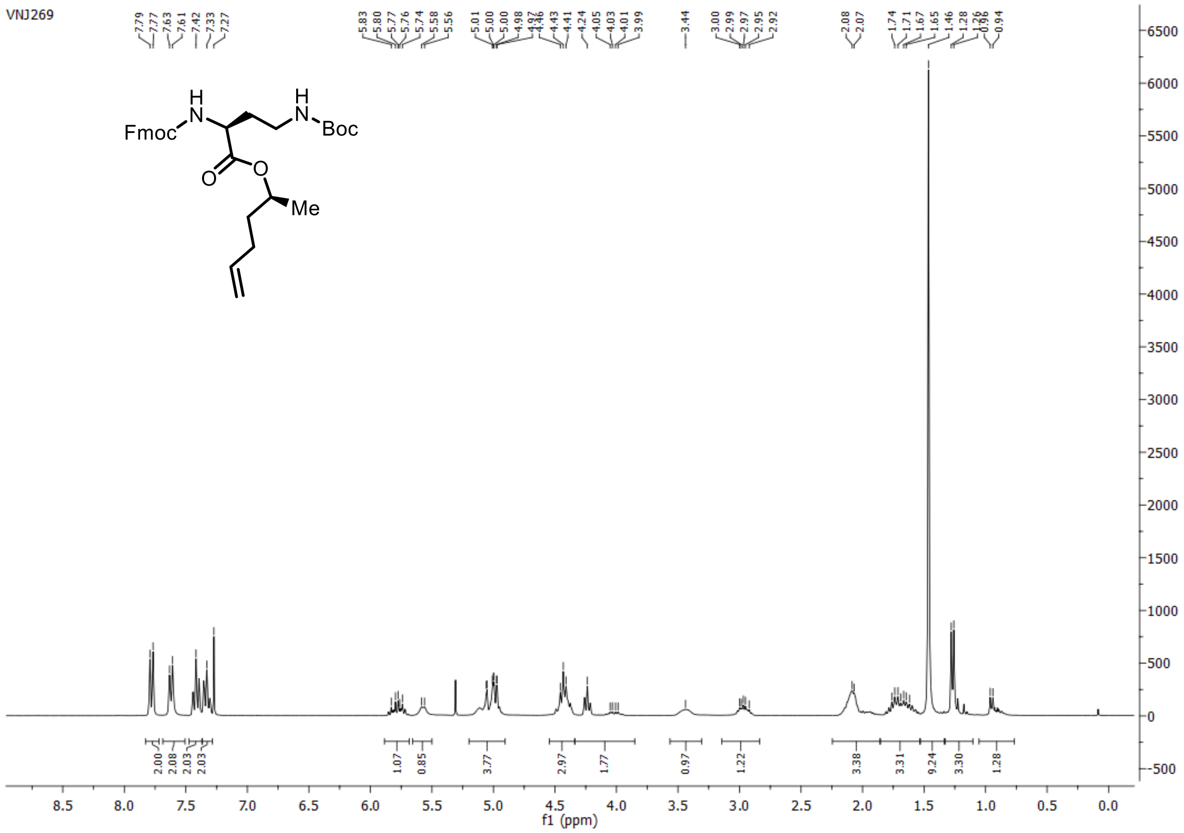

**[13]C{[1]H} spectrum of compound 21: 75 MHz, chloroform-*d*, 298 K**

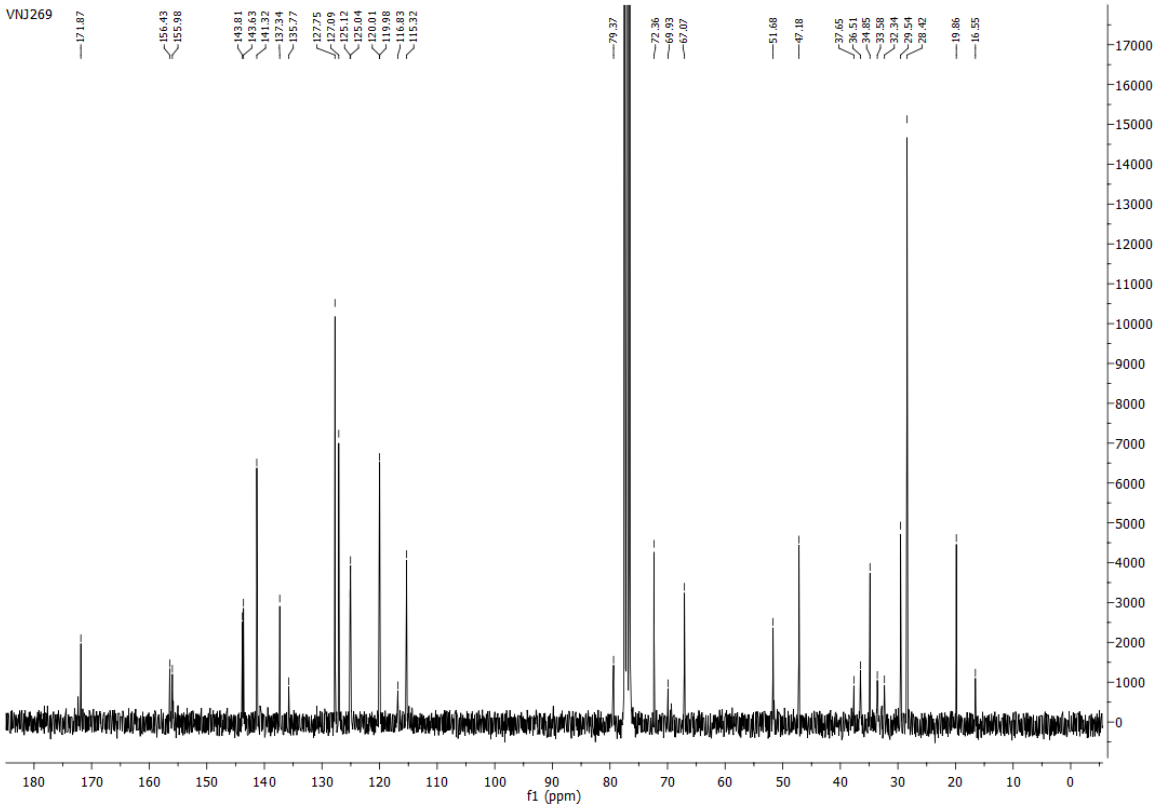

#### $^{1}H$ spectrum of compound 20: 400 MHz, chloroform-*d*, 297 K

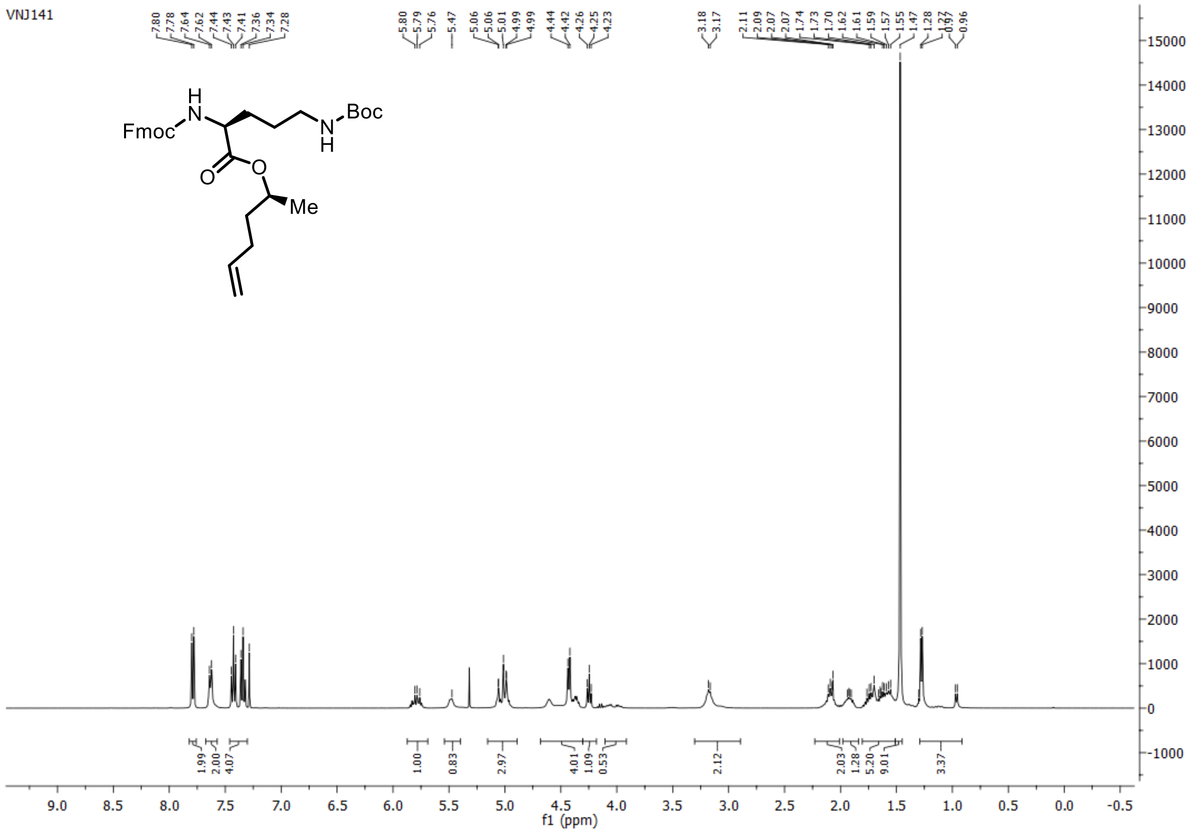

#### $^{13}C\{^{1}H\}$ spectrum of compound 20: 101 MHz, chloroform-*d*, 297 K

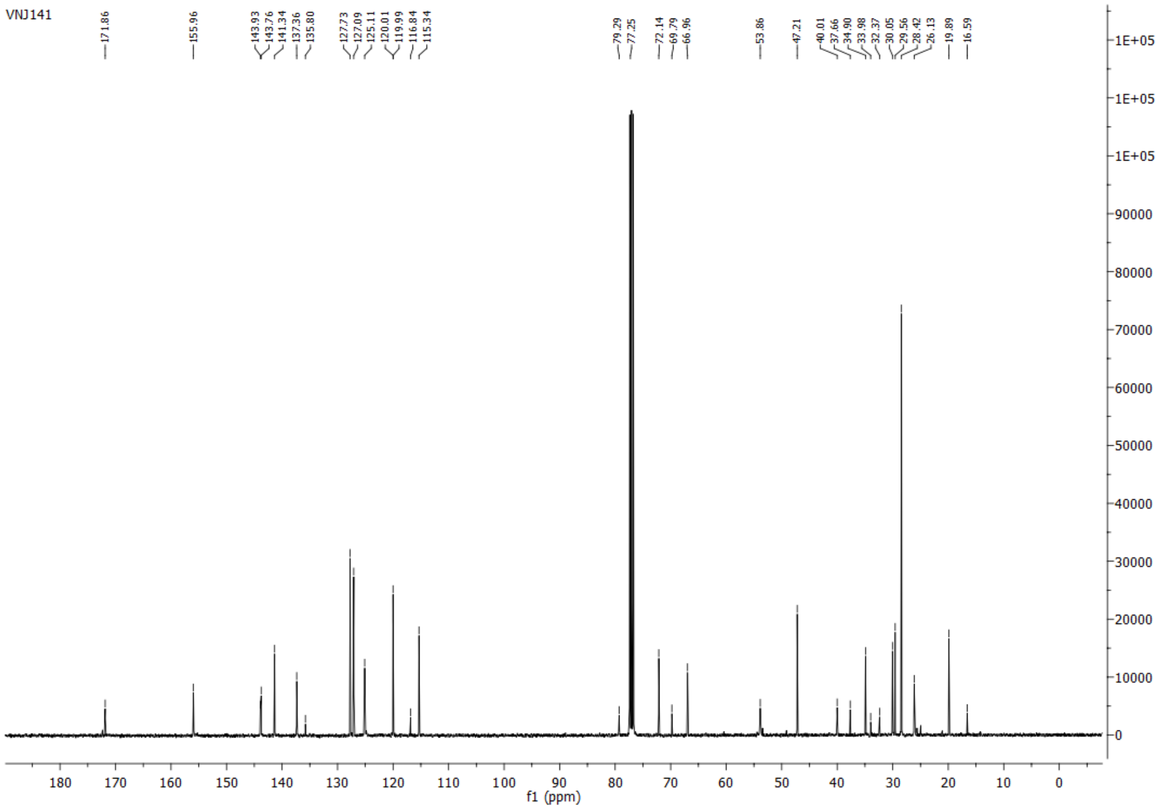

**$^{1}$H spectrum of compound 19: 400 MHz, chloroform-*d*, 297 K**

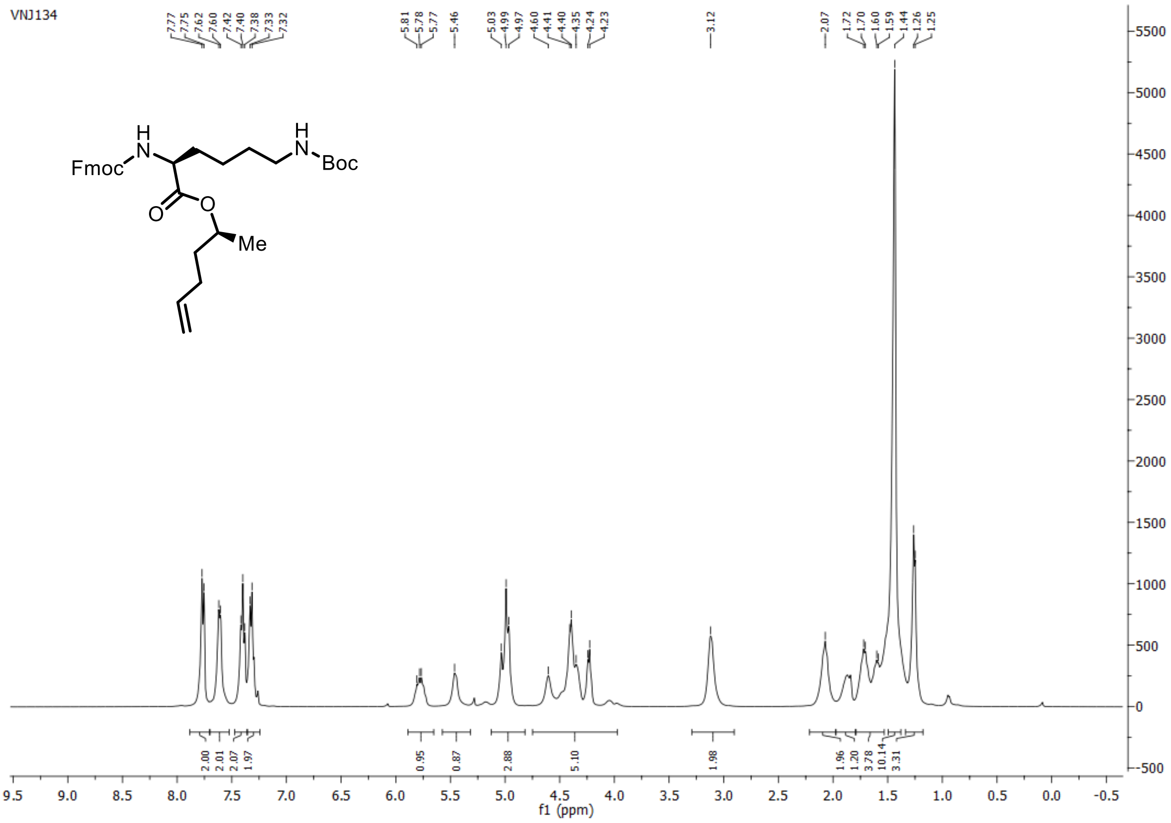

**$^{13}$C{$^{1}$H} spectrum of compound 19: 101 MHz, chloroform-*d*, 297 K**

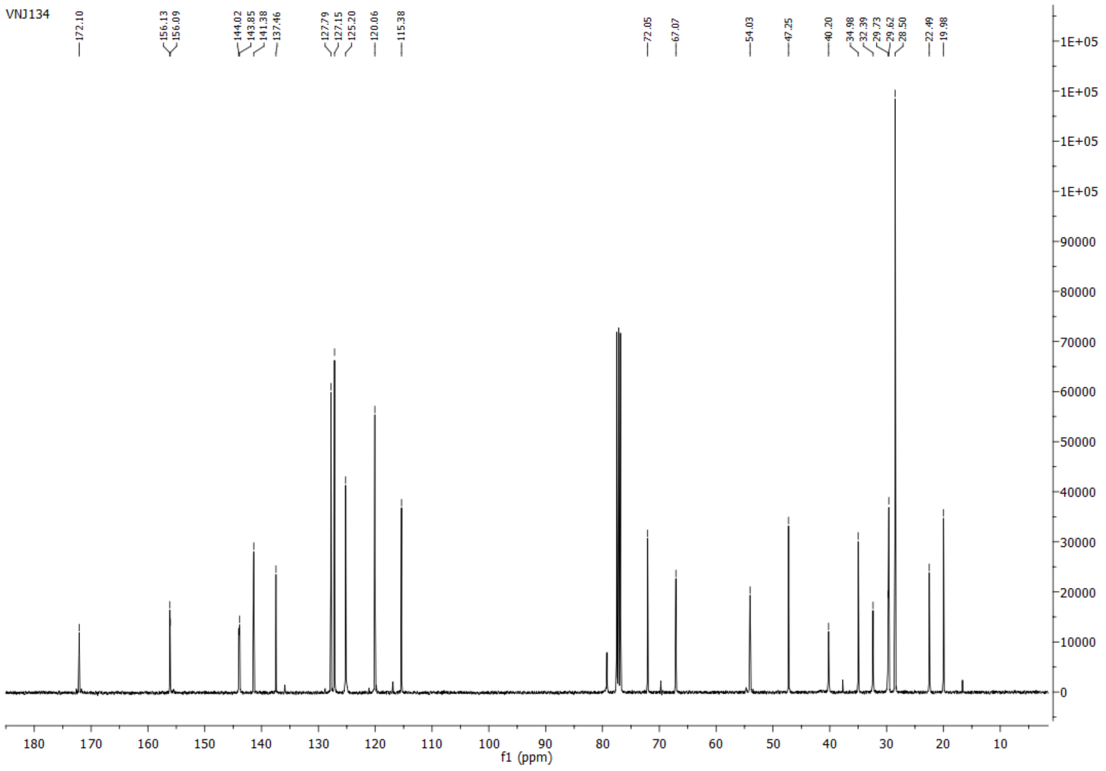

**[1]H spectrum of compound 18: 300 MHz, chloroform-*d*, 298 K**

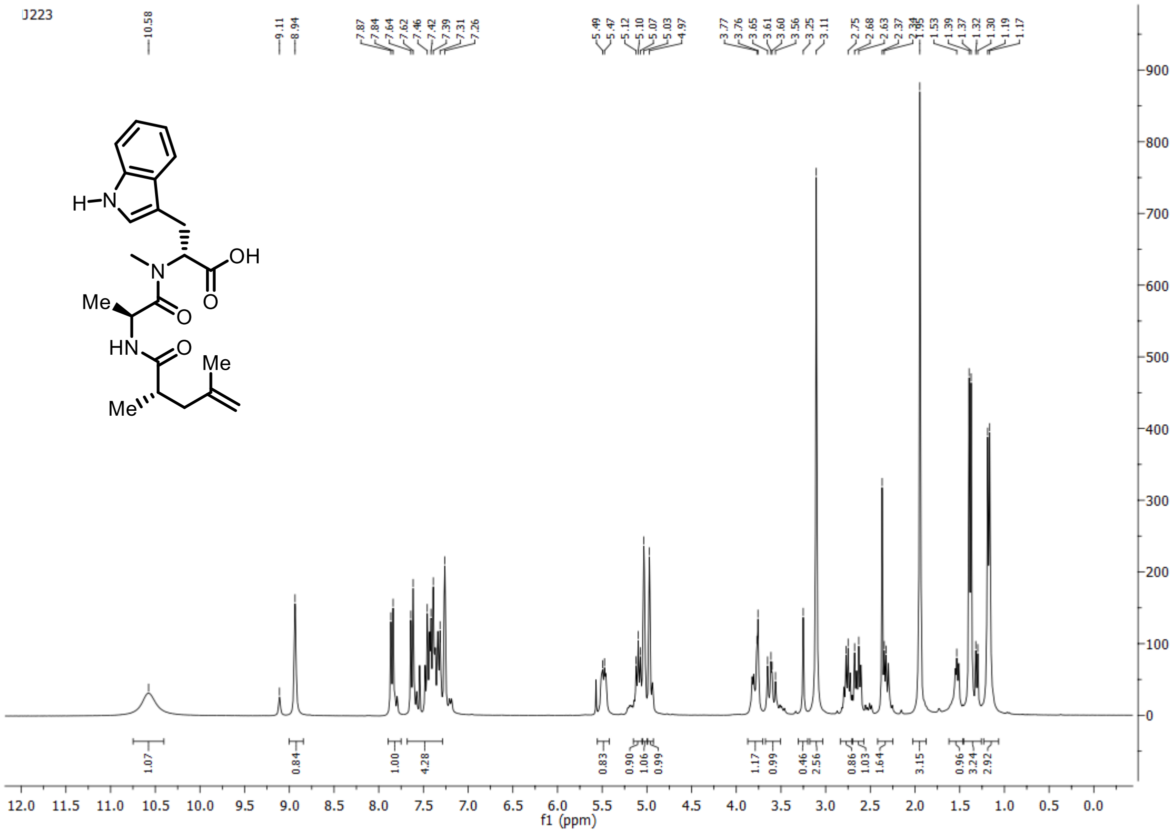

**[13]C{[1]H} spectrum of compound 18: 101 MHz, chloroform-*d*, 298 K**

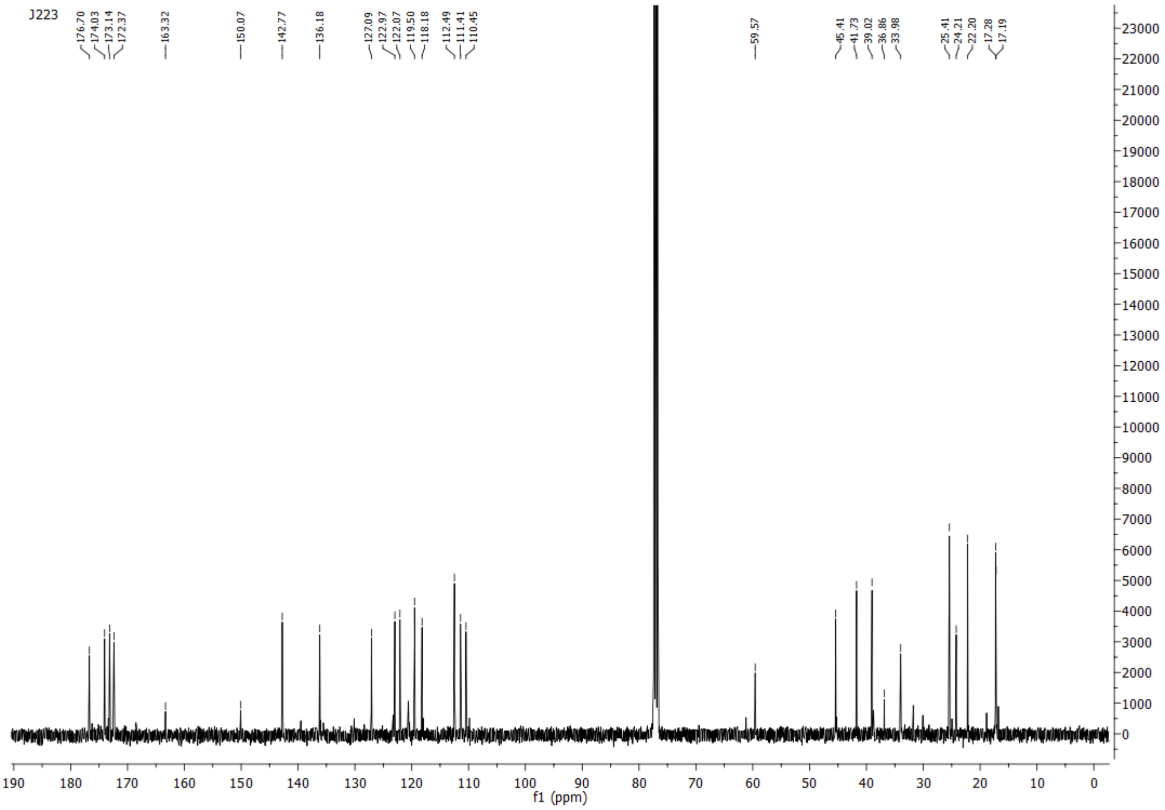

**$^1$H spectrum of compound 23: 400 MHz, chloroform-*d*:methanol-*d*$_4$ = 9:1, 297 K**

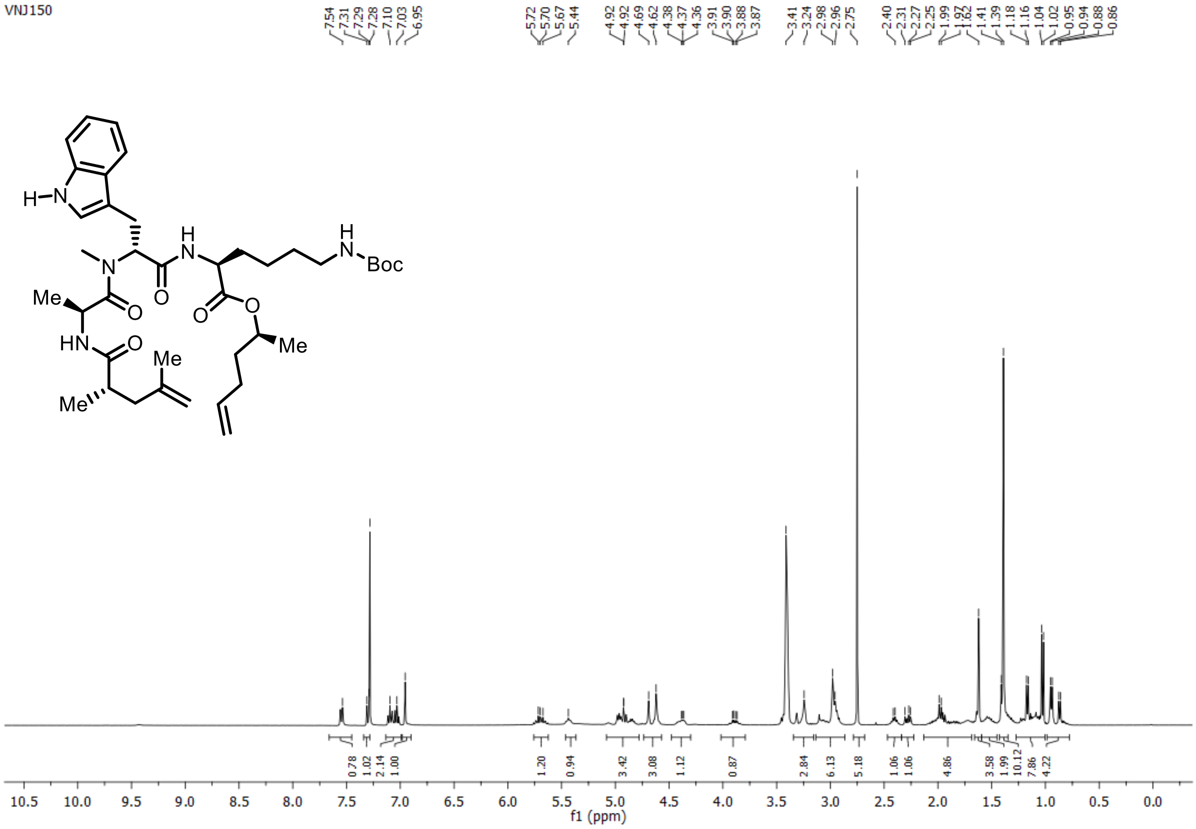

**$^{13}$C{$^1$H} spectrum of 23: 101 MHz, chloroform-*d*:methanol-*d*$_4$ = 9:1, 297 K**

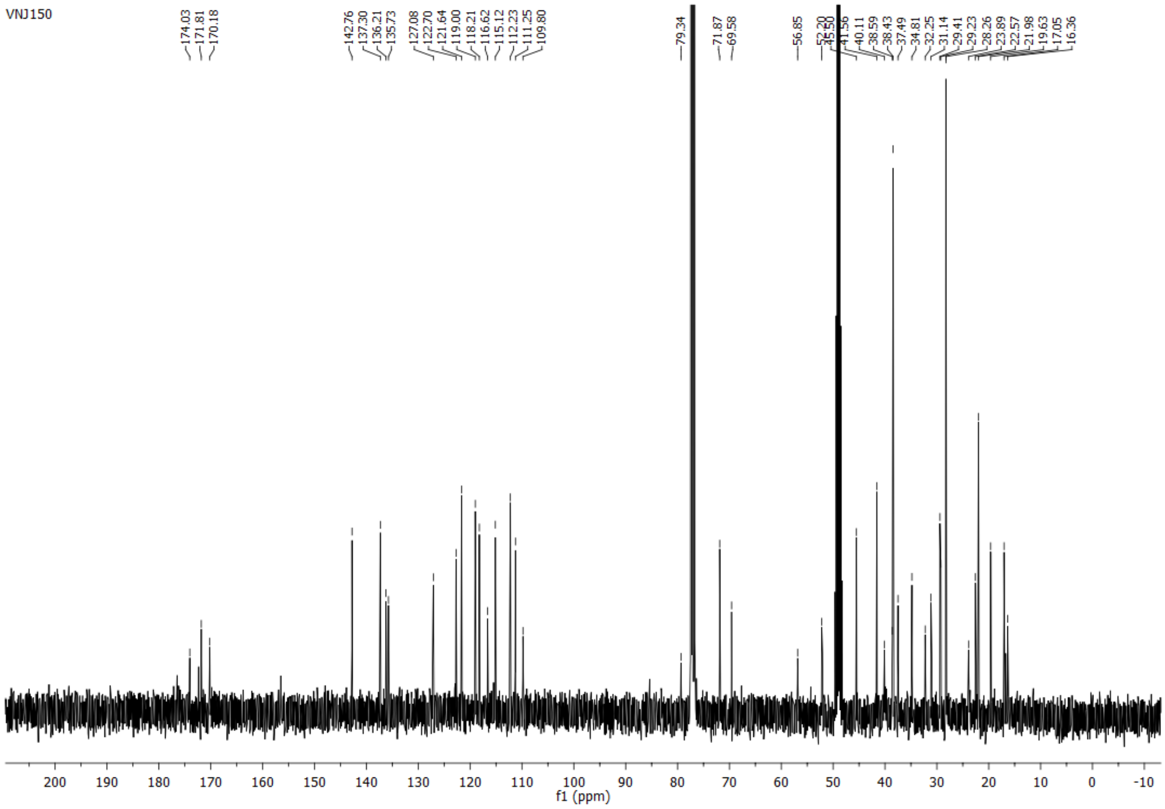

**$^{1}$H spectrum of compound 24: 300 MHz, chloroform-*d*:methanol-*d*$_4$ = 9:1, 298 K**

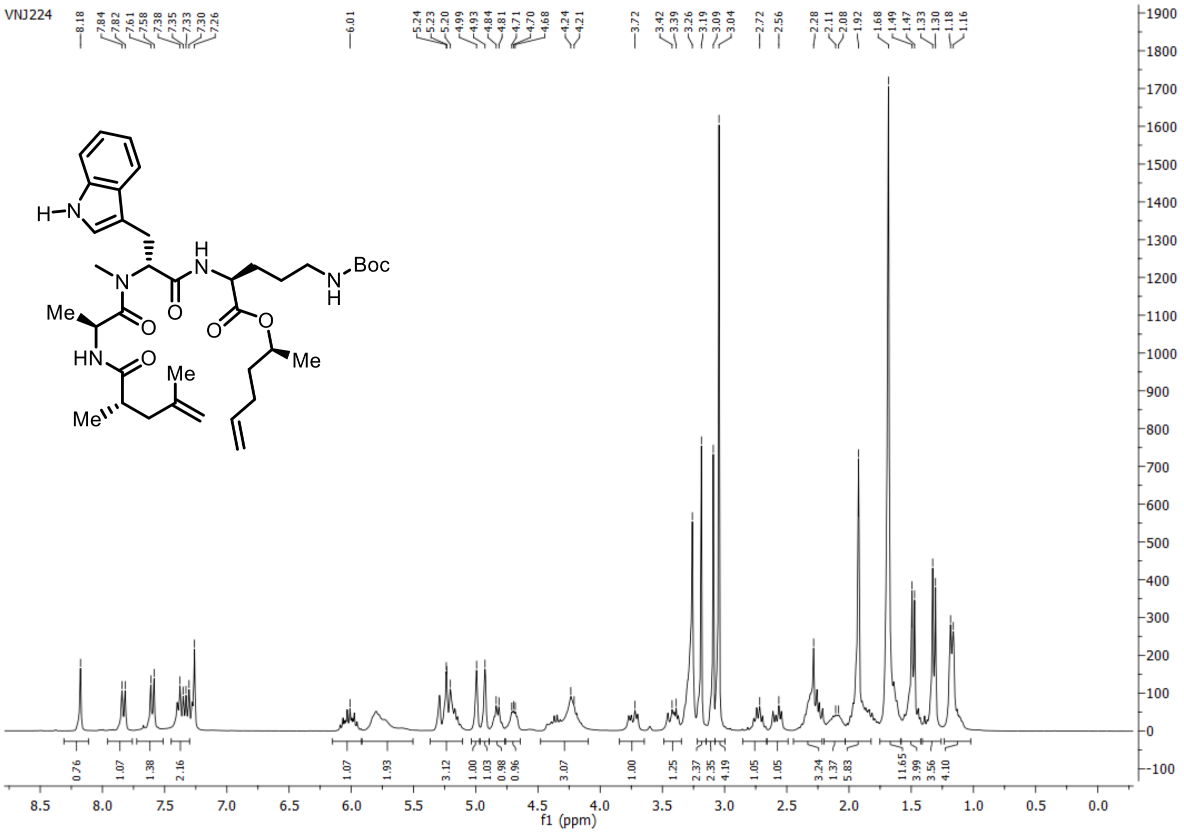

**$^{13}$C{$^{1}$H} spectrum of compound 24: 75 MHz, chloroform-*d*:methanol-*d*$_4$ = 9:1, 298 K**

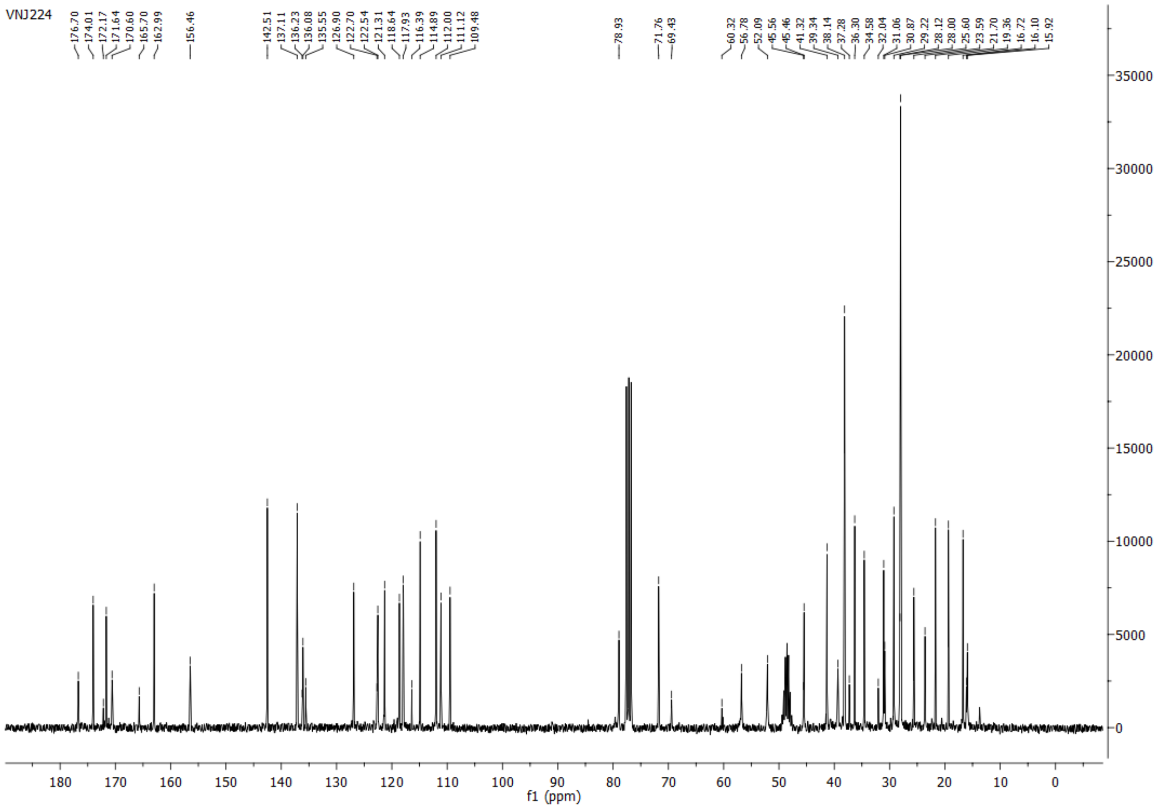

**$^{1}$H spectrum of compound 25: 400 MHz, chloroform-*d*:methanol-*d*$_4$ = 9:1, 297 K**

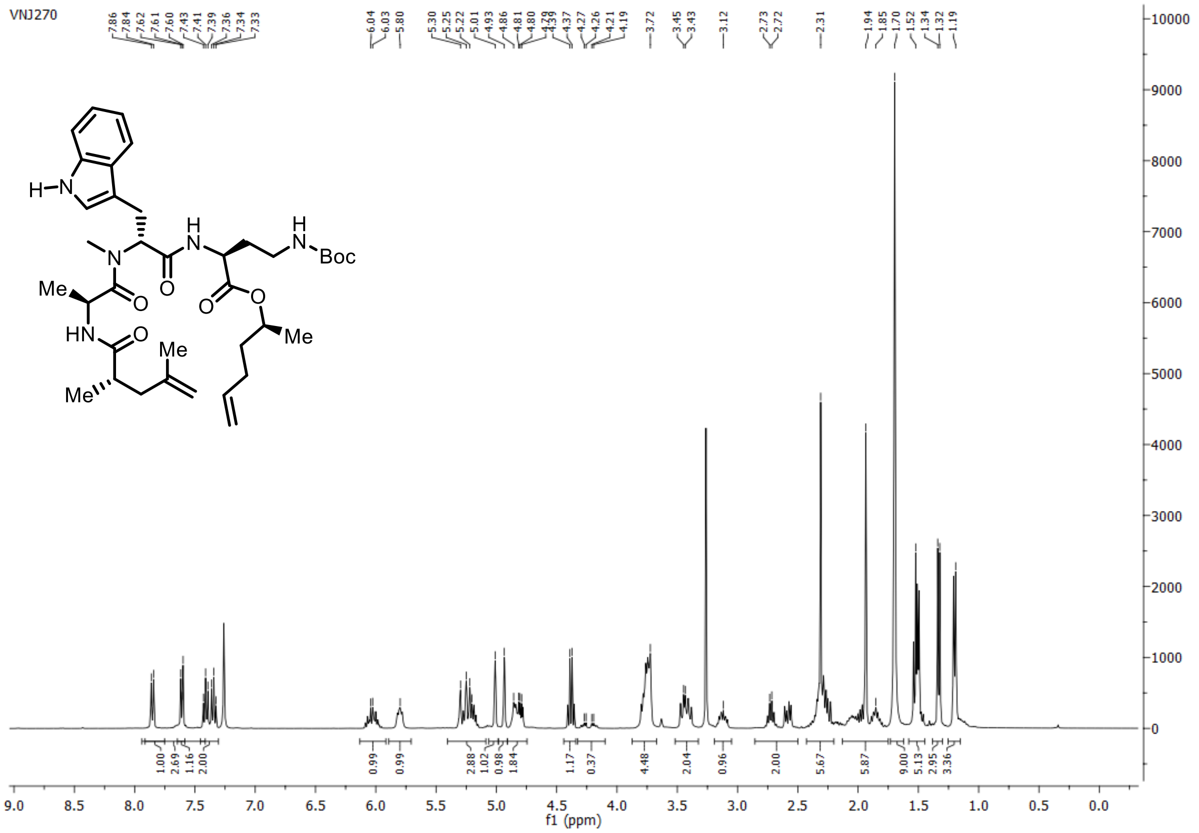

**$^{13}$C{$^{1}$H} spectrum of compound 25: 101 MHz, chloroform-*d*:methanol-*d*$_4$ = 9:1, 297 K**

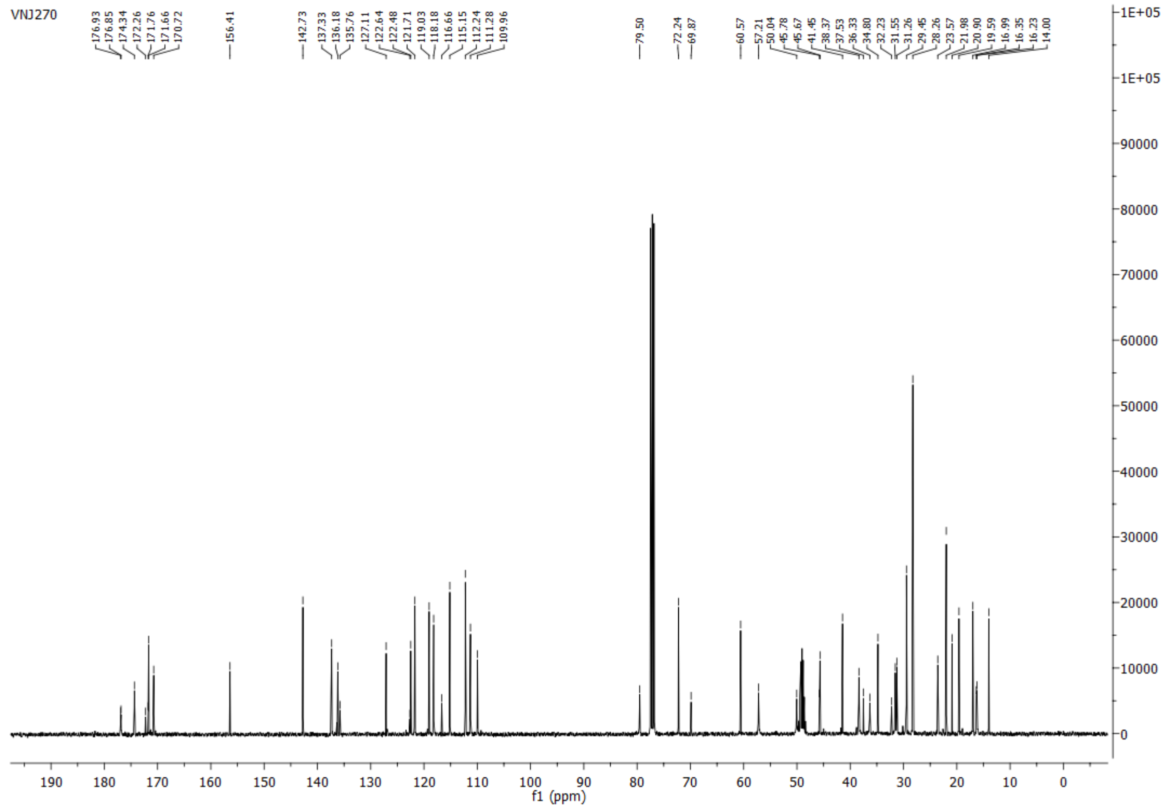

**$^{1}H$ spectrum of compound 26: 400 MHz, chloroform-*d*:methanol-$d_4$ = 8:2, 297 K**

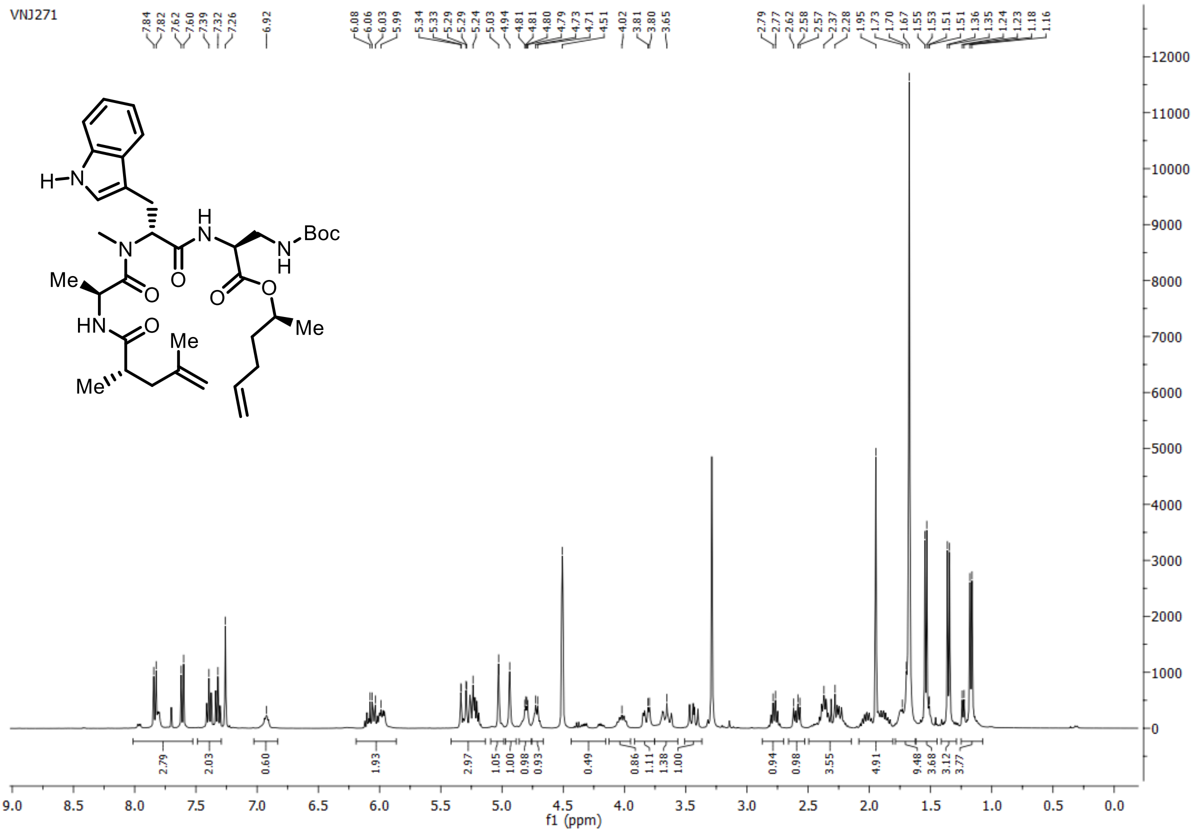

**$^{13}C\{^{1}H\}$ spectrum of compound 26: 101 MHz, chloroform-*d*:methanol-$d_4$ = 8:2, 297 K**

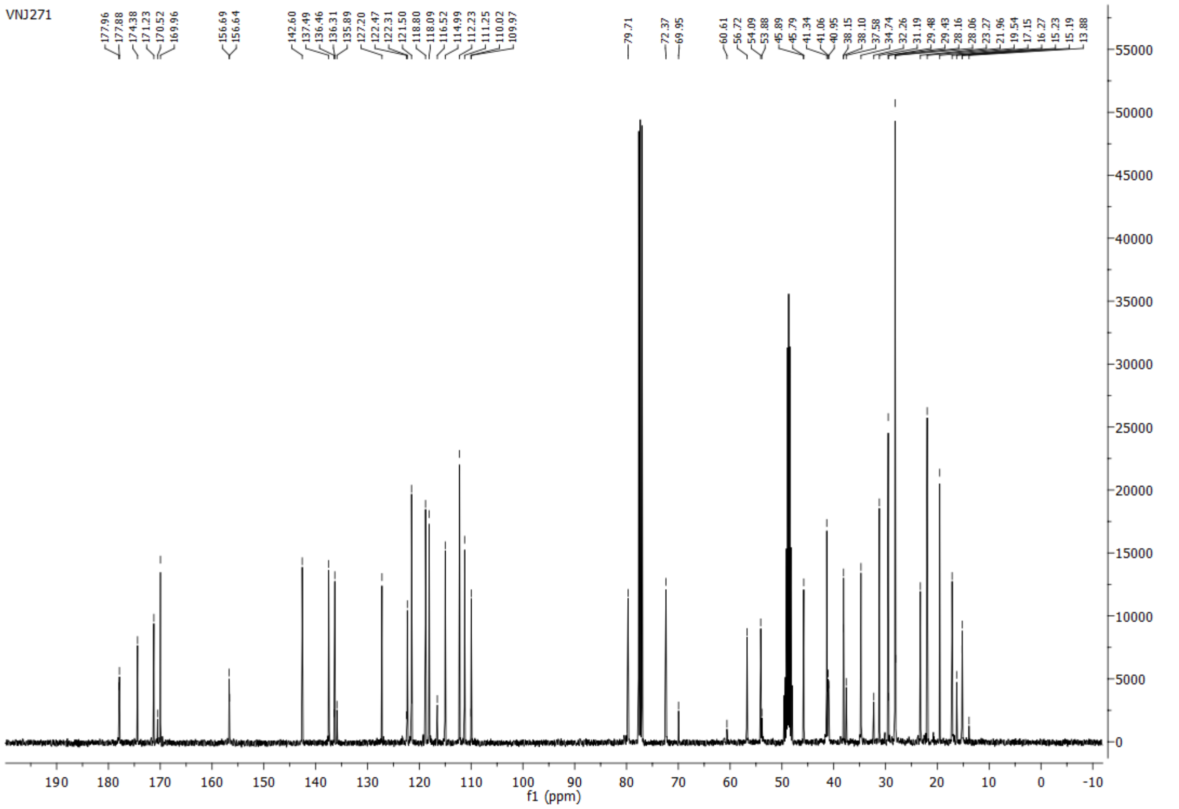

**$^{1}H$ spectrum of compound 7: 400 MHz, chloroform-*d*:methanol-*d₄* = 9:1, 297 K**

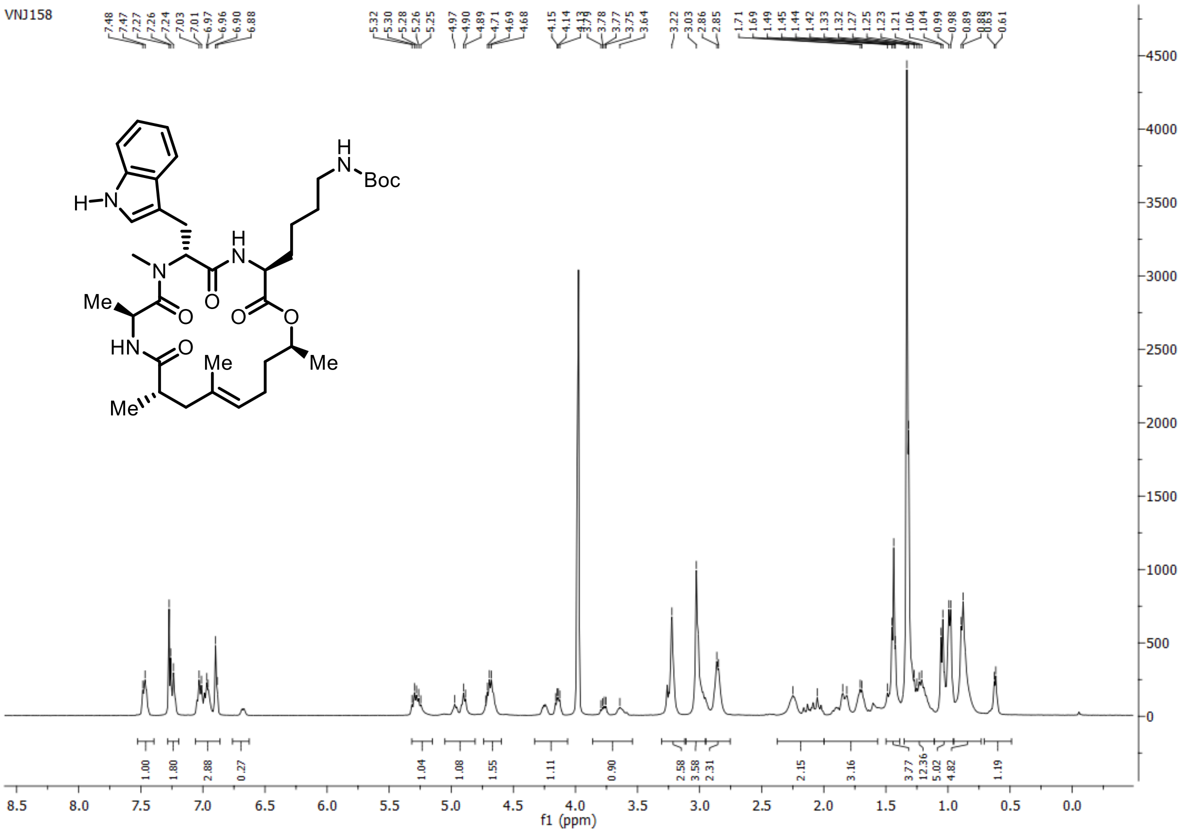

**$^{13}C\{^{1}H\}$ spectrum of compound 7: 101 MHz, chloroform-*d*:methanol-*d₄* = 9:1, 297 K**

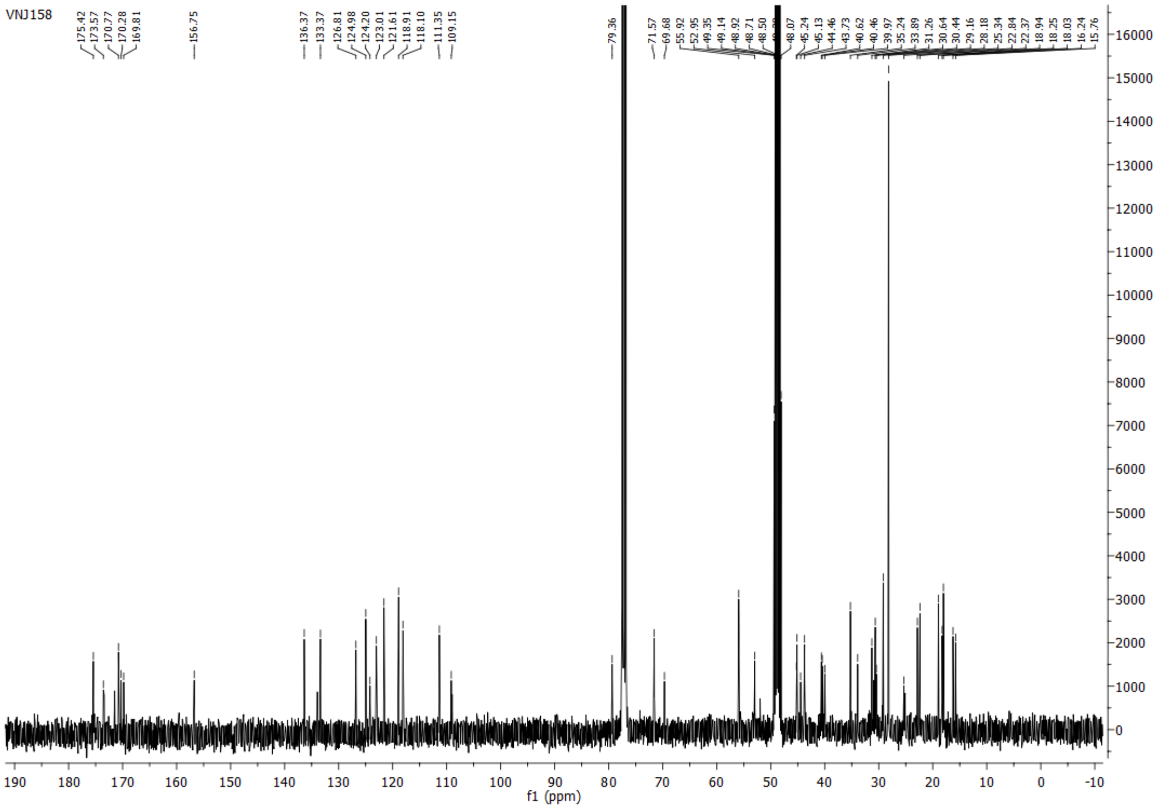

**$^{1}$H spectrum of compound 6: 250 MHz, chloroform-*d*, 297 K**

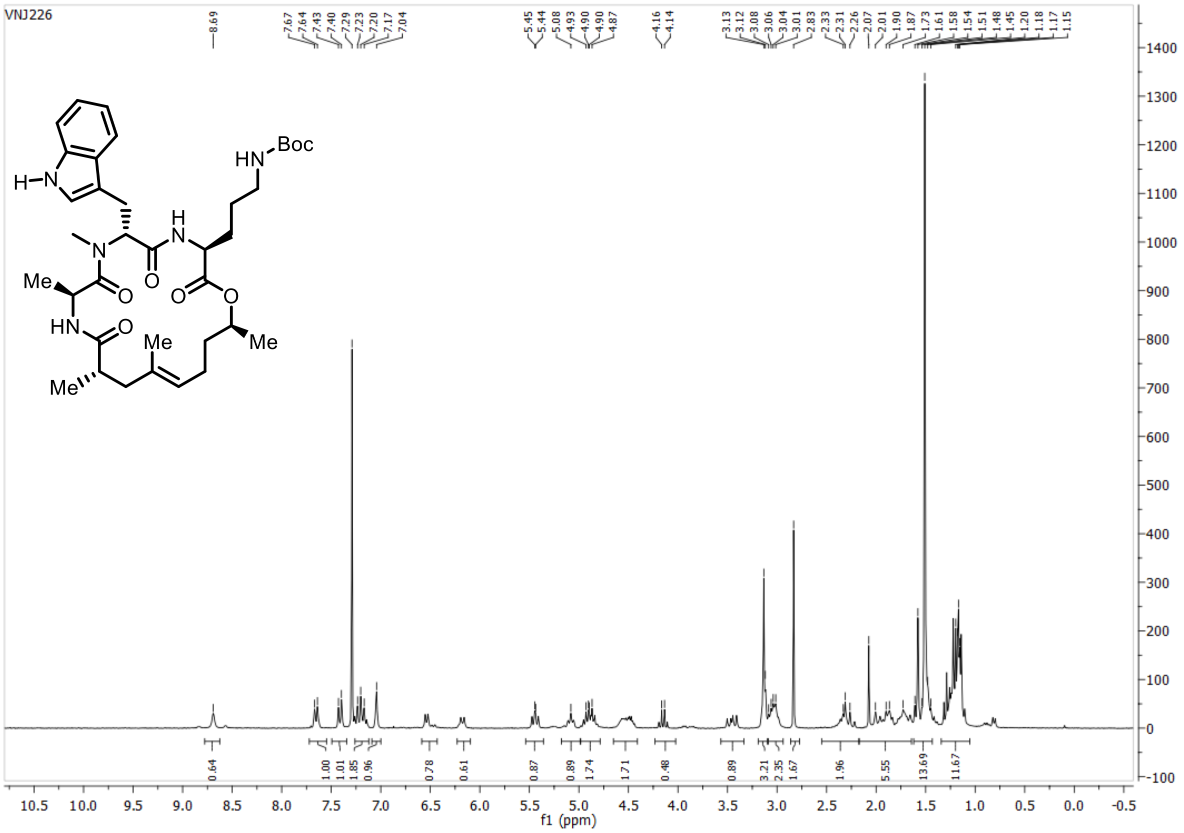

**$^{13}$C{$^{1}$H} spectrum of compound 6: 63 MHz, chloroform-*d*, 297 K**

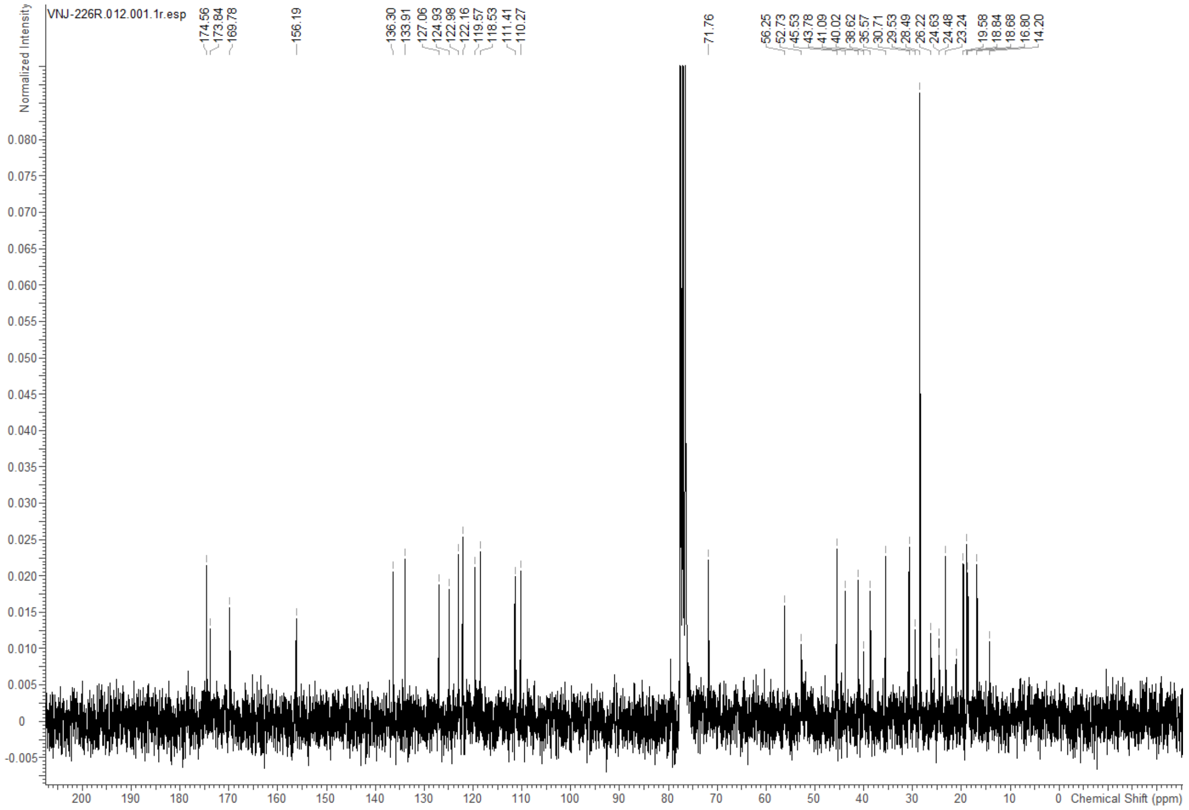

**[1]H spectrum of compound 5: 400 MHz, chloroform-*d*:methanol-$d_4$ = 9:1, 297 K**

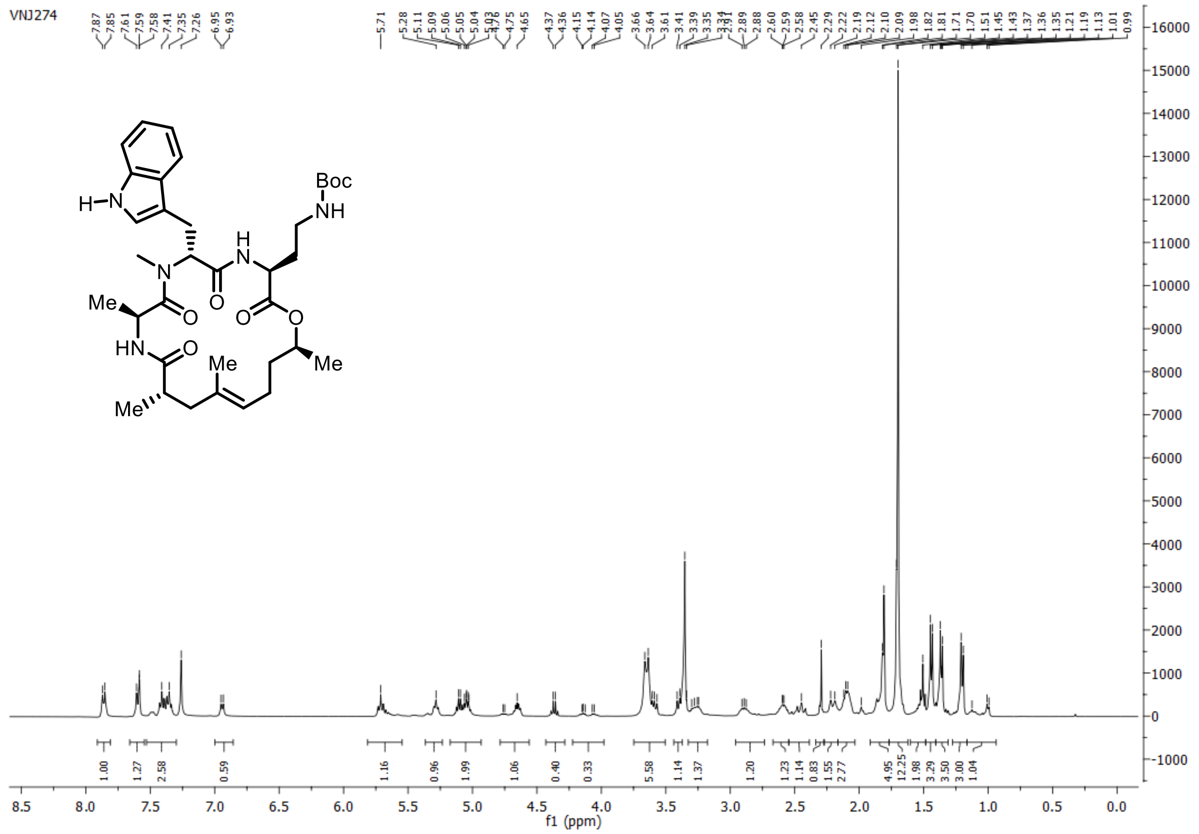

**[13]C{[1]H} spectrum of compound 5: 101 MHz, chloroform-*d*:methanol-$d_4$ = 9:1, 297 K**

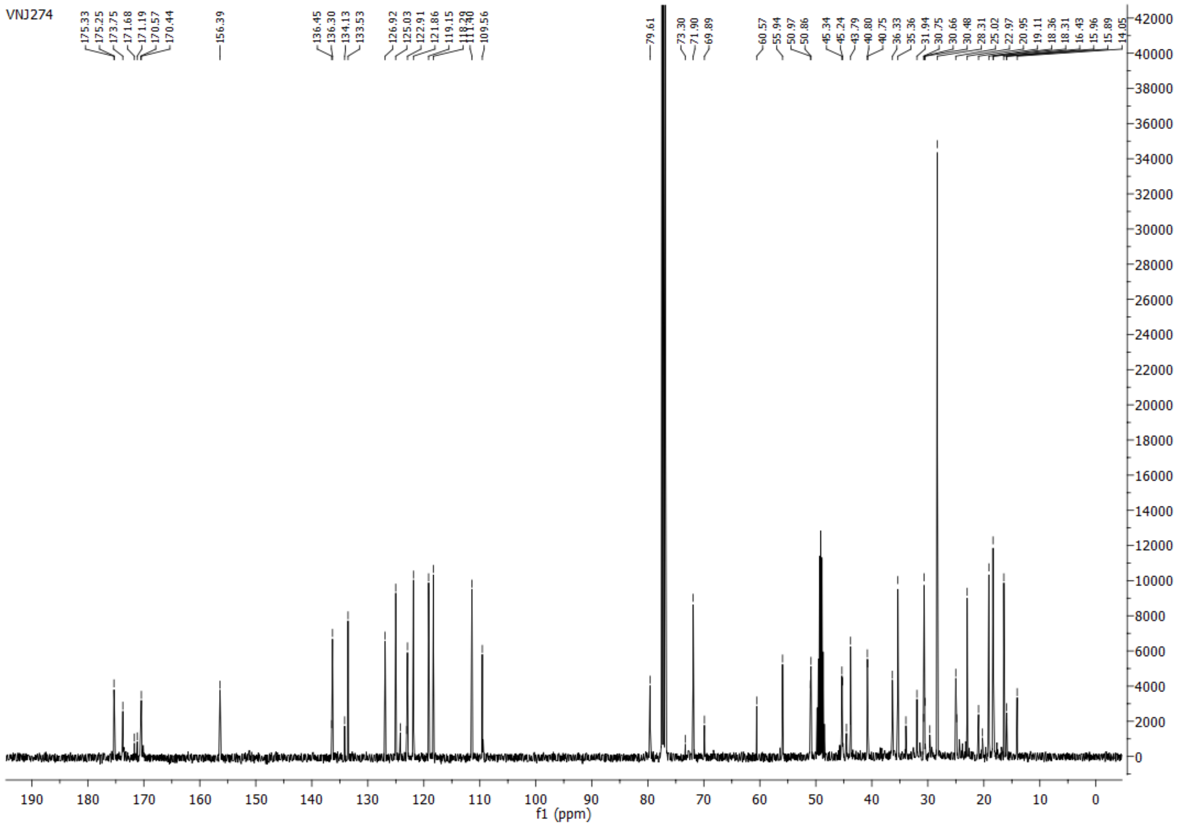

#### HSQC-DEPT spectrum of compound 5:

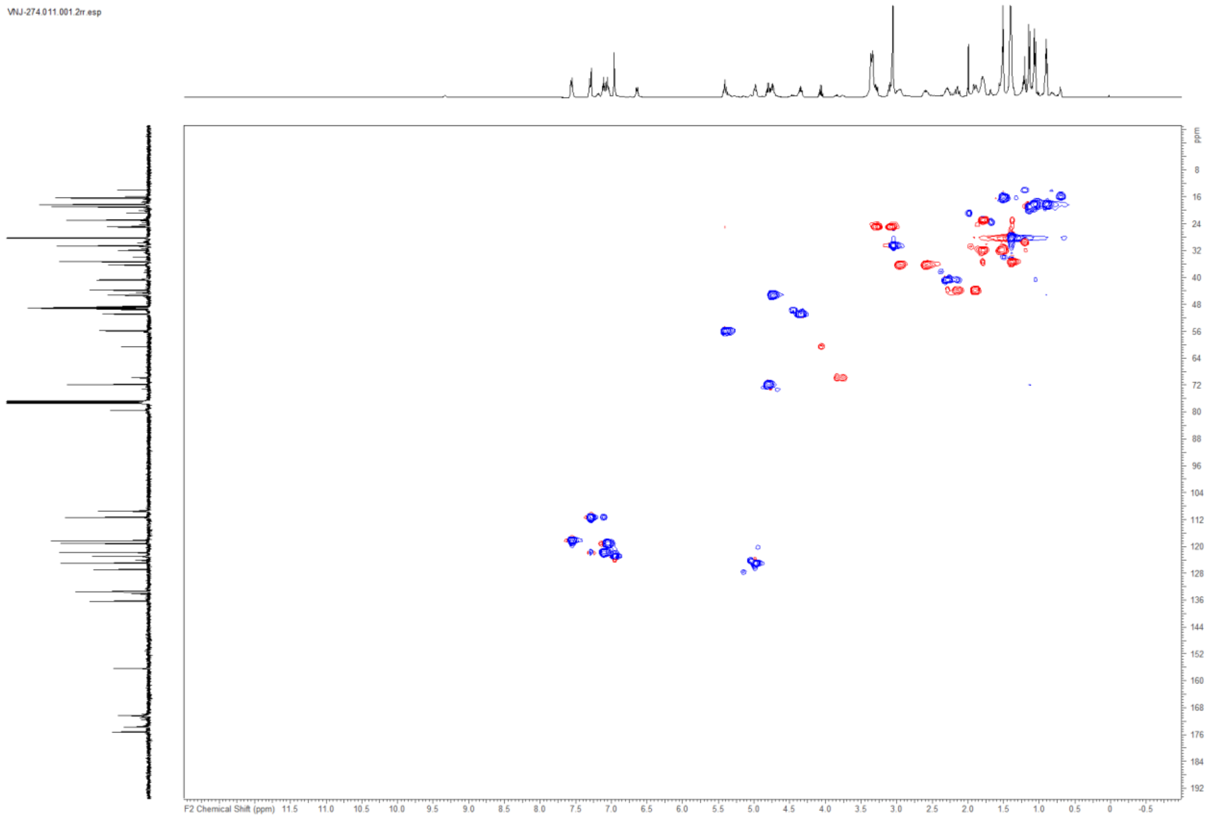

#### COSY-DEPT spectrum of compound 5:

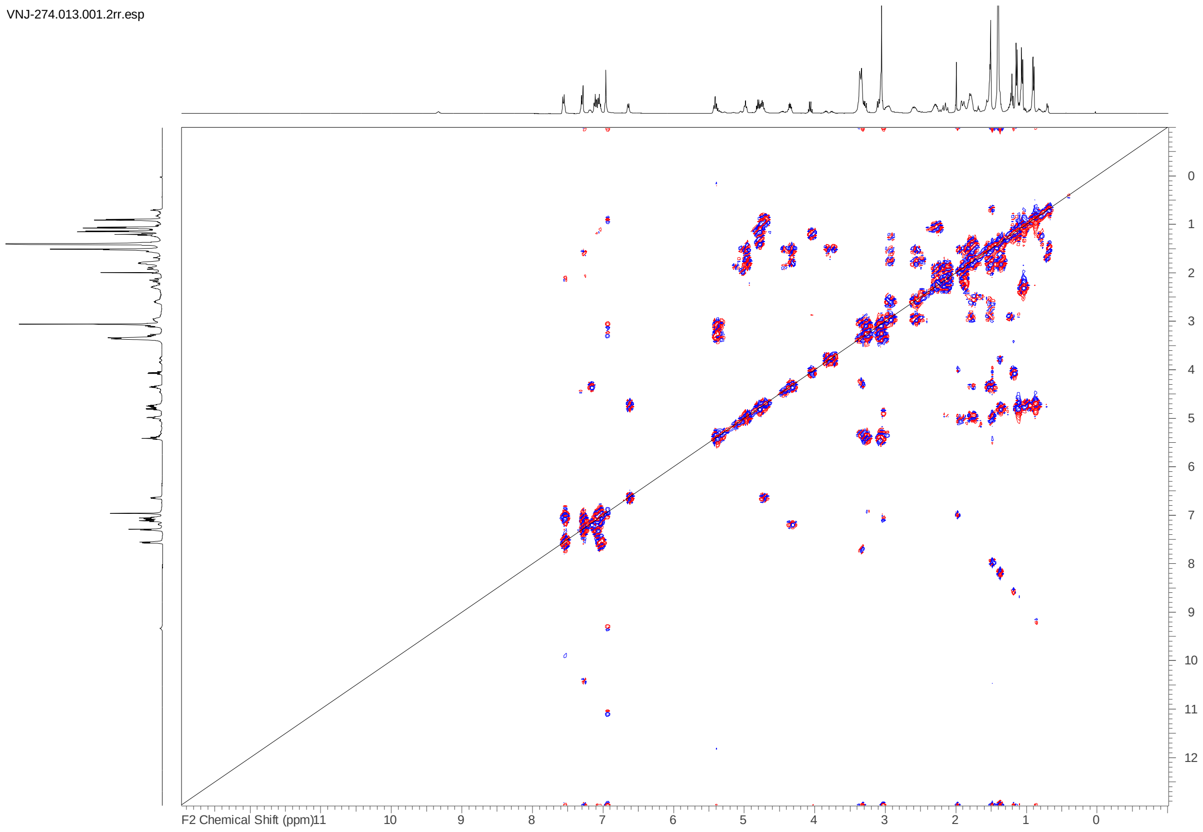

**$^{1}H$ spectrum of compound 4: 500 MHz, chloroform-*d*:methanol-*d*$_4$ = 9:1, 297 K**

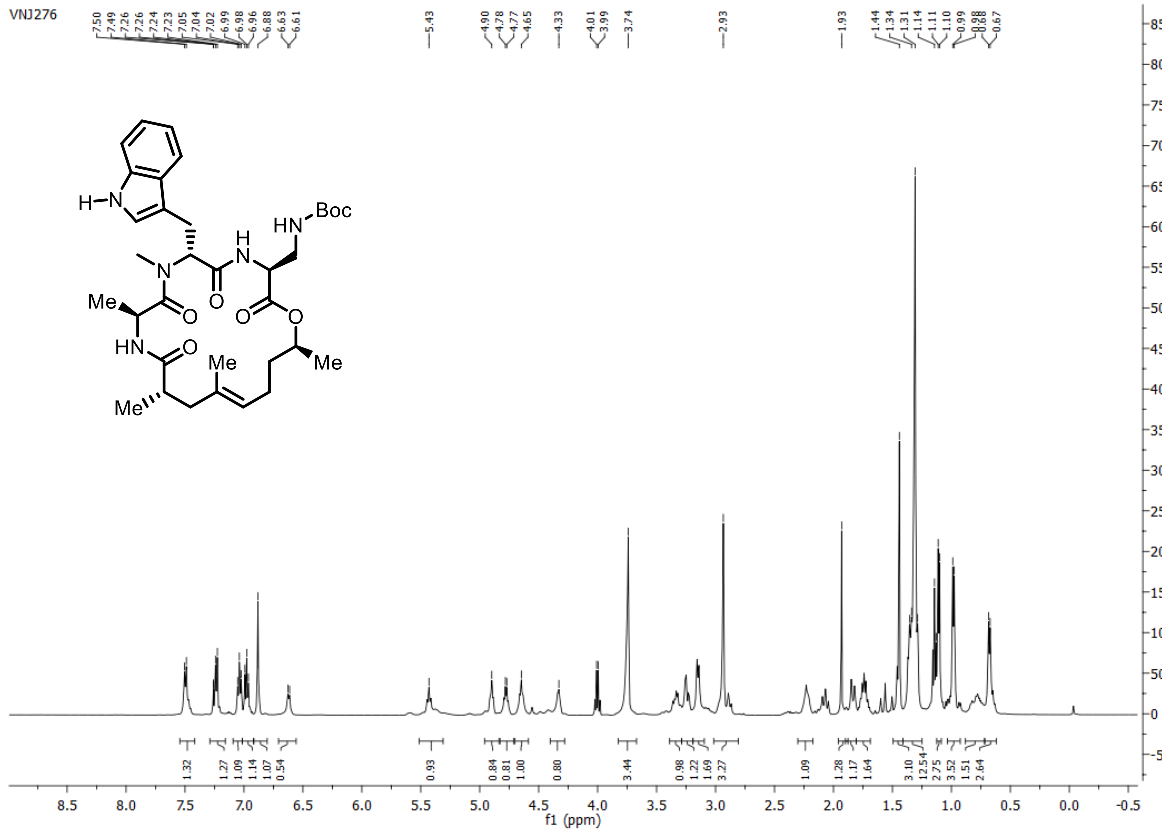

**$^{13}C\{^{1}H\}$ spectrum of compound 4: 125 MHz, chloroform-*d*:methanol-*d*$_4$ = 9:1, 297 K**

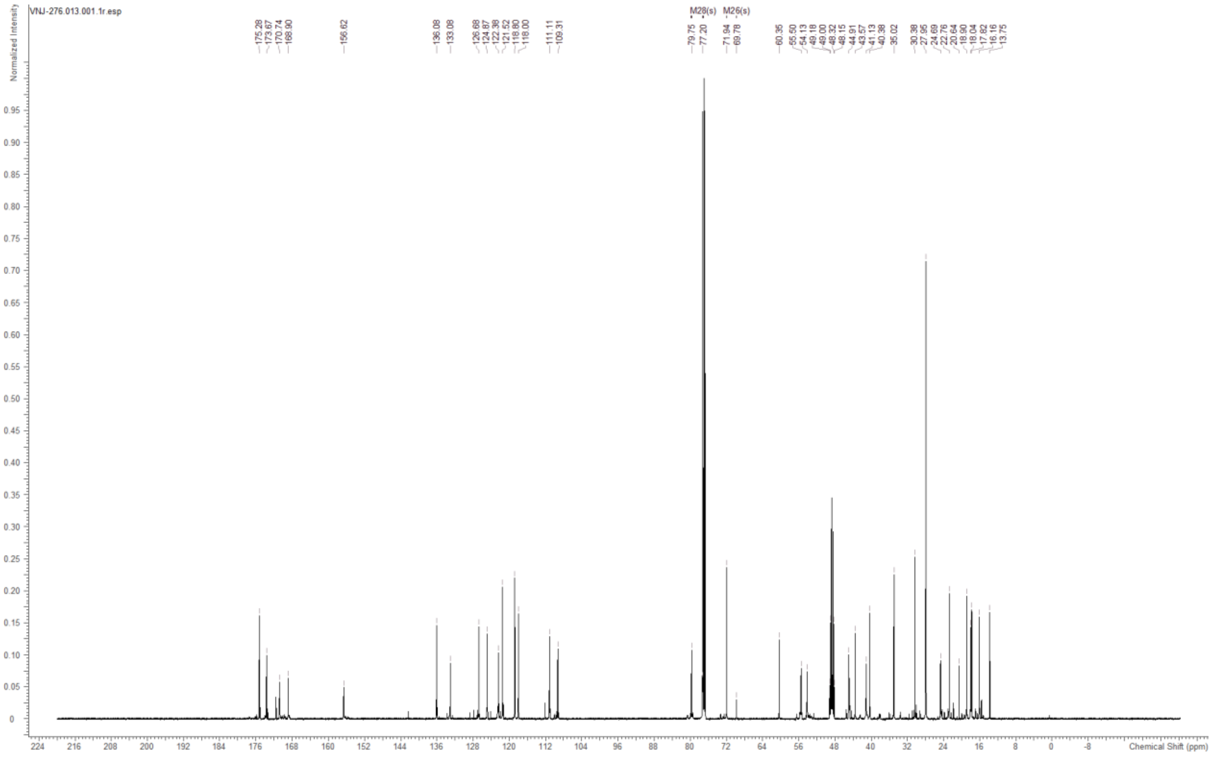

#### HSQC-DEPT spectrum of compound 4:

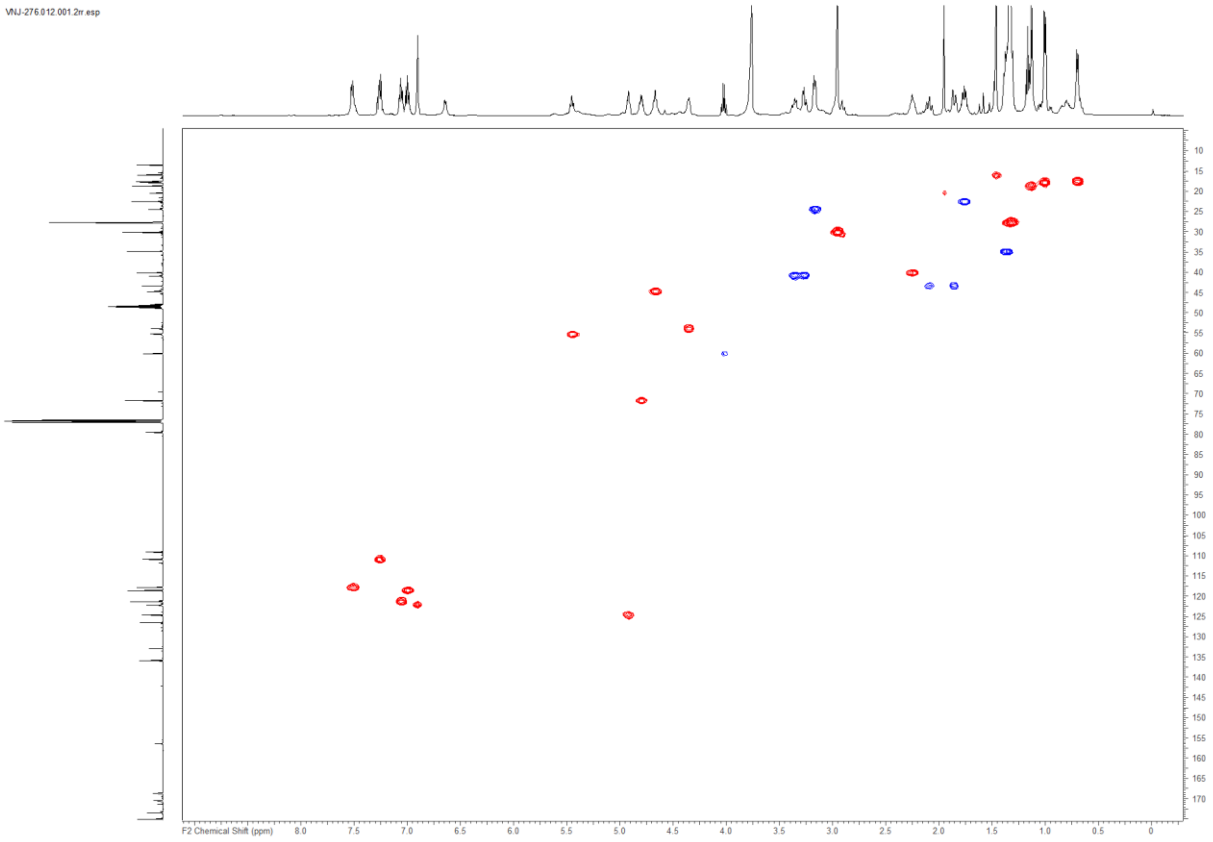

#### COSY spectrum of compound 4:

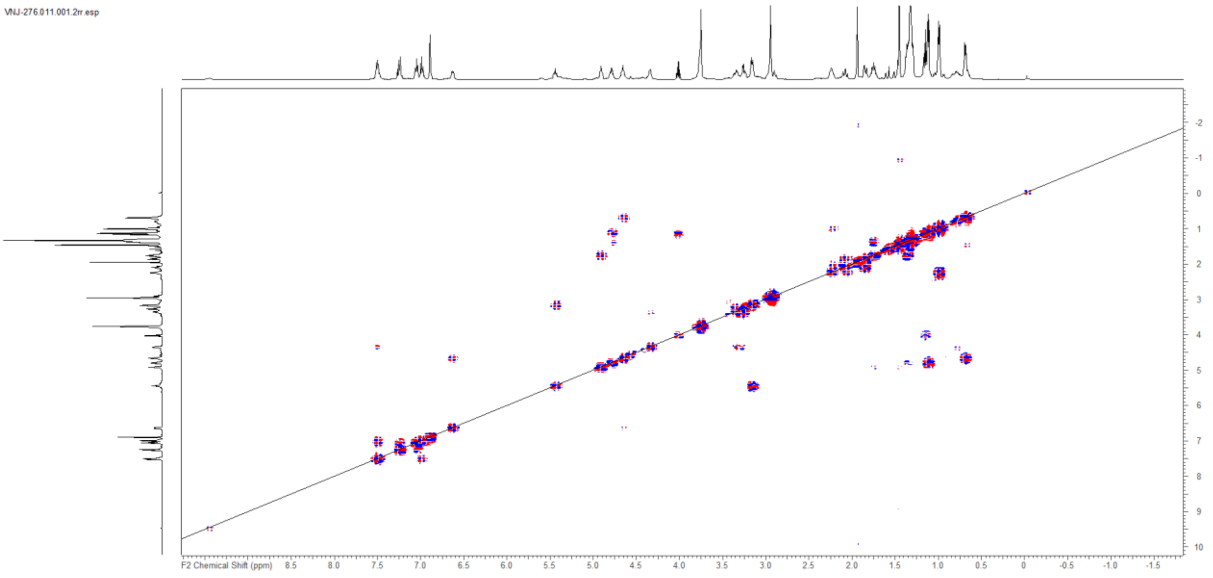

**$^{1}$H spectrum of compound 34: 400 MHz, DMSO-$d_6$, 297 K**

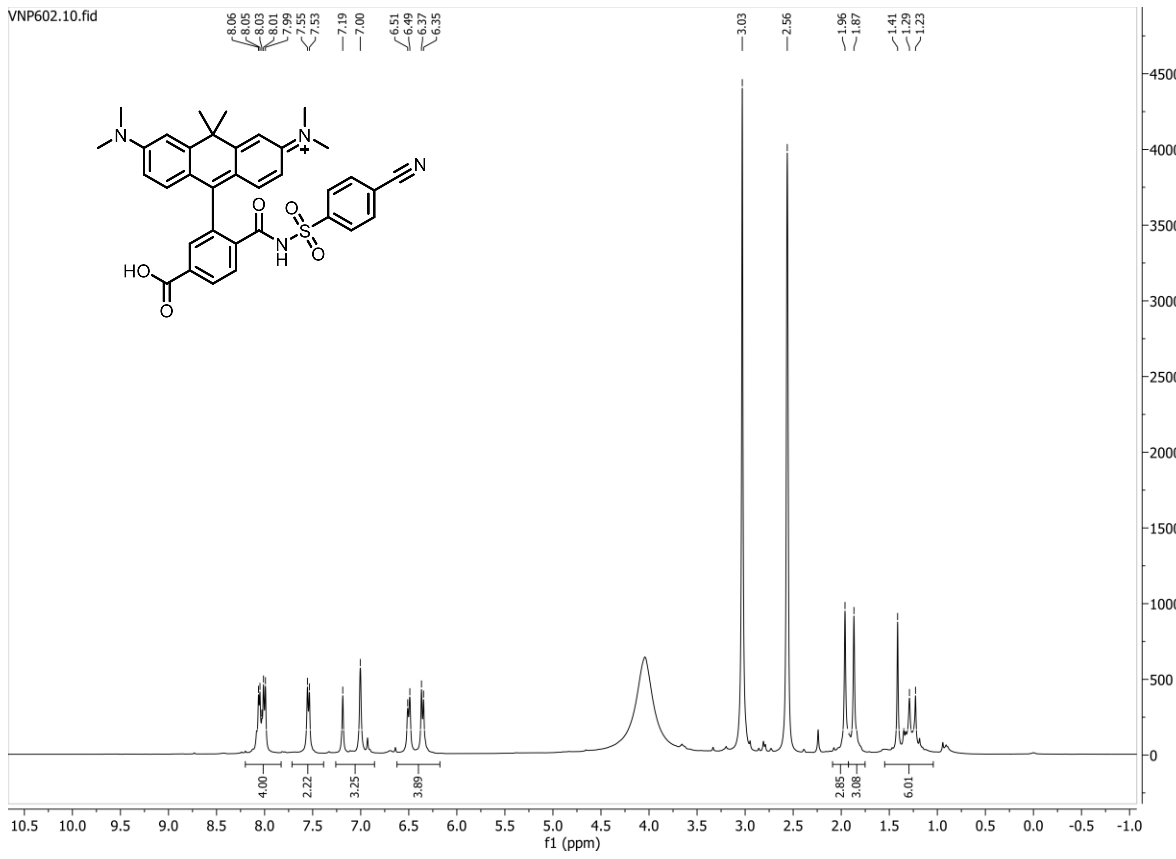

**$^{1}H$ spectrum of compound 8: 500 MHz, DMSO-$d_6$, 297 K**

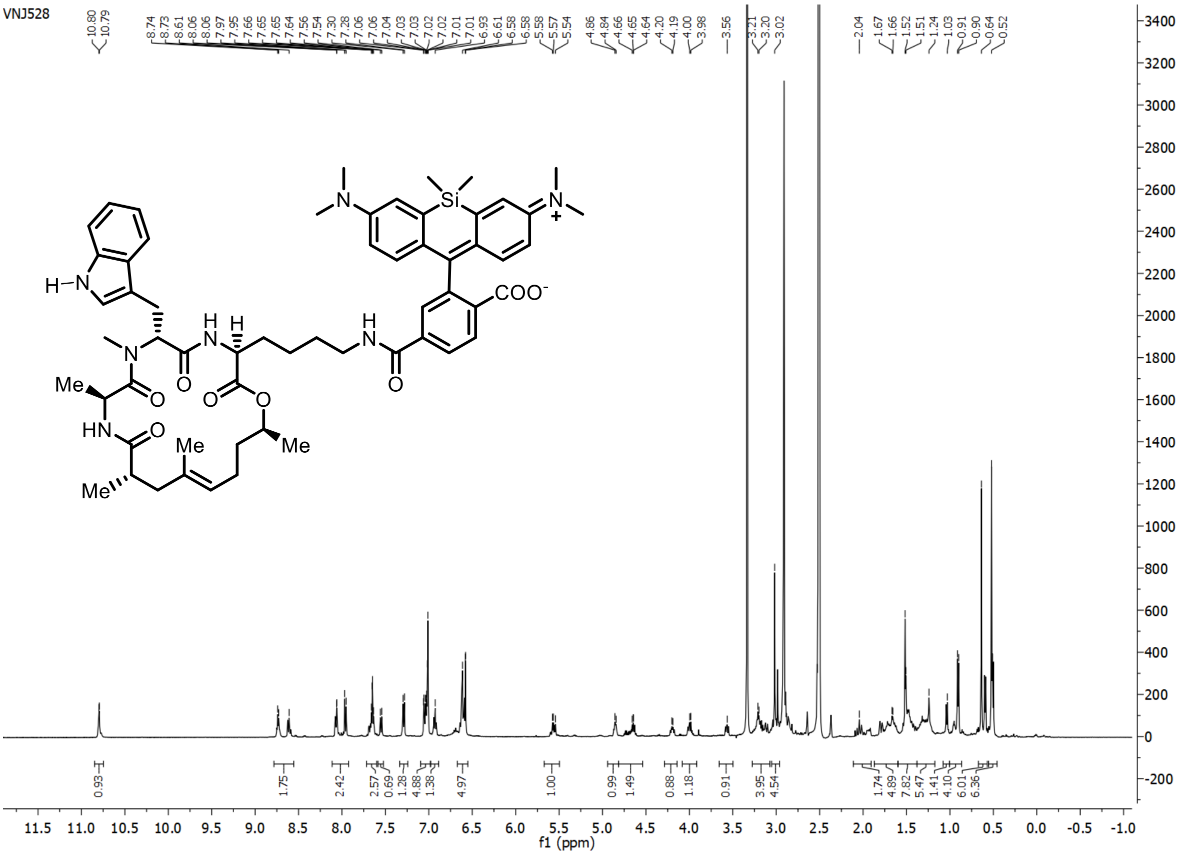

**$^{13}C\{^{1}H\}$ spectrum of compound 8: 125 MHz, DMSO-$d_6$, 297 K**

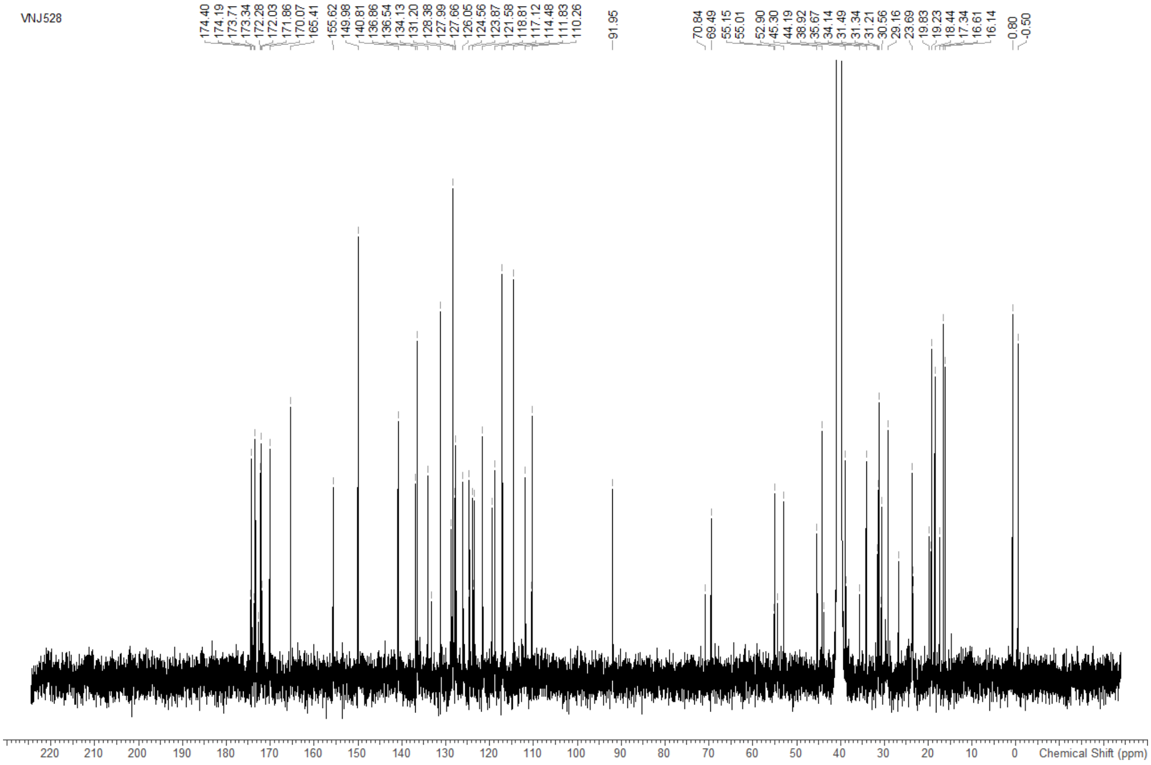
